# RNAprecis: Prediction of full-detail RNA conformation from the experimentally best-observed sparse parameters

**DOI:** 10.1101/2025.02.06.636803

**Authors:** Henrik Wiechers, Christopher J. Williams, Benjamin Eltzner, Franziska Hoppe, Michael G. Prisant, Vincent B. Chen, Ezra Miller, Kanti V. Mardia, Jane S. Richardson, Stephan F. Huckemann

## Abstract

We address the problem of predicting high-detail RNA structure geometry from the information available in low-detail experimental maps. Here, low-detail refers to resolutions ≈ 2.5-3.5Å, where the location of the phosphate groups and the glycosidic bonds can be determined from experimental maps but all other backbone atom positions cannot. In contrast, higher-resolution maps allow high-detail determinations of all backbone atomic positions. To this end, we first create a gold standard dataset of highly curated, experimentally supported RNA suites. Second, we develop and employ a modified version of the previously devised algorithm MINT-AGE to learn clusters that are in high correspondence with the gold standard’s conformational classes of suites based on 3D RNA structure. Since some of the gold standard classes are of very small size, a new modified version of MINT-AGE is able to also identify very small clusters. Third, we create a new conformer prediction algorithm, RNAprecis, which assigns low-detail structures to newly designed 3D shape coordinates. Our improvements include: (i) learned classes augmented to cover also very low sample sizes and (ii) replacing distances from clusters by Bayesian posterior probabilities. On test data containing suites modeled as conformational outliers, RNAprecis shows good results suggesting that our learning method generalizes well. In particular, we show that the modified MINT-AGE clustering can more finely delineate between previously unseen suite conformer separations. For example, the **0a** conformer has been separated into two clusters seen in different structural contexts. Such new distinctions can have implications for biochemical interpretation of RNA structure.

## 1 Introduction

In this work, we develop a new system, RNAprecis, for analysis of RNA backbone structure. To do this, we produce a quality-curated training set for automated use in analyzing the increasingly available number of RNA structures and develop a modified version of MINT-AGE. As a multidisciplinary collaboration between structural biochemistry and mathematical statistics, we adopt sophisticated treatments from non-Euclidean statistics, and we will cover a range of background material below.

### 1.1 Prior work and background in structural biology

A large fraction of RNA molecules form well ordered tertiary structures that have specific binding and catalytic functions. Achieving an accurate model of the RNA backbone is important for understanding RNA structure and function, because in RNA many of the significant functional contacts are with the backbone, as in the peptide transfer reaction of protein synthesis in ribosomes [1].

Correctly modeling RNA backbone conformations in X-ray crystallography and cryoEM experimental 3D structures is a long-standing challenge in structural biology. The RNA backbone has many degrees of freedom: 7 dihedral angles, plus the ribose sugar pucker. At “atomic” resolutions better than about 1.5Å, the positions of all individual atoms in the atomic model are experimentally determined, as by the electron density map depicted in Figure 1A. At resolutions worse than 2-2.5Å, the map for the backbone gradually becomes more featureless and ambiguous. The model’s fit to the map becomes less able to distinguish correct from incorrect local conformations, compounding the challenge presented by RNA’s many degrees of freedom.

**Figure 1:**
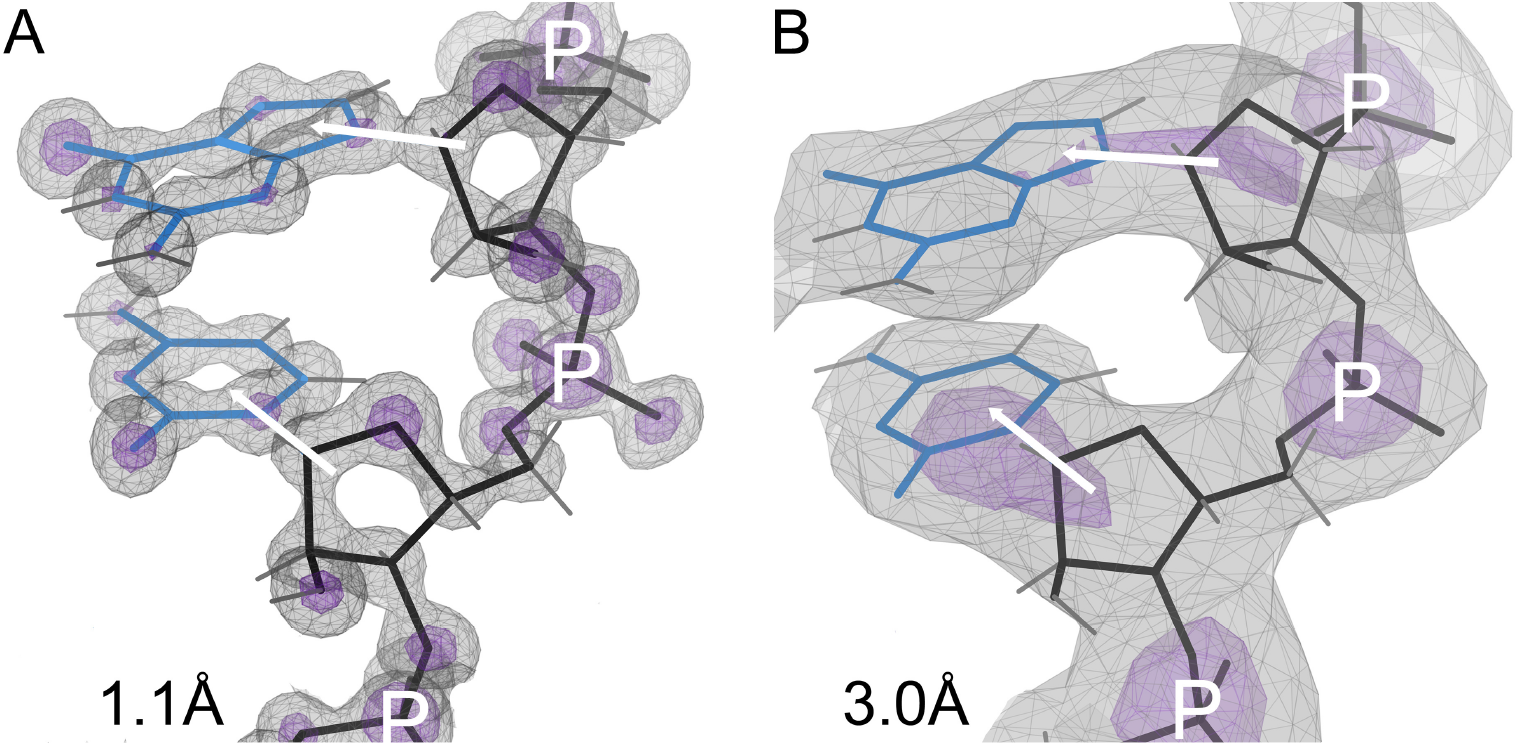
The positions of individual atoms in the derived atomic model are experimentally determined by the electron density when at high resolution (Panel A; 1.1Å), but not at low resolution (Panel B; 3Å). However, the phosphate positions and glycosidic bond directions can be seen directly at low resolution, and the pucker can be determined from them. Electron density contours are at 1.2*σ* (gray) and 3*σ* (purple) where *σ* is the standard deviation of the corresponding electron density map. Models have backbone in black, bases in blue. Phosphates are marked with P, glycosidic bond directions with a white arrow between sugar and base.

As shown in Figure 1B, at resolutions of 3-3.5Å, the phosphate positions and glycosidic bond directions can still be observed directly (and they may be the only features that can be reliably observed). The phosphorus is a much heavier atom than the rest, with a strong map signal, and is well located across this range of resolutions because it is the center of the symmetric PO_4_ group (the purple contours in Figure 1). The nucleobase is large, planar, and often held in place by base-pairing, and it usually produces a recognizable blob in the map, while the sugar forms a 3-way blob where the base and backbone join.

In contrast, the pucker state of the 5-membered ribose ring is hidden inside a large featureless blob at resolutions of 3-3.5Å. In RNA the pucker is quite cleanly bimodal (Figure 2A), as either C3^*′*^-endo (most common, and in A-form) or C2^*′*^-endo (frequent in irregular or non-helical regions). Other puckers are extremely rare in RNA. (DNA, in contrast, has many deoxyribose puckers.) The ribose pucker state is a dominant influence on RNA conformation, because a pucker change greatly alters the relative directions of the three large ring substituents: the base, the preceding backbone, and the following backbone (Figure 1B).

**Figure 2:**
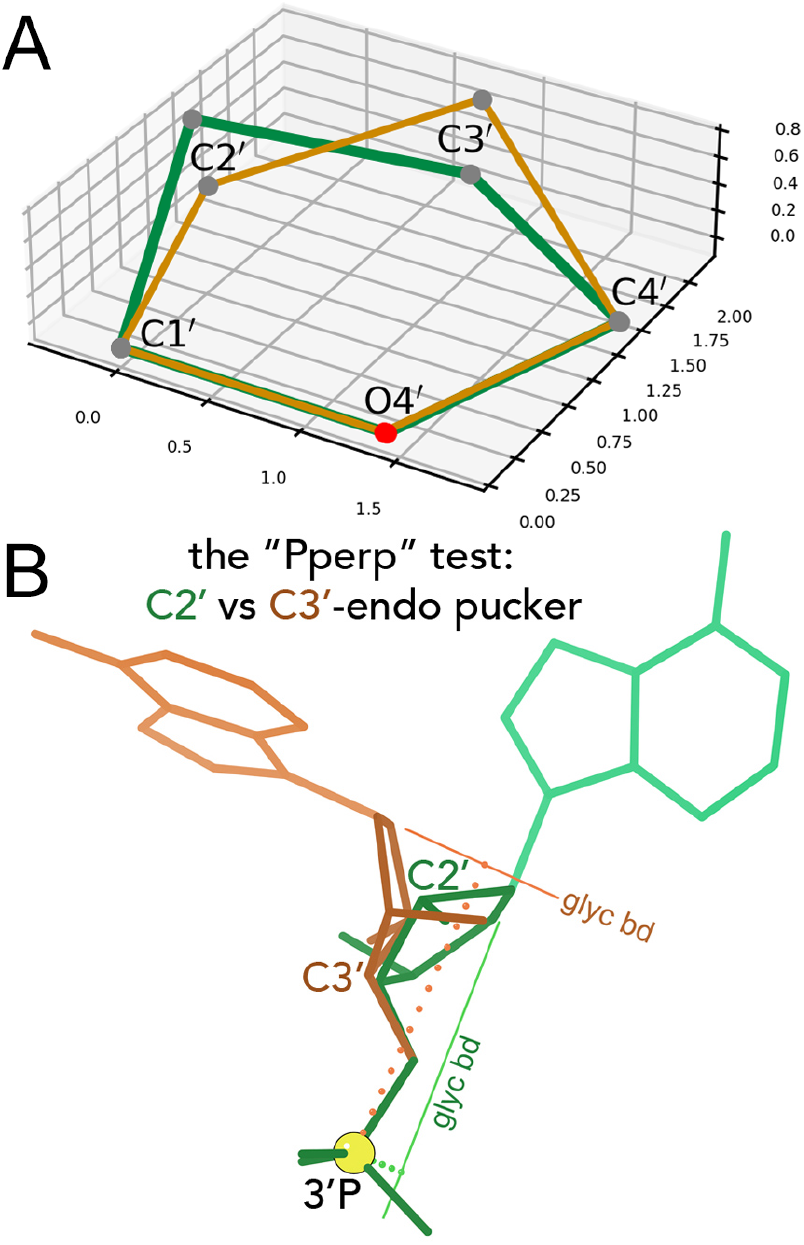
A: The two RNA ribose sugar puckers. In a C3^*′*^ endo pucker (brown), the C3^*′*^ atom is above (in positive orientation) the oriented plane spanned by C4^*′*^-O4^*′*^ and C1^*′*^-O4^*′*^. In a C2^*′*^ endo pucker (green), the C2^*′*^ atom is above the plane. Typically, all atoms but the puckered one are (nearly) co-planar. B: Contrasted cases, superimposed at their phosphate ends, to illustrate the Pperp criterion for distinguishing between C3^*′*^-endo (brown) and C2^*′*^-endo (green) sugar puckers in RNA. The atom puckered out of plane in each ring is labeled, and the vector extension of each glycosidic bond is drawn and labeled. The next P atom in sequence is marked by a large yellow ball. Dotted lines show the perpendiculars dropped from the P to each of the glycosidic bond vectors: long (and brown) for the C3^*′*^-endo puckered case; short (and green) for the C2^*′*^-endo case.

Almost 20 years ago [2], it was discovered that the ribose sugar pucker can be successfully and simply deduced from the more reliable observations of phosphorus position and glycosidic bond direction, even when the pucker itself has been modeled incorrectly. As shown in Figure 2B, the length of a perpendicular dropped from the next-in-sequence (3^*′*^) phosphorus atom to the extended vector direction of the glycosidic bond that connects base to ribose is *<* 2.9Å for C2^*′*^-endo and ≥ 2.9Å for C3^*′*^-endo. This process for identifying pucker was incorporated into MolProbity validation [2, 3], and the measure was named as the “Pperp” (*P*hosphate *perp*endicular) criterion [4]. Correcting a sugar pucker based on this deduction consistently corrects clashes and bond length and angle outliers within its suite, removes most difference density peaks, and in the aggregate lowers the main residual R and the residual R_free_ calculated with omitted data [5, 6, 4]. It works well with more complex systems such as ERRASER [7, 8], and it has been successfully in use within Phenix refinement as well as validation for over 15 years [9, 10], during which time Phenix has been disproportionately often used for RNA structures. There are other errors that cause outliers in RNA, but sugar pucker is a major one.

These map features and challenges generally hold across both X-ray crystallography’s electron density maps and single-particle cryo Electron Microscopy’s (cryoEM’s) Coulomb potential maps, the two predominant experimental methods for solving 3D structures of RNA. Sample preparation, data collection, and data processing are completely different, but both methods produce quite similar-looking 3D maps. CryoEM is sensitive to charge as well as mass, so for nucleic acids the positive bases are somewhat stronger and the negative phosphates somewhat weaker, but not to a degree that affects the Pperp criterion. In most experimental RNA structures, the overall resolution applies to a large majority of the residues or suites; however, local or effective resolution can vary. As local mobility or uncertainty increases, the map becomes less distinct, then patchier, and eventually disappears into noise. In such regions, the assumption of fairly robust PO_4_ position and glycosidic bond direction becomes inapplicable.

The overall success of the Pperp pucker analysis made us hope that other combinations of parameters, taken only from the best-observed features, might enable deduction of full-detail RNA backbone conformation from low-detail representations. We chose the name RNAprecis for this desired system, to indicate a concise description that captures the important aspects. (Precis: *pray*^*′*^*-see*, in writing composition, a short formal summary of a much longer document.)

Figure 3 diagrams the main features of full-detail local RNA 3D structure that we describe and analyze in this work. In Panel A the atomic model, with atoms named, covers two phosphate-to-phosphate nucleotide residues and one central sugar-to-sugar segment, known as a *suite*. For describing RNA backbone conformations, this suite is a more useful division than the chemical residue from P to P. The suite relates adjacent bases, including such important interactions as base stacking, and its atoms have many more contacts with one another than inside a residue [11]. A suite includes atoms from two adjacent, bonded residues. By convention, the suite is named by the residue number (i) of its second member residue—the same residue number as the P atom contained within the suite.

**Figure 3:**
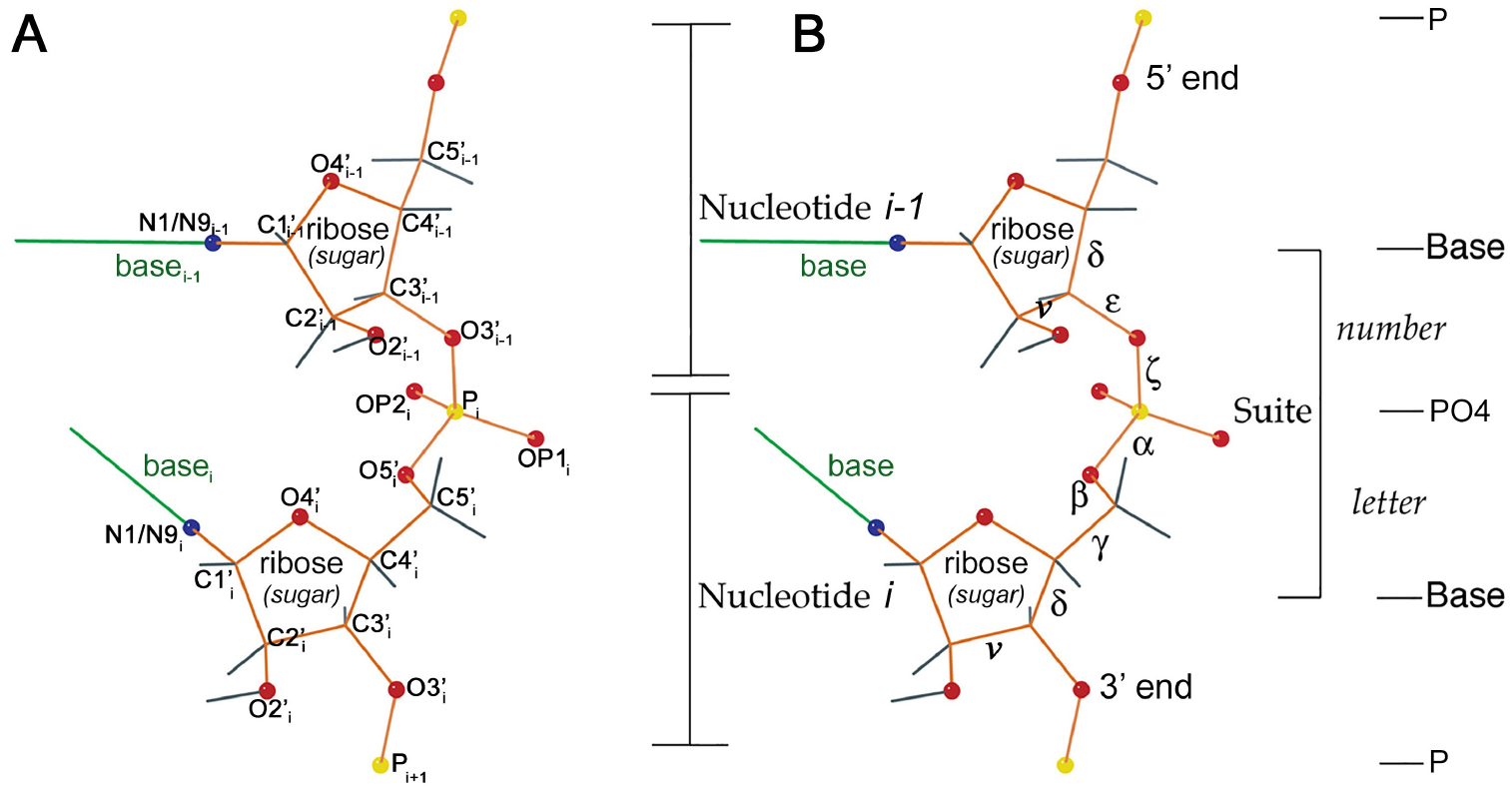
Atomic structure and nomenclature for the two residues and the central sugar-to-sugar suite of an RNA dinucleotide. The bases are simplified to single green lines. Backbone atoms other than carbon are marked with balls, red for O, blue for N, yellow for P, and with gray bonds out to H. Panel A: Names of heavy atoms used in describing suites. Panel B: The 7 dihedral angles of the suite are labeled in order as *δ, ϵ, ζ, α, β, γ, δ*. Additionally we show the dihedral angle *ν* measuring the sugar pucker (see Fig. 4). Number and letter refer to suite conformer names as e.g. in Section 2.2 and Table S4 in the appendix.

Viewed from the suite perspective, the RNA backbone has the attractive property of rotamericity [11]. That means bond geometry and steric contacts constrain the backbone to lie in generally distinct conformational clusters. Figure 3B names the backbone dihedral angles used by the RNA Society’s multi-lab RNA Ontology Consortium to define consensus suite conformers [12]. The number and letter labels denote the parts of those conformer names, such as **1a, 5z, 6n**, etc. At the time of that work, naive clustering algorithms failed at satisfactorily identifying and separating suite conformers, at least partly because 2/3 of the suites in the single A-form helical conformation overshadowed the rarer conformations. Therefore the 2008 conformers were first separated automatically, and then further conformer class boundaries were assigned by hand in a 7D graphics system developed for that purpose [12]. Additionally we tested whether close clusters should be separated or not by whether they were distinct or interchangeable in their local structural motif or functional role. Suites with conformations that do not match any of the consensus suite conformers are considered outliers for validation purposes and given the **!!** designation (pronounced “bang-bang”).

The original suite conformers were identified based on the complete set of 7 backbone dihedrals. However, these dihedrals are only reliably modeled at resolutions where all their member atoms are clear in the experimental map. Just as the Pperp criterion predicts the correct sugar pucker from minimal model information, and even in the presence of modeling errors, RNAprecis seeks to predict, as accurately as possible, correct rotameric suite conformers from minimal model information.

### 1.2 Prior work and background in geometric statistics

Statistical analysis of geometric objects such as suite conformers requires firstly their mathematical representation. For instance, aggregating the positions **x**_*i*_ ∈ ℝ^3^ of their *k* atoms (*i* = 1, …, *k*) results in a configuration matrix

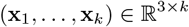

where each column is called a *landmark* vector.

Secondly, it is necessary to mathematically account for the fact that a translated and rotated suite conformer represents an object of the same shape. Thus the *shape* of a configuration matrix is the *configuration equivalence class* of all configuration matrices that can be reached from a single configuration by applying a single rotation and a single translation to all of the columns. Developing statistical methods for the analysis of such and related concepts of shapes has picked up momentum since the middle of the last century and a broad overview is given by [13]. The following two representations of shape are elaborated in [14]

As detailed above, a favorable representation of rotamer shape at high detail, avoiding configuration equivalence classes altogether, is given by a dihedral angle representation. Indeed, while individual bond lengths and angles between bonds at atomic centers are fairly fixed (with standard deviations on the order 10^−2^Å for lengths and ∼2 degrees for angles [15]), structural diversity is caused by rotation around bonds. These rotations, given by dihedral angles, can be measured in arc length on the unit circle,

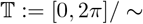

where “*/*∼ “ indicates formation of the quotient space (circle) from the interval by identifying the endpoint 0 with the other endpoint 2*π*. Then, the shape of one suite at high detail, determined by 7 dihedral angles, see Figures 3 and 4, is given by an element of the 7-dimensional torus

**Figure 4:**
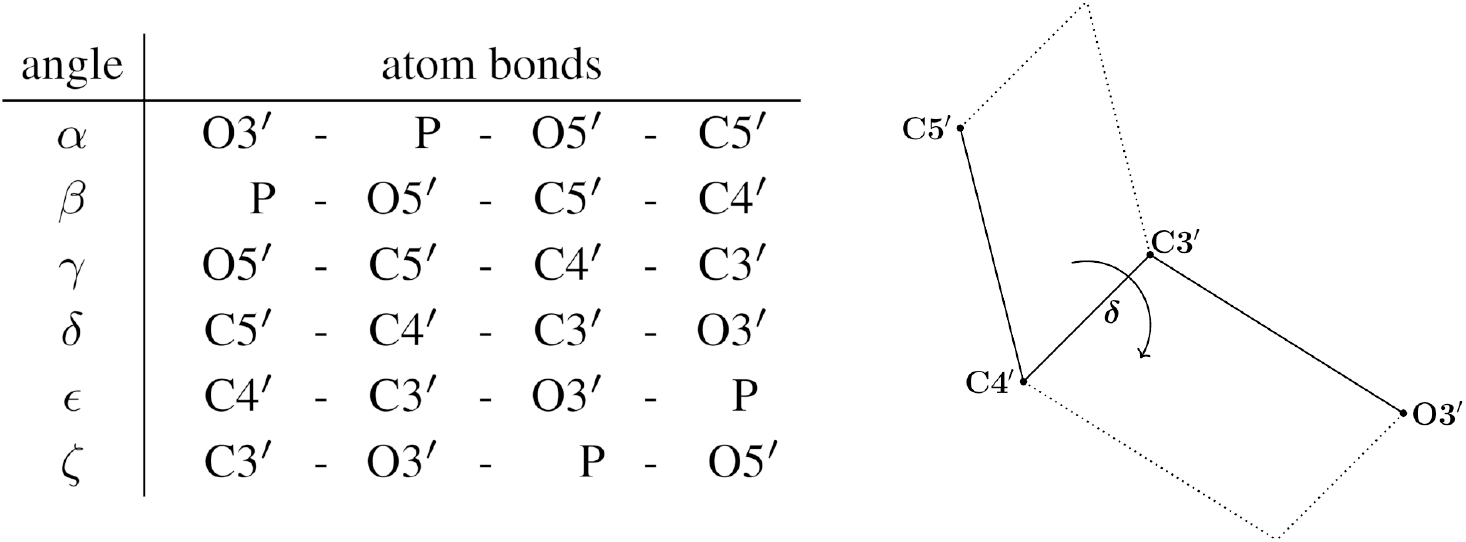
Left: the dihedral angle (first column), gives the rotation from the plane spanned by the last three atoms with respect to the plane spanned by the first three atoms, as depicted in the exemplary right panel (Figure and table adapted from [31]), see Figure 3 The first 7 measure backbone dihedral angles, distinguished from *ν* which measures the sugar pucker, see Figure 2A.

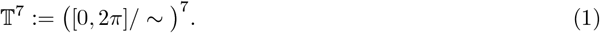

**MINT-AGE.** In order to statistically cluster torus data in an unsupervised way, in previous work we had developed MINT-AGE (Mode huntINg after Torus pca on Adaptive iterative cutting linkaGe trEes). Its building block, AGE, can be applied to data on any metric space. AGE first builds a linkage tree (e.g. average linkage), as introduced by [16] and then performs a specifically designed iterative adaptive clustering method inspired by [17, 18], identifying first the lowest density cluster, then iteratively higher density clusters. The method is detailed in [19] and our specific tuning parameters are listed in Table 1. The building block MINT is then applied to every (pre-)cluster found by AGE. Each cluster is subjected to *torus PCA* from [20]. Circumventing the fundamental difficulties of straightforward PCA approaches on a torus, here the 7-dimensional torus is mapped to a 7-dimensional stratified sphere (where the stratification mimics the topology of the torus) and an adaption of the *principal backward nested subsphere* algorithm (PNS) from [21] (a variant of PCA for spheres), using torus distances rather than spherical distances, is applied to the data, which is then projected to its main one-dimensional small circular component. These projected data are subjected to circular mode hunting (e.g. [22, 23]) where each statistically significant mode stands for a new subcluster. Originally, MINT used nonparametric circular mode hunting ([20] building on the linear version by [24]), where cluster boundaries had been placed at antimodes of suitably kernel smoothed data utilizing a similar approach to the WiZer (Wrapped Gaussian kernel smoothed derivative’s significant Zero crossings, from [23] building on [25]). In order to identify also very small subclusters, even of size 2, we use a parametric approach, namely a likelihood ratio test for the likelihoods of fitted unimodal and bimodal Gaussians, recently developed in [26]. This test can be performed iteratively to find several subclusters on a main one-dimensional small circular component.

**Table 1:** Selection of tuning parameters required for the AGE algorithm from [19] for the four different pucker-pair HD data sets: relative branching distance *q*, minimal cluster size *κ* (set to *κ* = 3 for all four data sets inorder to also identify clusters with very few elements) and the maximum outlier distance *d*_max_ (given as the percentage of elements that are in a cluster with less than *κ* elements).

| dataset | $q$ | $\kappa$ | $d_{\text{max}}$ |
| --- | --- | --- | --- |
| HDP33 | 0.09 | 3 | 1.1% |
| HDP32 | 0.05 | 3 | 0.34% |
| HDP23 | 0.07 | 3 | 1.4% |
| HDP22 | 0.05 | 3 | 0.7% |

**Multicentred coordinates for constrained shape.** At low detail, where variation of pseudo-bond angles and lengths contribute considerably to shape variability, we resort to a landmark-based shape representation. Once again, however, we avoid the abstract modeling by configuration equivalence classes and develop a *frame-based* method. Here, specific landmarks (i.e. specific prominent atom positions) are singled out and mapped to a standardized frame, thus uniquely determining rotation and translation of the other landmarks. Frame based methods have been successfully utilized as *Bookstein coordinates* [27], *Goodall-Mardia coordinates* [28] and also for *affine* and *projective shape* [29]. In all of these approaches, one landmark is subtracted from all others, thus becoming the origin. This role will be taken by the landmark describing the central PO_4_ group. For data at low detail the rotation will be uniquely determined by rotating the pseudo-bonds from *P* to each of the two C1^*′*^ atoms into a shared coordinate plane. Since data at low detail also contains the glycosidic bonds of nearly fixed lengths between each C1^*′*^ and its neighboring N1/N9, the location of the former will be subtracted from the latter, leading to *multicentred coordinates* for *constrained shape* as proposed in [14]. Fréchet means [30] of MINT-AGE clusters will be computed in multicentred coordinates and varying cluster breadth will be accounted for by a Gaussian likelihood in Bayesian inference.

## 2 Data and Methods

### 2.1 Data sets used

**Sugar pucker-pair sets.** Since the ribose puckers are bimodal, a good start on automatically identifying full-detail backbone conformers is to sort conformers into 4 categories by each suite’s two sugar puckers at each side. We call these the *pucker-pair* sets:

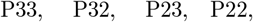

where C3^*′*^-endo and C2^*′*^-endo have been shortened to just 3 and 2. The P33 group is much larger than the others, because it includes the dominant A-form double-helical conformation (**1a**).

At high detail, the sugar pucker is determined directly from atoms in the ribose ring, using the *ν* dihedrals (see Figures 2 and 4), where *ν*_1_ refers to the first ribose in the suite and *ν* _2_ to the second. At low detail, the ribose ring atoms are not assumed to be reliable, and sugar pucker is instead determined using the Pperp criterion (see Figure 2B). Note that the P atom from the next sequential suite (with the index *i* + 1) is required for determining the second sugar’s Pperp distance and pucker conformation.

**High-detail gold standard.** The first training dataset, denoted by HD, comprises complete atomic positions of RNA suites obtained by high-resolution (*<* 2.0Å) X-ray crystallography and CryoEM. It is derived from a preliminary version of a carefully curated RNA residue dataset, RNA2023 [32]. The 3.150 release of BGSU RNA 3d Hub^1^ classified RNA chains of similar function and nucleic acid sequence into *sequence equivalence classes*. (Note that sequence equivalence classes are an entirely distinct concept from the configuration equivalence classes introduced and used only in Section 1.2; the overlap in terminology represents one of the challenges in interdisciplinary work.) Then the best chain is selected from each sequence equivalence class based on resolution, RSR, RSCC, Rfree, clashes, and completeness [33]. Chains from two high-resolution CryoEM ribosome structures, namely 8b0x (E. coli) [34] and 8a3d (human) [35], were added to this set of chains.

Since even high-resolution structures may contain local disorder or local modeling errors [36], the selected RNA chains were then filtered at the suite level. Suites with strong indicators of modeling errors, such as Pperp/pucker mismatches, steric clashes, and covalent bond geometry outliers have been removed from the dataset. Likewise, suites with insufficient support from experimental data, as measured by local 2mFo-dFc map values (or inclusion in cryoEM map envelope), local real-space correlation coefficients, or alternate conformations were removed. Additionally, suites involving a non-standard RNA base (i.e. other than A, U, C, or G), and suites identified as “ !!” suite outliers by `phenix.suitename` validation were removed.

This quality-filtered data set serves as the *gold standard*. Since puckers are also recorded, H D comprises the following four datasets:

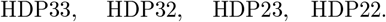

**Low-detail** datasets comprise only positions of the large atomic agglomerations, namely the positions of the central P of the phosphate groups and the ends of the glycosidic bonds, i.e. the positions of the C1^*′*^ and N1 or N9 atoms, respectively. For training purposes, from the *high-resolution* dataset a low-detail dataset denoted by LD has been obtained by taking the only corresponding 5 atom positions, yielding, four datasets featuring corresponding sugar puckers:

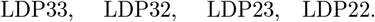

In Section A in the appendix, the PDB files of the high-resolution dataset are listed with the corresponding resolution in Table S1 and the HD and LD representations, with their pucker-pairs, are plotted in Figure S1 and Figure S2 in the appendix.

**Low-detail test data and related conformer classes as best-guess answers.** For testing how well our trained system can predict correct conformations, test sets were created. These test sets address two related questions. First, are the model features used by RNAprecis reliably preserved in typical RNA modeling errors? Second, given reliable parameters, can RNAprecis predict best guess solutions for RNA modeling errors? The first test set, named LDT, contains only conformational outlier suites from structures solved at lower resolution than the training set. The second data set stems from the 23S large ribosomal subunit described by the PDB identifier 8b0x chain a (lower case), which is a member of the final RNA2023 dataset but has no correspondence in the training data and can therefore be considered an out-of-sample test set. These datasets comprise, for each member suite, measurements of the same five atomic positions as in LD (the central P and the two neighboring C1^*′*^ and N1/N9), plus an additional sixth atom (P from the next residue following the suite). LDT is divided using the Pperp criterion into the following pucker-pair datasets:

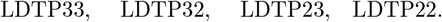

and correspondingly for the 8b0x dataset we have the pucker-pair datasets:

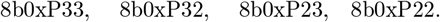

Suites for the test sets were selected to provide examples of modeling errors with probably known solutions from related structures. For each chain in the HD training set plus 8b0x chain a, related chains were identified based on the sequence equivalence classes from the 3.328 release of http://rna.bgsu.edu/rna3dhub/nrlist [33]. If a sequence equivalence class contained many chains, a representative chain was selected for the test set, based on that chain having an experimental resolution near the median resolution for that class and a count of **!!** outliers near the median count for the class. Such chains represent a “typical” degree of modeling challenge for their respective classes. A single equivalence class could contribute multiple chains to the test set if those chains had outliers at different sequence locations. Sequence equivalence classes of low sequence variety (such as poly-U) were excluded, as the relatedness of their member chains’ conformations is uncertain.

A sequence alignment was performed on the selected chains with their reference chain from HD. From the aligned chains, suites that were modeled (i.e. identified) as **! !** outlier conformations were selected as members of the LDT test set if the reference chain had a recognized, non-**!!** conformer modeled at the same sequence location. The reference chain’s conformer was taken as the best guess correct answer. These reference chain suites all passed the stringent residue-level filtering for the RNA2023 dataset.

In Section B in the appendix, the PDB files, the chain names, and the residue numbers of the elements that comprise the low-detail test datasets are listed with their associated resolutions. Table S2 lists the suites of the LDT dataset and the four pucker-pair sets are plotted in Figure S3, while Table S3 and Figure S4 provide the equivalent information for the 8b0x dataset.

Notably, we are not trying to analyze low detail suites where our presumptions are not met, i.e., where phosphates or glycosidic bonds are not placed well in density or do not have any density.

### 2.2 Conformer classes, MINT-AGE clusters and RNAprecis

**Conformer classes.** For each suite in the high-detail training data HD, the suite conformer class was determined with `phenix.suitename`. `phenix.suitename` uses the complete 7 dihedral angles of the suite (Figure 3) to assign the suite to one of the manually defined rotameric conformers (e.g. **1a, 1c, 5z**, etc.) or to **!!** if the suite does not match any recognized conformation. The conformer class assignments from `phenix.suitename` represent the current (best available to date, to best knowledge of the authors) “ground truth” suite identities for the HD datasets [12].

Each suite conformer class corresponds to a single pucker configuration pair—identifiable from the *ν* dihedral angle at high detail—and training data suites were assigned to pucker-pair sets accordingly. The HD and LD datasets comprise 4148 suites that qualified for analysis (3529 are P33; of those, 3183 are **1a** and 346 non-**1a**; 216 are P23; 265 are P32; and 138 are P22.)

**Clustering** of the four high-detail datasets has been performed based on the purely statistical method of MINT-AGE developed by [31]. In order to accommodate very low conformer class sizes, we have used the new *mode hunting method for very small dataset sizes* from [14]. Tuning parameters have been learned such that the *clusters* found are in high agreement with the above suite conformers, see Section 3.

**MINT-AGE training** was carried out on the HD datasets. The HD datasets are based on a preliminary version of the ‘nosuiteout’ RNA2023, as described above and available at https://zenodo.org/doi/10.5281/zenodo.8103013 [32]. The files that have been used are listed in Appendix A. The source PDB files were downloaded from the RCSB PDB database [37]. MINT-AGE used the full 7 dihedral angles to automatically cluster suites into clusters similar to the manually-defined suitename conformer classes.

**RNAprecis training** was carried out on the LD datasets. The LD datasets comprise a minimal set of atom positions for the same files and suites as the HD datasets. The LD datasets are split into pucker-pair sets based on the Pperp ribose pucker criterion described above in Section 1.1, which requires only atom positions from this minimal set. Because the training data is filtered to remove any ambiguous or incorrect ribose puckers, the pucker-pair sets are also reliably identified by *ν* values.

**RNAprecis testing** was carried out on the LDT and 8b0x datasets. LDT datasets were based on the 3.328 version of the http://rna.bgsu.edu/rna3dhub/nrlist RNA list, dated 27.3.2024; files were downloaded from the RCSB PDB on 1.4.2024. The high-detail atom positions needed to determine *ν* angles were not considered reliable for test set suites, so the suites were separated into pucker-pair sets based on the low-detail Pperp criteria (Figure 8). The LDT test set comprises 244 suites: 136 P33, 38 P32, 50 P23, and 20 P22. The 8b0x test set comprises 178 suites: 98 P33, 40 P32, 26 P23, and 14 P22. The PDB files and residues used are listed in Section B in the appendix.

### 2.3 Non-Euclidean statistical learning methods for classification

#### 2.3.1 Multicentered coordinates for low-detail constrained size-and-shape

Low-detail data is given by 5 *landmarks*

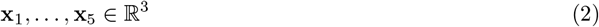

of atom center positions in three dimensions. Here the 5 landmarks denote the estimated atom centers of *N* 1*/N* 9_*i*−1_, C1^*′*^_*i*−1_, P^*′*^_*i*_, C1^*′*^_*i*_ and *N* 1*/N* 9_*i*_ in that order, see Figure 3.

**Size-and-shape** information of such a configuration is given by filtering out Euclidean motions (translations and rotations), and to this end, as alluded to in the introduction, we single out a *frame* of prominent landmarks as illustrated in Figure 5: The P atom center naturally serves as a reference point, i.e. its position is subtracted from all other atom centers. P in conjunction with the two C1^*′*^ atom centers naturally defines a planar nondegenerate reference triangle, which we rotate into the (*x, y*)-plane such that the rays from *P* to each of the two C1^*′*^ have the same projection onto the positive *y*-axis and such that the ray to the later-in-sequence C1^*′*^ (see Figure 3, usually but not always the longer of the two rays) has a positive projection onto the *x*-axis. Then, the opening angle *α* at the P atom of the reference triangle lives on a proper half circle. The two landmarks giving the N1/N9 atoms, however, are not centered with respect to the P^*′*^ atom but with respect to their neighboring C1^*′*^ atoms defining the glycosidic bond. Since, as noted in the introduction, these bonds lengths can be considered as a known constant, only the directions are variable, thus the coordinates of the two N1/N9 atoms can be viewed to live on a unit sphere.

**Figure 5:**
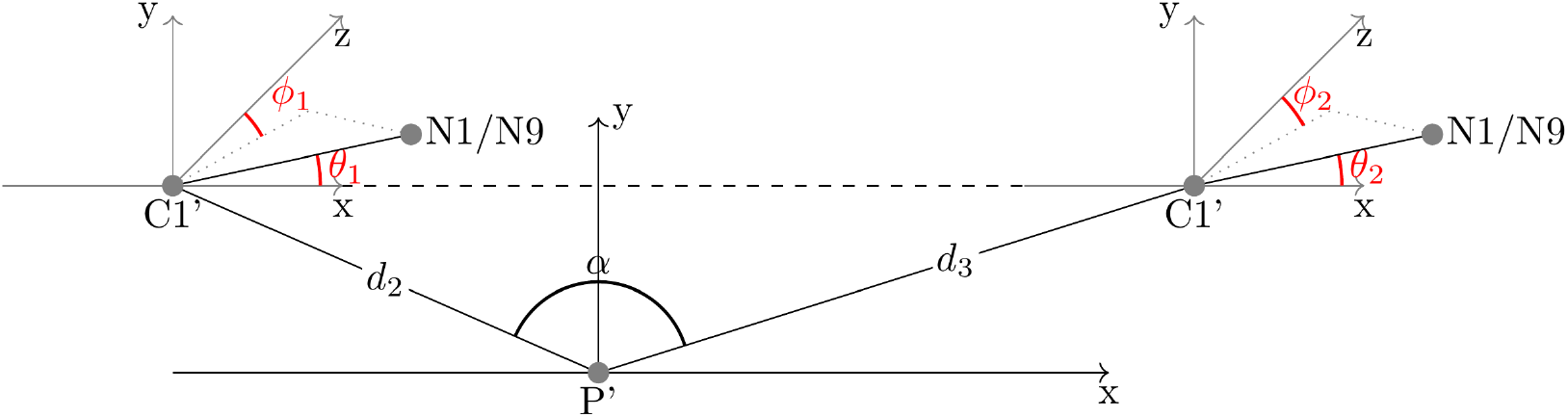
MUCCSS coordinates for LD suite size-and-shape. Part of the parameters describes the triangle spanned by the phosphorus as foot point and the two neighboring C1^*′*^ atoms. After normalization for rotation and height, three descriptors remain: the lengths of the two pseudo-bonds and the opening angle. The rest of the parameters describe the two N1/9 atoms at the corresponding glycosidic bonds (which have fixed lengths) each expressed in spherical coordinates with respect to the x-y-plane defined by the triangle.

**Detailed derivationof coordinates.** As sketched above, first subtract the phosphorus’ coordinates from the two C1^*′*^ coordinates and the respective C1^*′*^ coordinates from the corresponding *N* 1*/N* 9, to obtain the 4 new landmarks

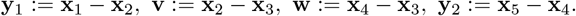

In order to rotate the triangle, which we may assume to be nondegenerate, spanned by the origin, **v** and **w** into the (*x, y*)-plane, as sketched above, define orthogonal unit vectors (where **e**_1_ and **e**_2_ define the new *x*- and *y*-axis, respectively)

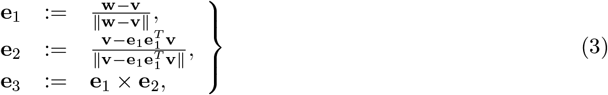

and the orthogonal matrix *R* := (**e**_1_, **e**_2_, **e**_3_). Indeed, then

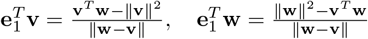

and

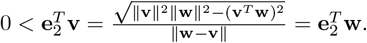

Thus, the *frame-based, multicentered coordinates* for the above 5-point configuration (2) are given by

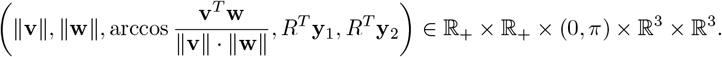

Taking into account that the two glycosidic bonds are constrained to have equal lengths that are the same for all configurations, we may assume ∥**y**_1_∥ = 1 = ∥**y**_2_∥, to obtain the MUCCSS (MUltiCentred Constrained Size-and-Shape) coordinates given by

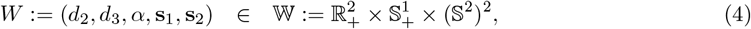

where

- 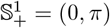is the unit half circle,
- *d*_2_ := ∥**v**∥ and *d*_3_ := ∥**w**∥ denote the distances between the atomic centers of P and the two C1^*′*^, respectively,
- 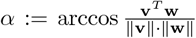 gives the angle at P between the two pseudo-bonds from P to the two C1^*′*^, respectively, and
- **s**_*i*_ := *R*^*T*^ **y**_*i*_, *i* = 1, 2 give the directions of the two glycosidic bonds, respectively, with polar angles *ϕ*_1_, *θ*_1_ and *ϕ*_2_, *θ*_2_, respectively (see Figure 5), *θ*_*i*_ ∈ [0, *π*], *ϕ*_*i*_ ∈ [0, 2*π*), *i* = 1, 2.

Notably, MUCCSS coordinates for LD data have 7 degrees of freedom, this is also the number of degrees of freedom for HD data modeled on the 7-dimensional torus, see (1).

**Metric, means, tangent space coordinates and tangent space covariance.** Since the space of MUCCSS coordinates for LD data, given by (4), is a direct product, its canonical metric is that of a direct product:

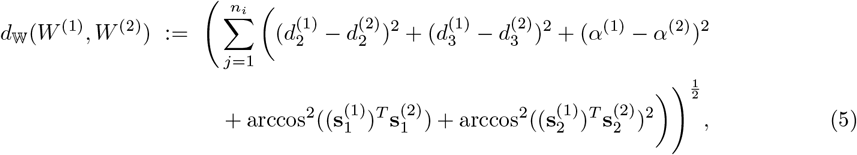

For

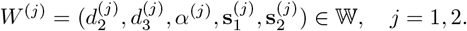

Further, for a sample

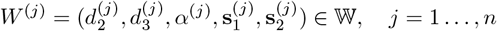

of suite representatives in LD we have their *Fréchet function*

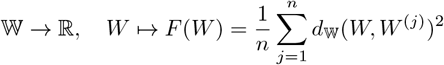

and every minimizer

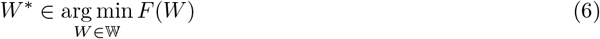

is a *Fréchet mean*.

Then, every *W* = (*d*_2_, *d*_2_, *α*, **s**_1_, **s**_2_) ∈ *W* has the following *tangent space coordinates* in the tangent space of *W* at 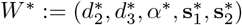, given by

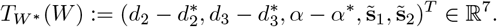

Here, 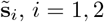, is obtained as follows: With the unit vectors from (3) select

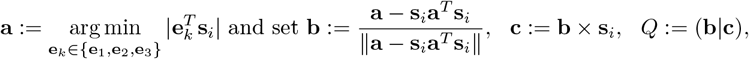

to obtain

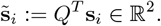

This yields the orthogonal projections of **s**_*i*_ (*i* = 1, 2) to the tangent space of *S*^2^ at **s**_*i*_ in a coordinate system only depending on **s**_*i*_.

Notably, for numerical robustness, **b** and **c** have been chosen from {**e**_1_, **e**_2_, **e**_3_} such that **s**_*i*_ is mainly explained by them. Note also that the mapping 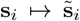 is only bijective if **s**_*i*_ is in the hemisphere {**s** ∈ *S*^2^ : **s**^*T*^ **s**_*i*_ ≥ 0}. The data driven choice of **b** and **c** results is a robustification of the well known as *residual coordinates* [38].

Finally, the tangent space coordinates of a sample

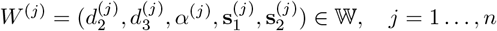

at a Fréchet mean 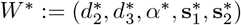, allow for the definition of a *tangent space covariance matrix* given by

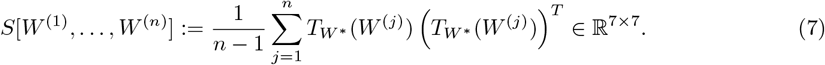

##### Remark 2.1 (Uniqueness).

*The RNAprecis algorithm below requires Fréchet means of MINT-AGE clusters in W representation. Due to [39], such sample means are unique on a complete connected Riemannian manifold if their distribution is absolutely continuous with respect to Riemannian measure. Indeed*, *W is a connected Riemannian manifold. Although W is noncomplete, as the boundary points corresponding to α* = 0, *π are missing, it is geodesically convex, since the standard Riemannian product metric inducing the metric (5) is Euclidean on the problematic* 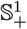. *Hence, these boundary points will never be coordinates of Fréchet means. In consequence, we can assume uniqueness of means and tangent space coordinates, and hence, also of tangent space covariance matrices*.

#### 2.3.2 RNAprecis

The following algorithm transforms LDT data to MINT-AGE clusters (introduced in Section 2.2) of HD gold standard data (see Section 2.1) and via its correspondence with suite conformation classes to suite conformations (see Figure 1).

##### Algorithm 2.2 (RNAprecis).

***Input***:

1. *Clusters obtained by MINT-AGE from HD gold standard data*

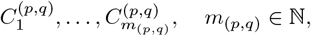 *for every pucker configuration pair* (*p, q*) ∈ *{*2, 3*}*^2^,
2. *for every cluster* 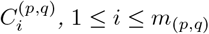 *a Fréchet mean* 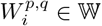 *from (6) of its LD representatives along with the corresponding tangent space covariance matrix* 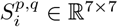 *from (7)*,
3. *a regularization parameter* 0 *< λ <* 1 *(with default value λ* = 0.5*) and for all* (*p, q*) ∈ *{*0, 1*}*^2^, 1 ≤ *i* ≤ *m*_(*p,q*)_ *the* regularized covariance matrix

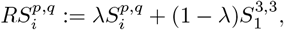
4. *a test suite in LD representation*

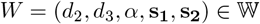

*from the LDT test data*.

***Steps:***

1. *Determine the sugar pucker configuration pair* (*p, q*) ∈ {2, 3 ^2^}*of W via the Pperp criterion (see Section 1*.*1)*,
2. *using cluster sizes* | 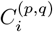 | *as (unnormalized) prior probabilities, determine (unnormalized, logarithmized) posterior probabilities*

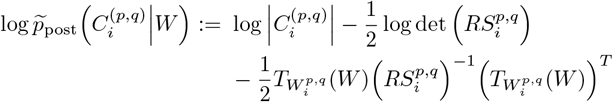

*and normalize them*

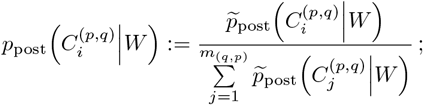
3. *determine the most probable cluster index*

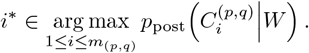

***Output***:

*The conformation class(es) corresponding to the MINT-AGE cluster* 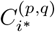.

##### Remark 2.3.

1. *RNAprecis employs a Gaussian likelihood with estimated covariance for each cluster in order to ensure that variation of all coordinate representatives is taken equally into account: the* Mahalanobis distance *[40]*,

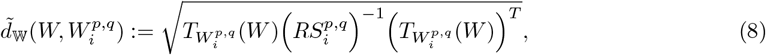

*is the only term in the posterior probability which depends on the test suite coordinates. Then for fixed W-distances between a test suite and a cluster mean, the more concentrated the cluster, the larger the distance* 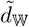.
2. *Regularization for the covariance with the overwhelming largest cluster (***1a** *has over 3,000, the second largest* **1c** *has only approx. 150 members) guarantees invertibility also for low-size clusters whose variation does not cover the entire 7-dimensional space*.
3. *The interpretation of posterior probabilities must take into account the covariance regularization. While these probabilities are valid posterior probabilities, they are connected to a model with regularized covariances, giving “regularized” credibility regions*.

## 3 Results

### 3.1 Clustering of HD and corresponding LD regions

In this section, the cluster results of the adapted MINT-AGE algorithm (see Section 2.2) for the individual data sets HDP33, HDP32, HDP23 and HDP22 (see Section 2.1) are reported and analyzed. The AGE algorithm requires 3 tuning parameters (see [19] and [31]) listed in Table 1 for the four different data sets. The resulting clusters are listed in Table S4 in Appendix C. Notably, as the confusion matrices in Figures S9, S10, S11 and S12 (all in Section C of the appendix) teach,

8 out of 11 clusters in the HDP33 data set,
13 out of 15 clusters in the HDP32 data set,
10 out of 16 clusters in the HDP23 data set,
7 out of 10 clusters in the HDP22 data set

belong to a single suite conformer class. Figure 6 shows the dihedral angle combination of *α* and *β* for the data set HDP33 (by far the largest) in the left panel (see Figure 3 and Figure 4 for an explanation of the dihedral angles), in which the MINT-AGE clusters are best visually separated. The plots of all angle combinations for all data sets are shown in Appendix C. There is also a detailed comparison between the cluster results and the suite conformer classes identified with `suitename`, using confusion matrices.

**Figure 6:**
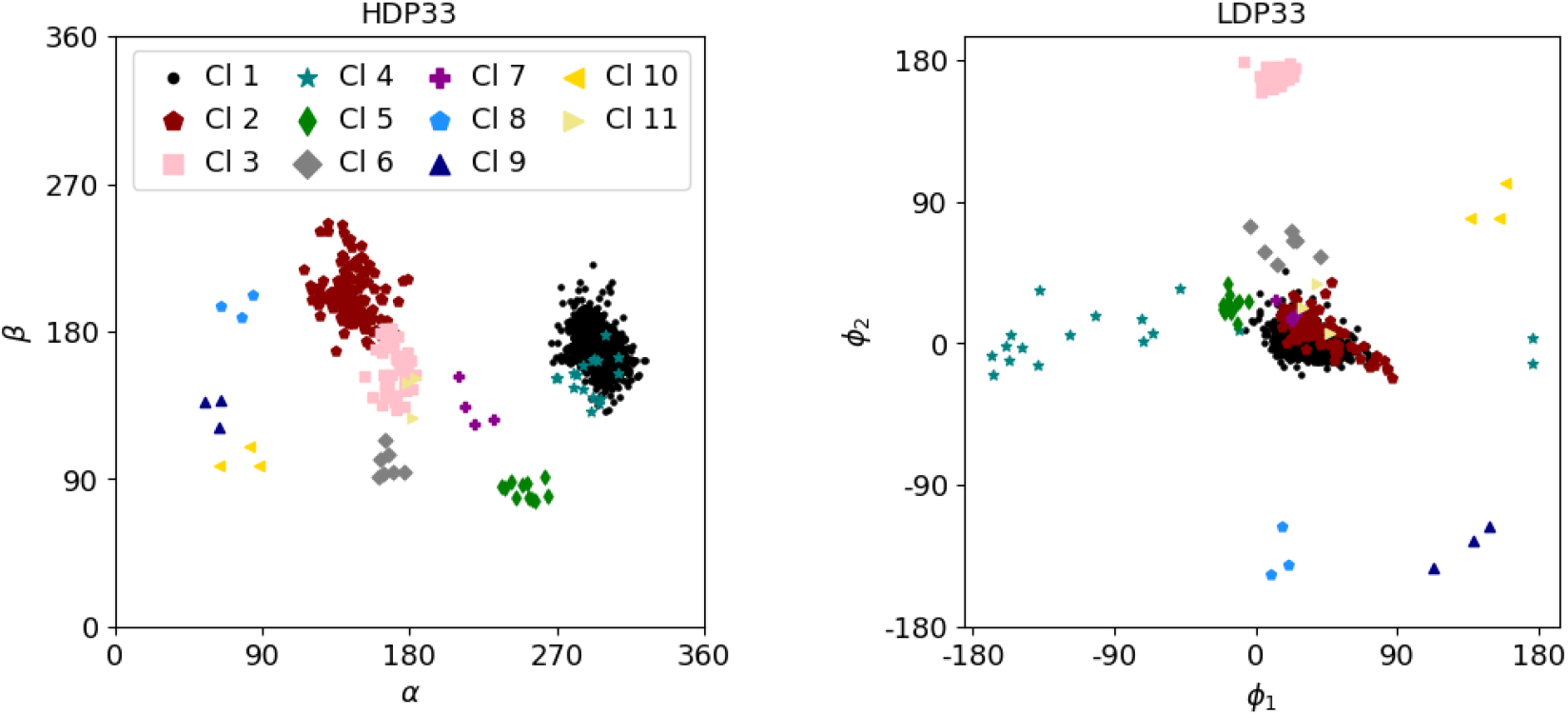
Left panel: Illustration of HDP33 clusters in the *α*-*β* plane of the high-detail coordinates. Right panel: The low-detail coordinates corresponding to the same suites, representing the distribution of the clusters in the *ϕ*_1_-*ϕ*_2_-plane. One can see that many of the clusters already well visible in the dihedral angle plot (left panel) are mapped to distinct clusters also in the low-detail plot (right panel) of longitudes of glycosidic bond directions.

MINT-AGE successfully identifies clusters with small numbers of individual suites for many suite conformers. **7d, 5d, 5j, 4g, #a, 4d, 5p, 5q, 0b, 2u**, and **2o** are all assigned unique clusters based on 4 or fewer examples. Nevertheless, some conformers remain unclustered: **7a, 3d, 5n, 3g, 5r, 0k**, and **4s** could not be assigned to unique MINT-AGE clusters. Conformers **7a, 3d, 5n**, and **3g** each have enough individual suites for MINT-AGE clustering, but their data clouds are obviously too spread out to assign to a single cluster, while **0k, 5r**, and **4s** comprise only one individual suite each.

### 3.2 MINT-AGE clusters comprising multiple conformer classes

In broad overview (see Figures S9, S10, S11 and S12 in Section C of the appendix), HDP33 Clusters 1 and 4 combine two or more different conformer classes into a single cluster, as do HDP32 Clusters 1 and 5. It was expected that the corresponding suites would be difficult to separate automatically, as their conformer classes boundaries were based on structural context in the previous work. We now go into detail.

HDP33 Cluster 1 combines **1a, &a, 1L**, and **1m. 1a** is the dominant conformer class (A-form helix) across all RNA and exerts a strong influence on clustering. The **&a, 1L**, and **1m** conformer classes are shoulders on the main **1a** cluster, each with a distinct behavior, and are not cleanly separated from **1a**. Similarly to the **1b**/**1[** case, these satellite conformers are distinguished from **1a** more by context than by conformation.

HDP33 Cluster 4 combines suite conformer Classes **9a, 3a**, and possibly **7a**. These conformer classes differ only in the *ζ* dihedral. As in the other cases, the distribution is quite continuous, and the distinction is found in the suite’s context. **9a** conformers start or end loops, while **3a** conformers enable stack switches. **7a** conformers, of which a single instance is included in Cluster 4, are also stack switches.

HDP32 Cluster 1 combines suite conformer Classes **1b** and **1[**, as does Cluster 5. The **1b**/**1[** distinction is illustrative of the challenge that MINT-AGE faces in reproducing the manual conformer classes. In the suitename parameter space, **1b** and **1[** are closely related and share a continuous distribution with no clear conformational breakpoint between the behaviors. Instead the primary distinction comes from elsewhere in the structure: **1[** has a ligand or another base intercalated between the bases of the suite, while the bases of 1b are often stacked directly on each other. In defining the original suite conformer classes [12], the cutoff between **1b** and **1[** was manually set at *β*=206° to provide the cleanest possible separation between these behaviors.

Some clusters combine a main conformer with a small conformer that is otherwise unclustered. HDP33 Clusters 1 and 4 each included a single instance of **7a**. HDP23 Cluster 2 is majority **0a**, but also includes four **#a**. HDP23 Cluster 5 is majority **8d** and also contains two **4d**. HDP22 Cluster 6 does not contain a majority conformer, but combines three **2u** with two **2o**. These cases reflect the limitations of our relatively small training dataset; there are not enough examples of **#a, 4d, 2u**, and **2o** to create independent clusters for these conformers.

Several clusters include one or two instances of a conformer that is otherwise clearly identified in its own cluster. HDP23 Cluster 2 includes two instances of **4a**, which is otherwise cleanly assigned to Cluster 7. HDP23 Cluster 3 includes two stray **0i** and one stray **6j**. HDP23 Cluster 11 includes one stray **9d**. These cases are a reminder that RNA conformer space is complex and biological conformer boundary definitions are often somewhat ambiguous.

Finally, some conformers and clusters are particularly challenging to untangle. HDP23 Cluster 4 is majority **6g**, but contains two instances of **4g**, while **4g** is further split across Clusters 15 and 16. Cluster 16 also contains a single **4a**, which is otherwise assigned to Cluster 7. HDP22 Cluster 7 combines 4 instances of **4b** with two of **0b**, both of which conformers themselves are heavily split across multiple clusters. Expansion of the training dataset should help resolve these ambiguous conformers. However, the splitting of suite conformers into multiple clusters is not necessarily due to the limitations of the training data and may, as discussed below, present an opportunity to revisit the existing suite conformer definitions.

### 3.3 Conformer classes split across multiple MINT-AGE clusters

Some suite conformer classes were split into multiple clusters by MINT-AGE, most notably **0a** in HDP23, but also **6d** in HDP23, **7p** in HDP32, **1f** in HDP33, and **0b** in HDP22 (see Figures S9, S10, S11 and S12 in Section C of the appendix). For reproducing the suite conformer classes, it is easy to simply re-combine the clusters. However, the separation suggests that MINT-AGE has discovered real conformational divisions that were not visible during the previous work.

MINT-AGE strongly separates the **0a** conformer into two subclusters, with 16 suites in Cluster 2 and 12 suites in Cluster 6 (see Figure 7 and Table S4 in Section C of the appendix). These subclusters separate the two major structural motifs found within **0a**, the stack switch and the kink-turn.

**Figure 7:**
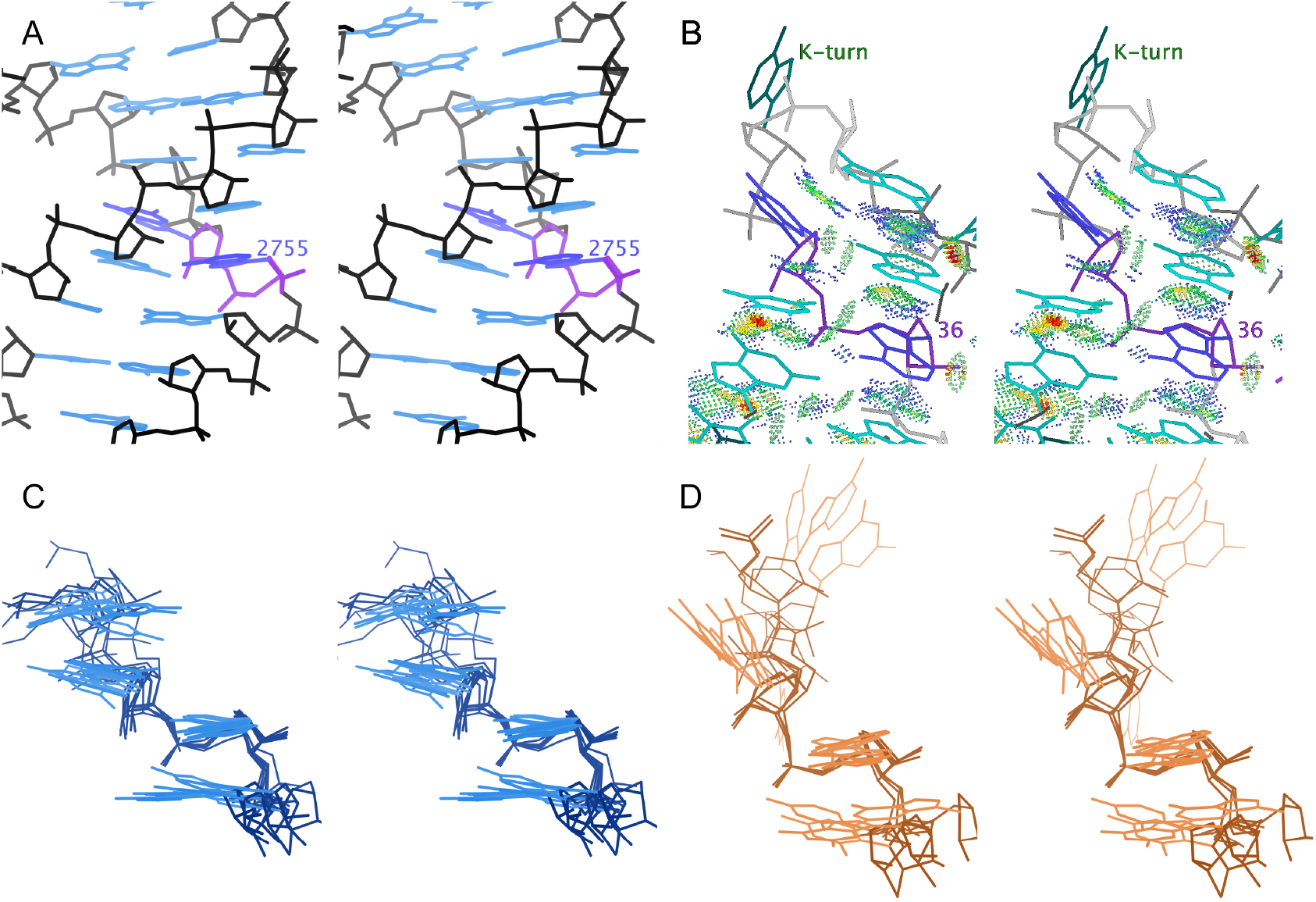
Stereo images of **0a** conformers, as separated by MINT-AGE. A. Stack switch from 8a3d chain A ([35]). Suite 2755 is highlighted in purple, and its bases stack into two different helix strands. B. Kink-turn from 5fjc chain A ([41]). Suite 36 is highlighted in purple. Below the outturned base, the upper base of suite 36 shows blue and green dots for favorable van der Waals packing with the backbone of the strand on the right. C. Superpositionof 6 representative MINT-AGE Cluster 2 suites. These are stack switches and have bases close to parallel. D. Superpositionof 4 representative MINT-AGE Cluster 6 suites. These are kink-turns and have bases at a high angle to each other.

Cluster 2 contains stack switches. In standard **1a** conformation, the subsequent bases of the suite are near parallel and stack on top of each other. In these stack switches, the bases are still roughly parallel but change from base-stacking with each other to stacking with a different strand, usually in the same helix (Figure 7 A and C).

Cluster 6 contains kink-turns. The kink-turn motif [42] spans multiple suites – the most distinctive feature is a very tight turn with an outturned base that often interacts with proteins. The Cluster 6 suite comprises the two residues following the residue with the outturned base. The first base of the Cluster 6 suite stacks with the backbone of the residue on the other side of the outturned base, stabilizing the sharp turn. The bases of a Cluster 6 suite are angled relative to each other due to this interaction with the preceding backbone (Figure 7 B and D).

**0a** was previously treated as a single conformer class because these suites have a continuous distribution along the *ζ* dihedral angle, best seen in the *ζ/α* panel of Figure S6 in the appendix. However, Clusters 2 and 6 are more clearly distinguishable by *ϵ*, which also separates the related conformer Class **4a**/Cluster 7, best seen in the *ϵ/ζ* panel of Figure S6 in the appendix.

The split of suite conformer Class **0a** provides the strongest evidence of MINT-AGE identifying a systematic functional/conformational distinction that was not previously recognizable. The **7p** split in HDP32 is also suggestive. MINT-AGE finds 4 clusters, combining 7 instances into Cluster 9, 5 into Cluster 10, 4 into Cluster 13, and 3 into Cluster 14. This set of clusters is complex, but Cluster 9 is clearly conformationally distinct from the rest (Figure 8). That this distinction is conformational rather than “just” contextual demonstrates the value in approaching suite conformer identification from both perspectives.

**Figure 8:**
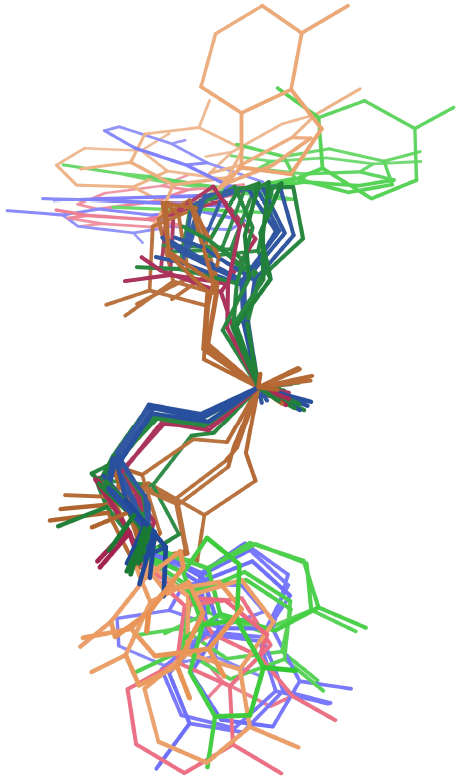
Superpositionof **7p** suites, colored by MINT-AGE cluster. Cluster 9 is orange, 10 is green, 13 is blue, 14 is red. Cluster 9 is clearly conformationally distinct from the other groupings within **7p**.

The **0a** and **7p** suite conformer classes should be split into new conformer classes, based on the MINT-AGE clustering. The success of an unsupervised method in recognizing new conformer classes not previously identified by manual work is an encouraging result.

### 3.4 Statistical classification of conformations for low-detail data

The detailed results of the RNAprecis test are listed in Section E in the appendix. The results are presented individually for the four pucker-pair data sets LDTP33 (Section E.1), LDTP32 (Section E.2), LDTP23 (Section E.3) and LDTP22 (Section E.4) formed by the classification in Step 1 of Algorithm 2.2. Analogously, 8b0xP33 (Section E.5), 8b0xP32 (Section E.6), 8b0xP23 (Section E.7) and 8b0xP22 (Section E.8) are treated separately. For each of the test elements, the probable answer conformer class determined by related experimental structures is provided for comparison (see paragraph “Low-detail test data and related conformer classes as probable answers” in Section 2.1). There are three possibilities, whose colors correspond to tables and figures below:

**Prediction matches.** The cluster determined by RNAprecis (Algorithm 2.2) contains a suite conformer class that matches the sequence equivalence-related probable answer for this element (see Table S4 in the appendix). Note that clusters containing multiple conformer classes, such as P32 Cluster 1 which contains both **1b** and **1[**, are considered a match for any of their member conformers.

**Prediction mismatches.** The cluster determined by RNAprecis (Algorithm 2.2) contains no members matching the probable answer for the test element (see Table S4 in the appendix). However, the probable answer conformer is a member of the same pucker-pair set as the test element.

**Pucker mismatches.** The pucker configuration pair of the cluster determined by RNAprecis from the test element (in Step 1 of Algorithm 2.2) does not match the pucker-pair set of the probable answer.

#### 3.4.1 Results for LDT test set

For all four pucker-pair sets except P23, at least 2/3 of the elements are sorted into the prediction match group (blue), see Table 2 and top left in Figure S17, Figure S18, Figure S20 and Figure S22 in the appendix. The lower success rate for the P23 pucker-pair set is not clearly associated to a specific type of misclassification. However, in the P33 pucker-pair set, suites with probable answer **1a** are in large numbers classified as **1c**, a systematic effect discussed below. The probable sequence-related conformer class answers and the clusters determined with RNAprecis for all individual elements (sorted according to the four pucker-pair sets and according to the blue, red and orange categories) are reported in Table S5 (blue category of LDTP33), Table S6 (red and orange category of LDTP33), Table S7 (blue, red and orange category of LDTP32), Table S8 (blue, red and orange category of LDTP23), and Table S9 (blue, red and orange category of LDTP22). A comparison between the probable answers and the clusters determined by RNAprecis is shown in a classification matrix for the four data sets in Figure 9 below and in Figures S19, S21 and S23 in the appendix. The prediction results are separated into the four pucker configuration pairs so that the **1a** conformer does not dominate the results. The RNAprecis prediction results are good uniformly over all pucker-pair sets and all clusters, even small clusters.

**Table 2:** List of the four pucker-pair test datasets in the first column formed by Step 1 in RNAprecis (Algorithm 2.2) with the respective total counts in the second column and counts and percentages in the third through fifth columns.

| dataset | count | prediction matches | prediction mismatches | pucker mismatches |
| --- | --- | --- | --- | --- |
| LDTP33 | 136 | 91 (67%) | 37 (27%) | 8 (6%) |
| LDTP32 | 38 | 30 (79%) | 3 (8%) | 5 (13%) |
| LDTP23 | 50 | 28 (56%) | 15 (30%) | 7 (14%) |
| LDTP22 | 20 | 16 (80%) | 2 (10%) | 2 (10%) |

**Table 3:** List of the four pucker-pair test datasets for the 8b0x test data in the first column formed by Step 1 in RNAprecis (Algorithm 2.2) with the respective total counts in the second column and counts and percentages in the third through fifth columns.

| dataset | count | **prediction matches** | **prediction mismatches** | **pucker mismatches** |
| --- | --- | --- | --- | --- |
| 8b0xP33 | 98 | 53 (54%) | 32 (33%) | 13 (13%) |
| 8b0xP32 | 40 | 33 (82%) | 4 (10%) | 3 (8%) |
| 8b0xP23 | 26 | 20 (77%) | 2 (8%) | 4 (15%) |
| 8b0xP22 | 14 | 9 (64%) | 5 (36%) | 0 (0%) |

**Figure 9:**
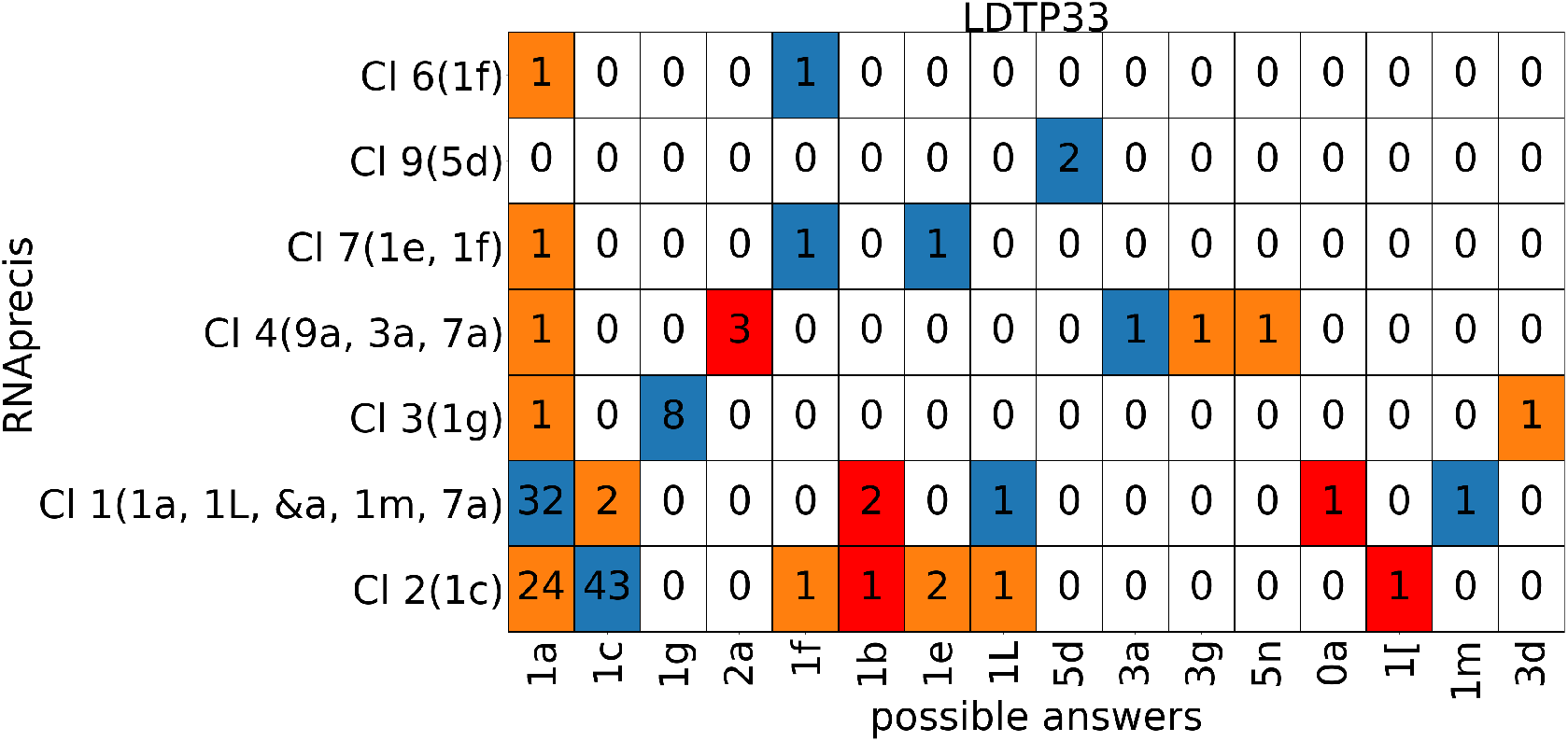
Classification matrix listing counts of probable sequence-related conformation classes (horizontal axis) versus MINT-AGE clusters (vertical, with conformer classes in parentheses from MINT-AGE training, see Figure S9 in the appendix) for the low-detail sugar test pucker-pair data set LDTP33 with colors: blue for match, orange for mismatch and red for pucker-pair mismatch. Columns listing multiple suite conformers are from sites solved in different conformations across sequence-related models, and any of the sequence-related conformations were accepted as a matching predictions.

The conformers **3d, 5n, 3g, 0k, 5r**, and **4s** were not clustered by MINT-AGE, probably due to having too few examples in the training dataset (between 1 and 4). Therefore, these conformers also could not be predicted by RNAprecis. Any test set suite where the probable answer conformation is one of these unclustered conformers will necessarily result in a prediction mismatch or pucker mismatch. This inability to predict matching conformers affects 3 suites in the test set, one each of **3d, 3g**, and **5n**.

#### 3.4.2 Results for 8b0x test set

In the “out of sample” test set from PDB 8b0x, RNAprecis again achieves high classification rates across the board, however in this case the performance on the P33 pucker-pair set lags behind due to the **1a** to **1c** misclassification also found in the LDT test set. Pucker mismatches are rare also in this test set. Notably, the 8b0xP23 configuration exhibits a lower rate of mismatches than the LDTP23, which further underscores that no systematic pattern of misclassification is apparent. Detailed results are presented in Sections E.5, E.6, E.7, and E.8 in the appendix.

Conformers not clustered by MINT-AGE were more prevalent in the “out of sample” test set. 9 prediction mismatches, including 3 **3d** suites, 1 **3g**, 2 **4s**, 1 **5n**, and 2 **5r** are attributable to unclustered conformers. This accounts for the worse apparent performance in the 8b0xP22 test, which contains the 2 **4s** suites, while LDTP22 contained no unclusterable suites.

While 8b0x a has provided a useful test for demonstrating that RNAprecis does not overfit, we expect that including this chain in future MINT-AGE training will allow clustering and prediction of additional suite conformers. In particular, we expect to be able to identify **4s**, which occurs as a well-defined part of the “S-motif” [43].

## 4 Discussion

### 4.1 MINT-AGE clustering

This clustering process is quite successful in both training and test stages. In the training stage with filtered, high-detail data, 38 of the 51 suite conformers with ≥ 3 individual suites (75%) were clustered with one-to-one agreement by MINT-AGE and only 4 were not clustered at all. The rest either combined 2 or more suite conformers into a single cluster or split one conformer into more than one cluster. Interesting and perhaps correct examples of those cases are described in Section 3 (Results).

For the testing stage, RNAprecis uses the trained cluster relationships in reverse to predict the high-detail suite conformer for each **!!** suite that has a probable valid conformer (or several) in related structures. 68% were clear wins of prediction matches to a sequence-related conformation, and 23% were prediction mismatches. The other 9% were mismatches of the Pperp-determined pucker-pair set (used by RNAprecis) to the pucker-pair set of the probable answers, indicating cases where the modeling errors were too severe for this method. The details of the two mismatch categories have taught us several feasible ways to make the RNAprecis process significantly better, described below.

The most obvious way to improve MINT-AGE performance is use more training data. The RNA2023 dataset, from which the HD and LD training sets were adapted, is a general purpose dataset. One of its purposes is to reflect the relative frequency of different features in RNA structures. This means that several rare suite conformers (e.g. **3d, 5n, 4s**) are too rare (see above) in the dataset to form clusters with MINT-AGE. A new dataset, built to include more conformational diversity and more rare suite conformers (for instance by allowing two entries per sequence equivalence class) would improve clustering and recognition of those conformers.

MINT-AGE clustering can also be improved through development of additional parameters. For instance, the N1/N9-to-N1/N9 distance across the suite may differentiate between stacking (shorter distance) and intercalation (longer distance) in **1b** and **1[**. Such parameters would be targeted to specific conformer families and invoked only when needed.

### 4.2 Pucker mismatches

The most concerning category of incorrect predictions are the pucker mismatches, although they account for only about 9% of the total predictions. In these cases, the ribose pucker combination predicted from a test suite’s Pperp distance test does not match the pucker configuration pair of the sequence equivalence class-related answer conformation. For example, 2qbz, chain X, suite 47-48 has a Pperp-predicted sugar pucker configuration pair corresponding to P23. The probable answer for this site (**6p**) has a different pucker configuration pair, corresponding to P22.

Quality-filtered residues from HD and LD are cleanly separated into pucker-pair sets by Pperp distance (Figure 10). Because the separation is so clear, MINT-AGE and RNAprecis treat each pucker-pair set separately, and RNAprecis prediction across pucker-pair sets is not possible. However, residues from the LDT test set can have ambiguous Pperp values near the 2.9Å cutoff (Figure 11). The otherwise useful Pperp criterion loses predictive power near this boundary.

**Figure 10:**
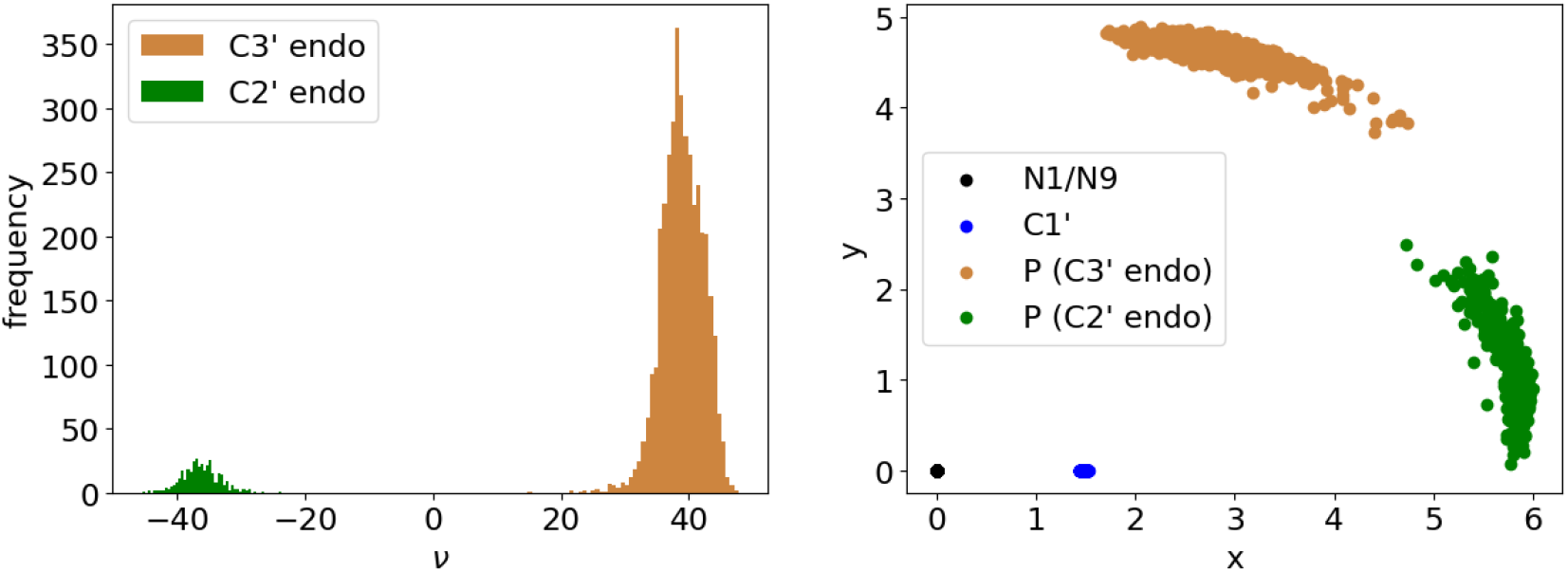
For the training data of both high quality and high resolution, the two different sugar puckers can be very well discriminated by the angle *ν* (left). Right: Using the coordinates described in Figure 2B, namely mapping *N* 1*/N* 9 to the origin and C1^*′*^ to the first positive axis (*x*), the perpendicular distance (Pperp, positive vertical *y* axis) similarly perfectly separates the two pucker configurations (this 2D display reflects true atom position in Euclidean space).

**Figure 11:**
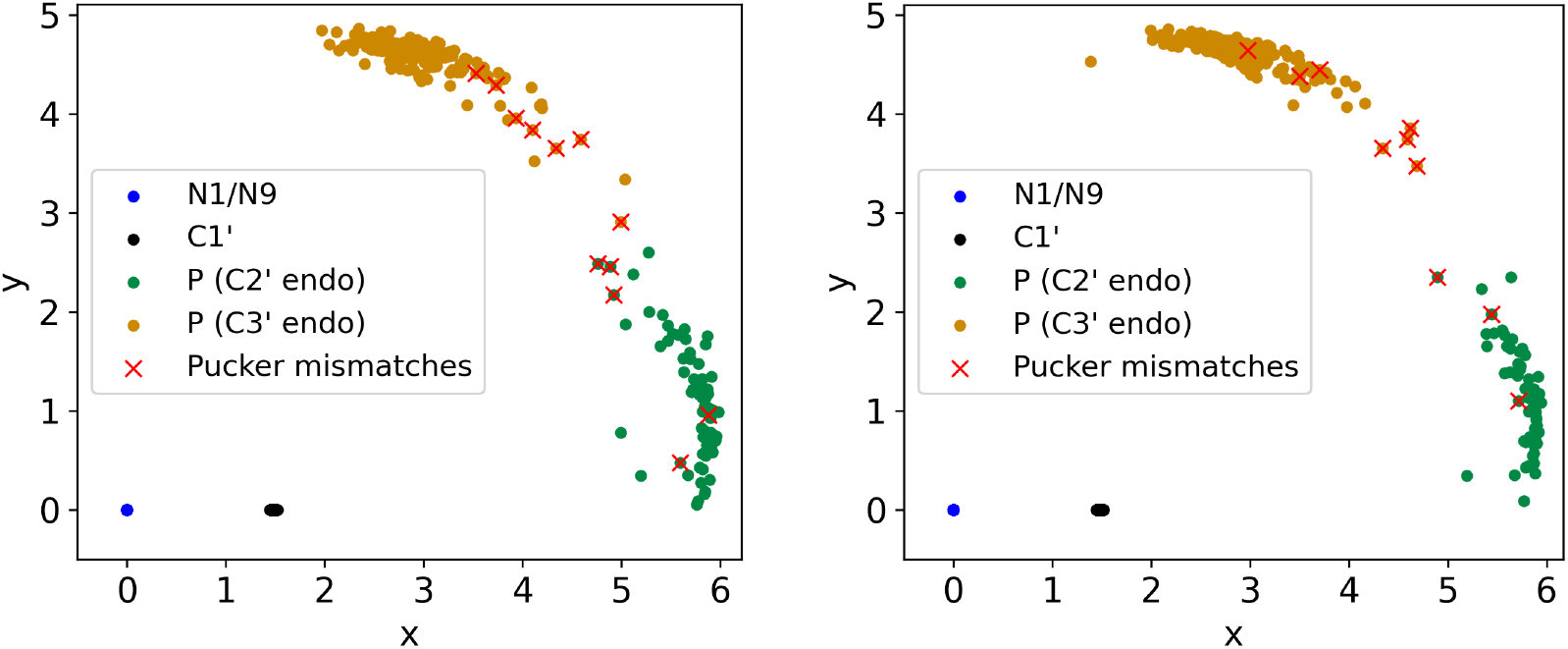
Depicting all low-detail test suites in the coordinates described in Figure 2B, namely mapping *N* 1*/N* 9 to the origin and C1^*′*^ to the positive *x*-axis. The left panel shows the respective coordinates for their first sugars (5^*′*^ end of the suite), the right panel shows the respective coordinates of their second sugars (3^*′*^ end of the suite). Considering *y* (the perpendicular distance to the positive *x*-axis), as in Figure 10, we predict the unknown true sugar pucker by vertically separating at 2.9, as learned from the high-resolution data, depicted in Figure 10. Matches of pucker predictions with probable answers are green and orange, respectively, red crosses indicate coordinates of pucker mismatches (see Section 3.4). There are only 12 pucker mismatches in the left panel and 10 in the right panel.

A poor experimental map—or a poor fit to map—is the clearest cause of pucker mismatches. If the phosphate is not modeled near the center of a high map peak, or if the base and ribose are not each in a blob with plausible local relationships, then the basic assumptions of RNAprecis are not satisfied. Figure 12 shows two such cases. Even without a pucker mismatch, RNAprecis is unlikely to find a correct answer in these cases, since the modeled suite does not necessarily bear any relationship to the correct conformation. Pucker mismatches thus represent a more severe category of modeling error than RNAprecis is intended to address, where the conformation of the model is very far from the intended structure.

**Figure 12:**
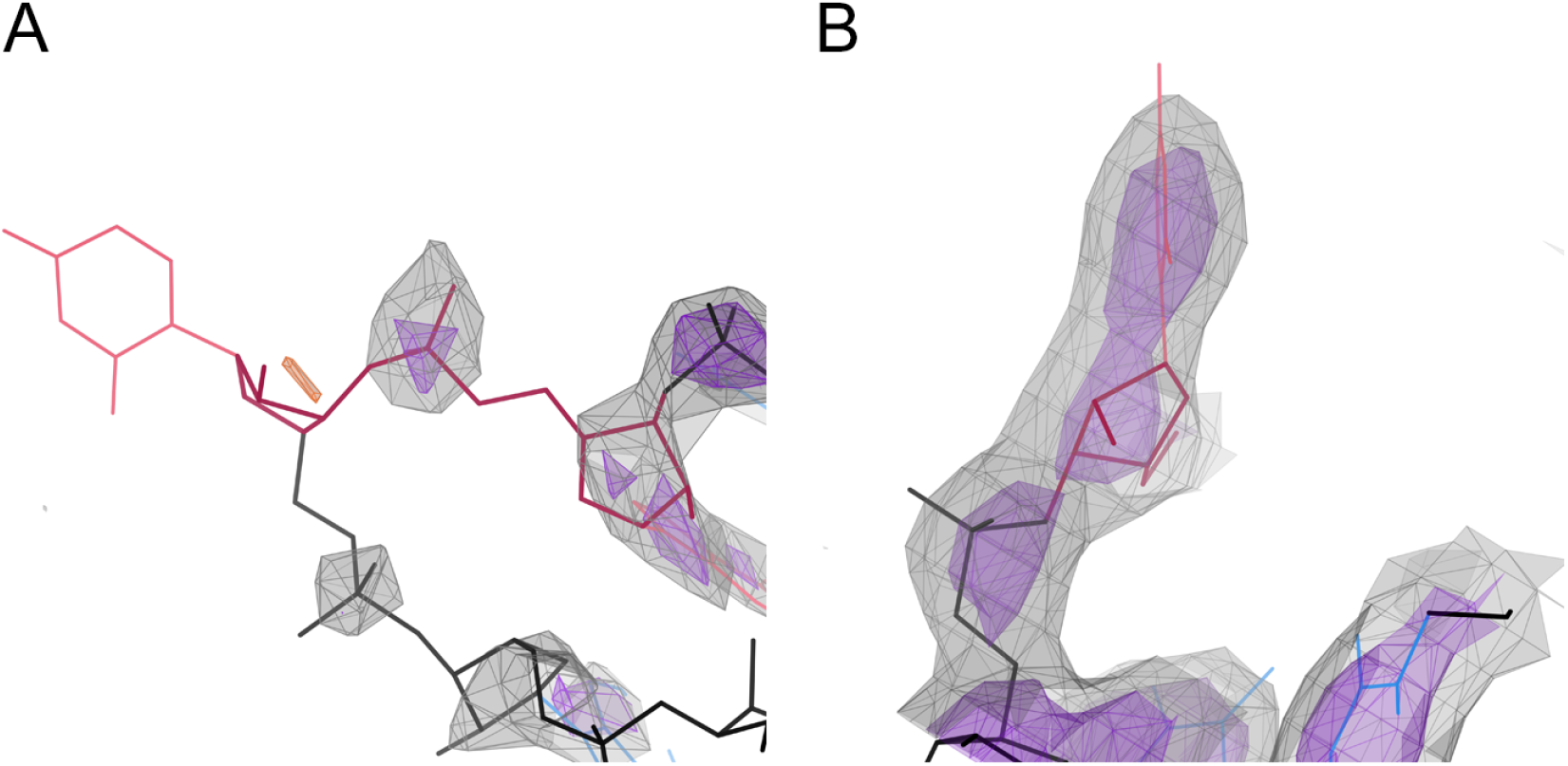
Two examples of problems causing RNAprecis pucker mismatches. A: Suite 47-48 of 2qbz, chain X, from LDT. Base 47 of this **!!** outlier suite has no density to define its position, and the corresponding ribose has a small difference density peak (orange, −3.5*σ*) which may indicate a misplacement. Map contours at 1*σ* (gray) and 1.5*σ* (purple). B: Residue 561 of 7st2, chain 3. This base is part of **!!** suites 560-561 and 561-562, both from LDT. This base is clearly modeled in the wrong orientation relative to its map, resulting in an incorrect glycosidic bond vector. CryoEM map contours at 6*σ* (gray) and 10*σ* (purple).

The test set likely under-represents the actual prevalence of regions with experimental maps that cannot confidently support a model. All suites in the test set were solved as a recognized conformer in a related structure, so truly disordered regions were selected against. While Pperp distances in cases of pucker mismatch are considerably closer on average to the 2.9 boundary, than for correct matches (see Figure 11), the variation is very high such that this property is not suited to predict pucker mismatches. Other methods that directly assess map or fit-to-map quality will be necessary to identify suites that are likely to contain pucker mismatches. In regions with very poor experimental support, it is usually better simply to build no model at all, rather than try to predict conformations from compromised data.

An illustrative case for the challenges in RNA backbone prediction is 2qbz, chain X, suite 138-139. This suite is predicted with a pucker mismatch, however, the key features are all reasonably well-placed in the map. The problem is in the preceding suite, 137-138. This suite has a severe error across the *α*-*β*-*γ* dihedrals, which results in an incorrect sugar pucker in ribose 138. This pucker error interferes with the RNAprecis prediction for 138-139. This sort of compound error requires an iterative validation-remodeling-refinement process to correct step by step.

### 4.3 1a vs 1c prediction mismatches

The classification matrix (Figure 9) for the LDTP33 portion of the test set shows that **1a** vs **1c** was one of the most difficult conformation pairs for RNAprecis to predict correctly. The difficulty is asymmetrical, with few **1c** suites being misidentified as Cluster 1, but many **1a** suites misidentified as Cluster 2.

A **1a**/**1c** ambiguity is not unexpected. A coordinated motion of *α* and *γ* allows ≈180^*°*^ changes in those dihedral angles to occur with minimal effects on the parameters that RNAprecis uses (Figure 13). This motion resembles that of a physical “crankshaft”, having opposite and equal changes in *α* and *γ* that drive changes in C5^*′*^/O5^*′*^ atom positions, while *β* is held unchanged. In the Figure 13 superpositions, the glycosidic bond vectors of the lower base are systematically different in **1a** versus **1c**, but the difference is subtle relative to the huge size and population of the **1a** conformer. RNAprecis therefore struggles to distinguish between these conformers.

**Figure 13:**
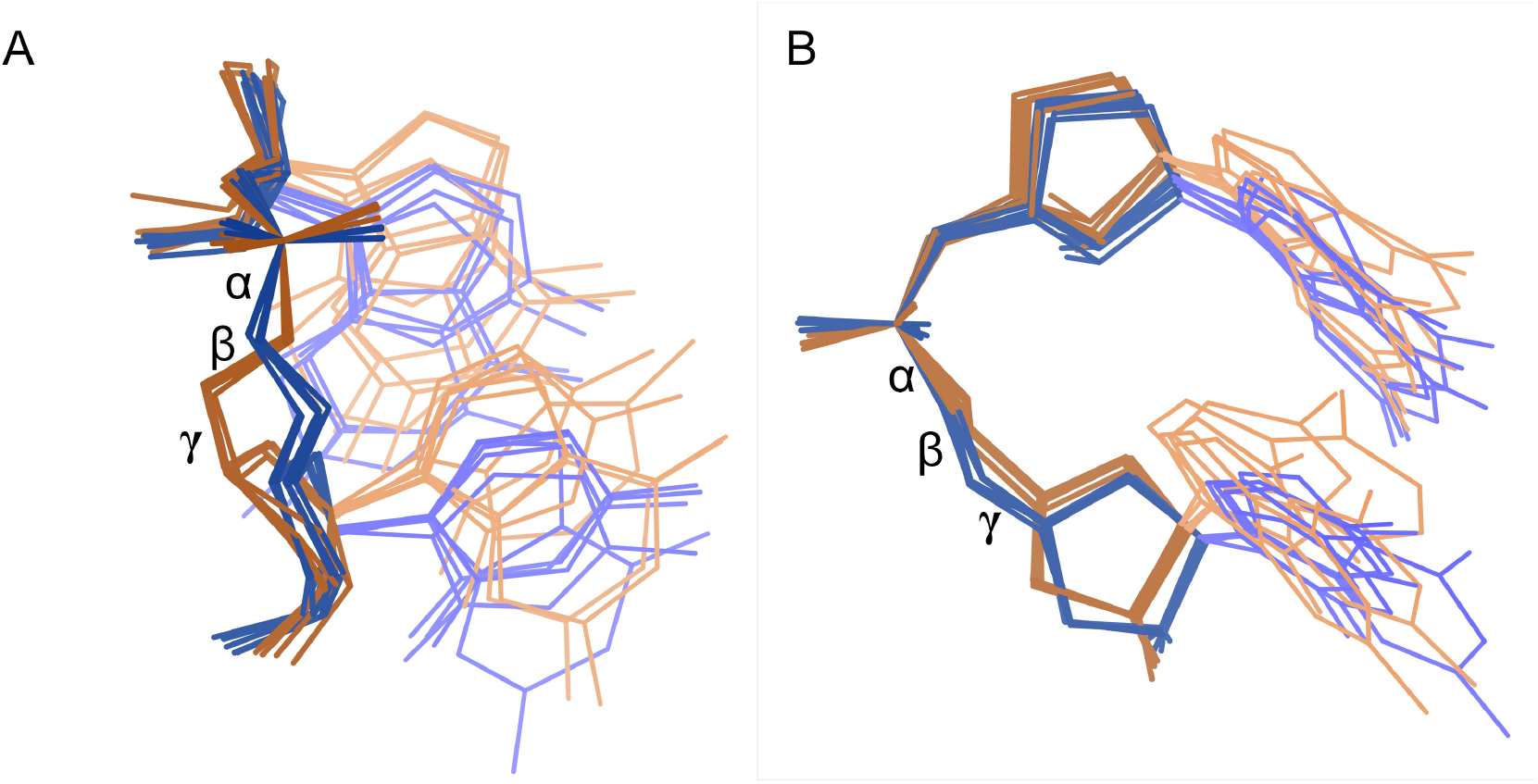
Superpositionof representative **1a** (orange) and **1c** (blue) suites. A. View showing full backbone detail. **1a** and **1c** are easily distinguished by differences in the *α* and *γ* dihedrals. Therefore, MINT-AGE can easily distinguish these conformers. B. Another view of the same superposition, showing the alignment of the riboses and bases. Despite the large difference in *α* and *γ*, the related changes in glycosidic bond directions are relatively small. Therefore, RNAprecis cannot readily see the difference between **1a** and **1c**.

We examined whether RNAprecis might be identifying the correct conformations in cases where the reference structure was mistaken. The reference structures of **1a**/**1c** prediction mismatches were manually reviewed based on visual agreement with their experimental maps. The reference structure suites showed fits to map ranging from plausible to definitely correct. This manual review of the **1a**/**1c** prediction mismatches also served to confirm that the sequence-related probable answers for the test set are reliable enough to use as targets. Where RNAprecis disagrees with the probable answers for this known difficult case, the sequence equivalence class-related conformation is generally correct.

The **1a**/**1c** ambiguity may be partially resolvable by increased reliance on the *d*_3_ parameter in this case. *d*_3_ is the distance across the relevant *α, β*, and *γ* dihedrals (see Figure 5). While there is some overlap between the Cluster 1/**1a** and Cluster 2/**1c** distributions in all parameters, *d*_3_ provides the cleanest single-parameter distinction between the two, as seen in the *d*_2_/*d*_3_ panel of Figure S13 in the appendix.

The asymmetry of **1a**/**1c** misidentification affords a pragmatic immediate solution. Cluster 2 should be considered a possible prediction of either **1c** or **1a**, much as Cluster 1 is considered a match to **1a** or any of its satellite conformers (but not **1c**) and Cluster 7 is considered a match to either **1e** or **1f**. In any of these cases, each member conformer of a cluster must be tested when making corrections to a structure.

### 4.4 Other prediction mismatches

We manually examined all non-**1a**/**1c** prediction mismatches, comparing the test set suites to their experimental maps and to their corresponding probable answer conformations using the KiNG viewer. We did not find a unifying explanation for prediction mismatches, but some features of interest emerged.

Several prediction mismatch test suites from ribosome structures—including 7f5s chain L5 suite 4554, 8eiu chain A suite 195, and 7st2 chain 3 suites 149, 306, 519, 642, 1502, and 1519—showed conformations similar to their references answers except for a rotation of the *α*/*γ* crankshaft. Given RNAprecis’s difficulty resolving the **1a**/**1c** conformers, which differ by this same crankshaft motion, this collection of suites is suggestive of a systematic challenge. However, outside the ribosome examples, only one prediction mismatch suite, namely 3ger chain A suite 54, showed a similar crankshaft motion between test and reference. This suite’s probable answer was **5n**, one of the conformer classes not clustered by MINT-AGE (since there were only 4 examples in the HDP33 training set), so a prediction match was impossible in this case. This suggests that this collection of suites instead represents a difficult-to-interpret modeling error idiosyncratic to these ribosome structures, especially 7st2.

Several test suites showed high similarity to the conformations of their respective answer suites, including 2qbz chain X suites 102 and 155, 5b2o B 18, 7st2 3 116 and 564, 6b3k R 58, and 8eiu A 13. In all of the listed cases, the answer suite had a low suiteness score less than 0.1. *Suiteness* is a score between 0 and 1 generated by `phenix.suitename` that reflects the distance between an observed suite conformation and the canonical conformation. The low suiteness scores of these answer suites indicate that the observed conformations are on the border between recognized conformations or are otherwise unusual. We expect that an expanded training set would improve prediction quality around conformer class boundaries. We found that 26% of the non-**1a**/**1c** prediction mismatches and 18% of the successful prediction matches had an answer suite with suiteness less than 0.1. Therefore, target conformers near class boundaries are somewhat more difficult but are not a systematic impediment to RNAprecis prediction.

Notably absent from the list of prediction mismatches are suite conformers with small MINTAGE clusters. Only one such test suite, namely 2qbz chain X suite 30, has a probable answer whose largest MINT-AGE cluster contains fewer than 5 members, **4g** with 4 members. The prediction match list contains 9 such suites. These suites represent 3% of the mismatches and 5% of the matches, indicating that MINT-AGE successfully produces clusters for low-population conformers, and that MINT-AGE cluster size is not a limiting factor in RNAprecis predictions.

### 4.5 Other methods

Updates to the suite conformer classes based on new data and MINT-AGE clustering will improve any methods that draw on the consensus conformer definitions. The emphasis in programs similar to ours has shifted. The excellent Erraser [7] and Brickworx [44] appear unavailable at time of writing. There are now more aids for getting approximate structures (that is, without consideration of details such as pucker or crankshaft) from as few as a single atom per residue, such as Arena [45]. There have been major developments in prediction, and in the related big calculations of approximate structures from 5–10Å resolution cryoEM data, such as Ribosolve [46]. And there are exciting new cryo-EM structures of ribosomes at very high resolution [34] and of ornate RNA-only structures at about 3Å resolution [47]. But for getting the details right at relatively high resolution, our closest comparison is RCrane [48], which is available and widely used in Coot [49]. It provides structure building of RNA from the phosphate and base and uses the Pperp and backbone conformers from our joint paper [12]. It does an excellent job. However, RCrane is primarily a model building tool, where RNAprecis is focused on model correction and on particularly difficult outliers that persist after refinement. The LDT test set demonstrates that solvable modeling errors still occur in recent RNA structures. In this sense, the methods are complementary.

## 5 Outlook

Ultimately, we intend to develop RNAprecis predictions into an RNA structure validation and recommendation tool for use in Phenix [10], MolProbity, and similar platforms.

Challenges in RNAprecis prediction have identified additional targeted structural tools to be developed alongside RNAprecis. The **1a**/**1c** ambiguity may yet be resolvable with the RNAprecis parameters such as the *d*_3_ distance. But such difficult cases might also be addressed with paired, comparative local refinements, once RNAprecis identifies a likely site. Automatically identifying suites where the critical RNAprecis atoms are not well-aligned with the map will be a more challenging problem, but would benefit both model building and RNAprecis predictions, especially avoiding pucker mismatches. RNAprecis predictions may be enhanced by employing a more suitable distribution model for clusters. Firstly, a larger training set could make it possible to forego the regularization of cluster covariances, thereby improving the probabilistic interpretability of posterior probabilities as probabilities for a certain cluster. Secondly, the Gaussian likelihood model used here for simplicity may not provide the best fit to the data. It may be worthwhile to explore more general probability distribution models to better capture the data distribution in low-detail coordinates.

MINT-AGE clustering provides an opportunity to revisit and expand previous suite definitions. Suite conformers such as **0a** and **7p** which contain multiple subclusters with distinct enough conformations to merit their own suite identities have been discussed above. Additionally, there may exist additional valid suite conformers that were too rare to be detected at all at the time of the previous work. In the current work, unrecognized suite conformers (**!!**s) were intentionally excluded from training. MINTAGE clustering can be performed on the **!!**s as well, to identify any new clusters of sufficient distribution and consistency to merit recognition as suite conformers, though human inspection will be required to distinguish between real conformations and decoys derived from common modeling errors.

## Code availability

The code used to generate the analyses and plots presented in this paper can be found at https://gitlab.gwdg.de/henrik.wiechers1/rnaprecis.

## 6 Acknowledgments

BE, SH and HW acknowledge support from DFG SFB 1456; SH acknowledges support from DFG HU-1575-7, DFG GK 2088 and the Niedersachsen Vorab of the Volkswagen Foundation; CJW, MGP, VBC, and JSR acknowledge support from NIH P01-GM063210 and the Phenix Industrial Consortium.

## Appendices

### A High detail training data set (HD)

This section introduces in Table S1 the PDB files with the corresponding resolution of the high resolution training data set (introduced in Section 2.1), from which the HD and LD data sets (and the corresponding four pucker-pair sets) are composed. In Figure S1 the four pucker-pair sets of HD are plotted and in Figure S2 the four sub-data sets of LD are plotted.

**Table S1:**
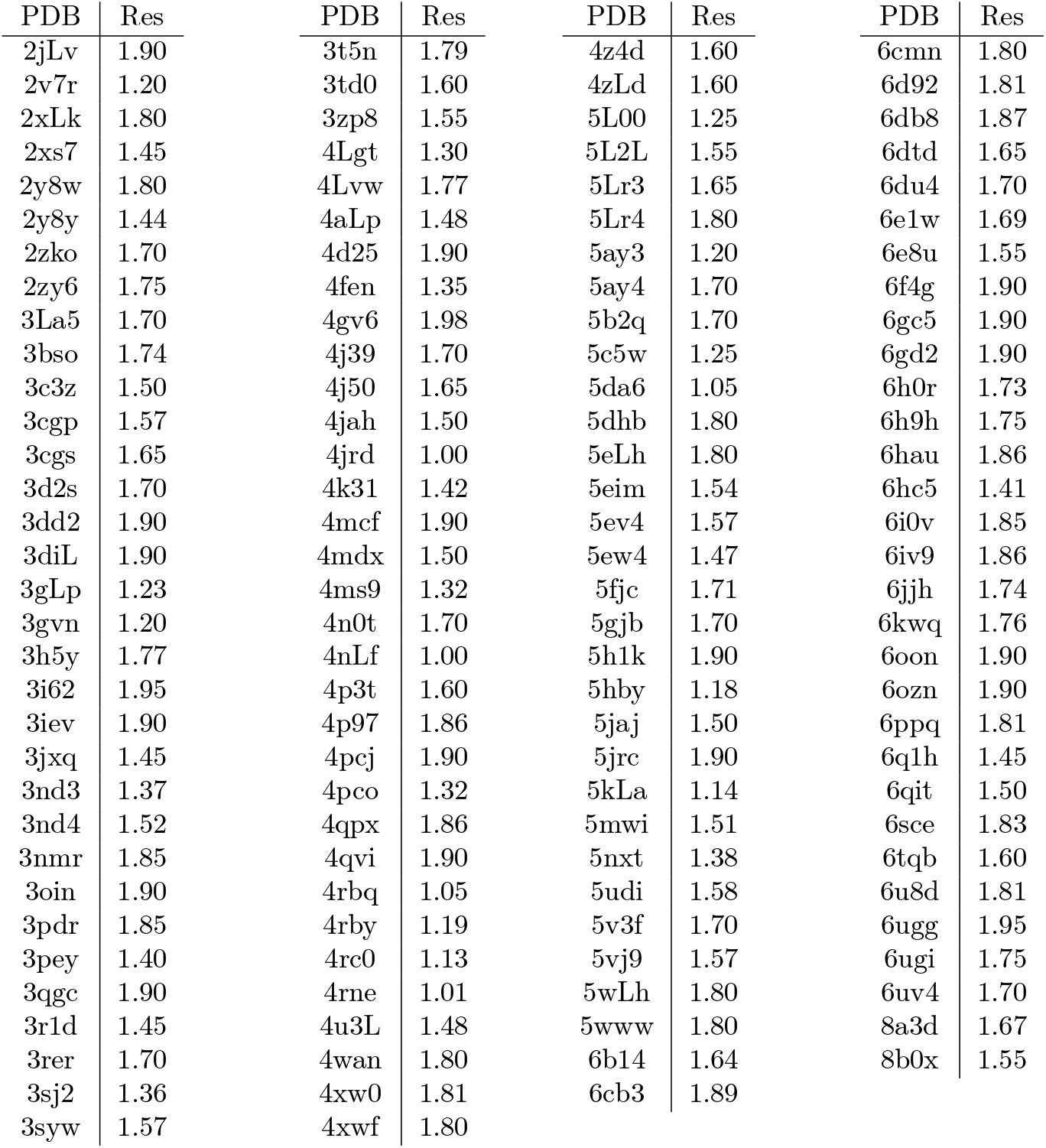
PDB IDs (sorted in alphabetical order) and resolutions of the different measurements of the HD data set (introduced in Section 2.1 in the main text)

**Figure S1:**
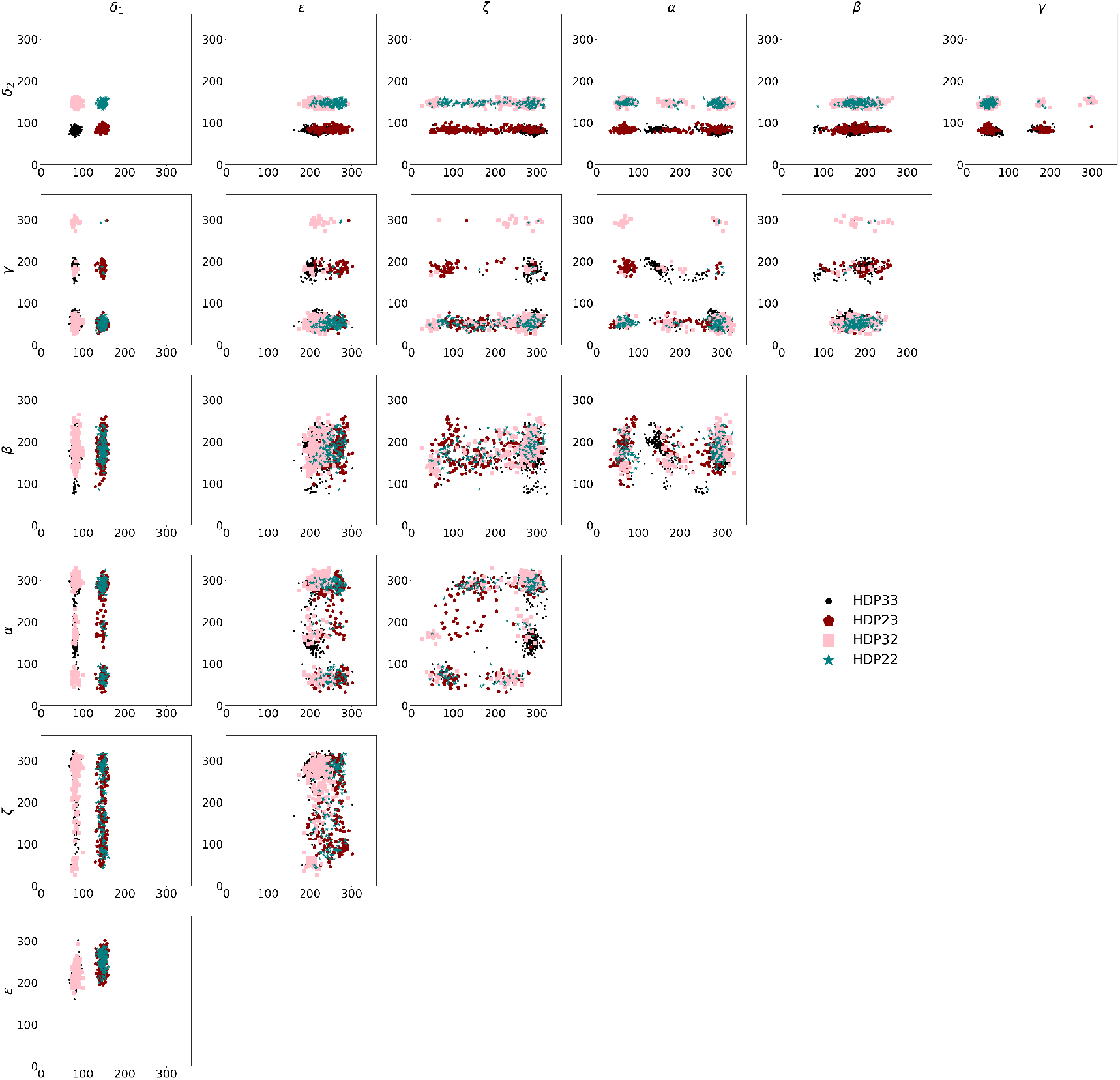
The HD representation of the high resolution training data set, resolved in pucker-pair (introduced in Section 2.1 in the main text), represented by scatter plots of all two-dimensional dihedral angle pairs.

**Figure S2:**
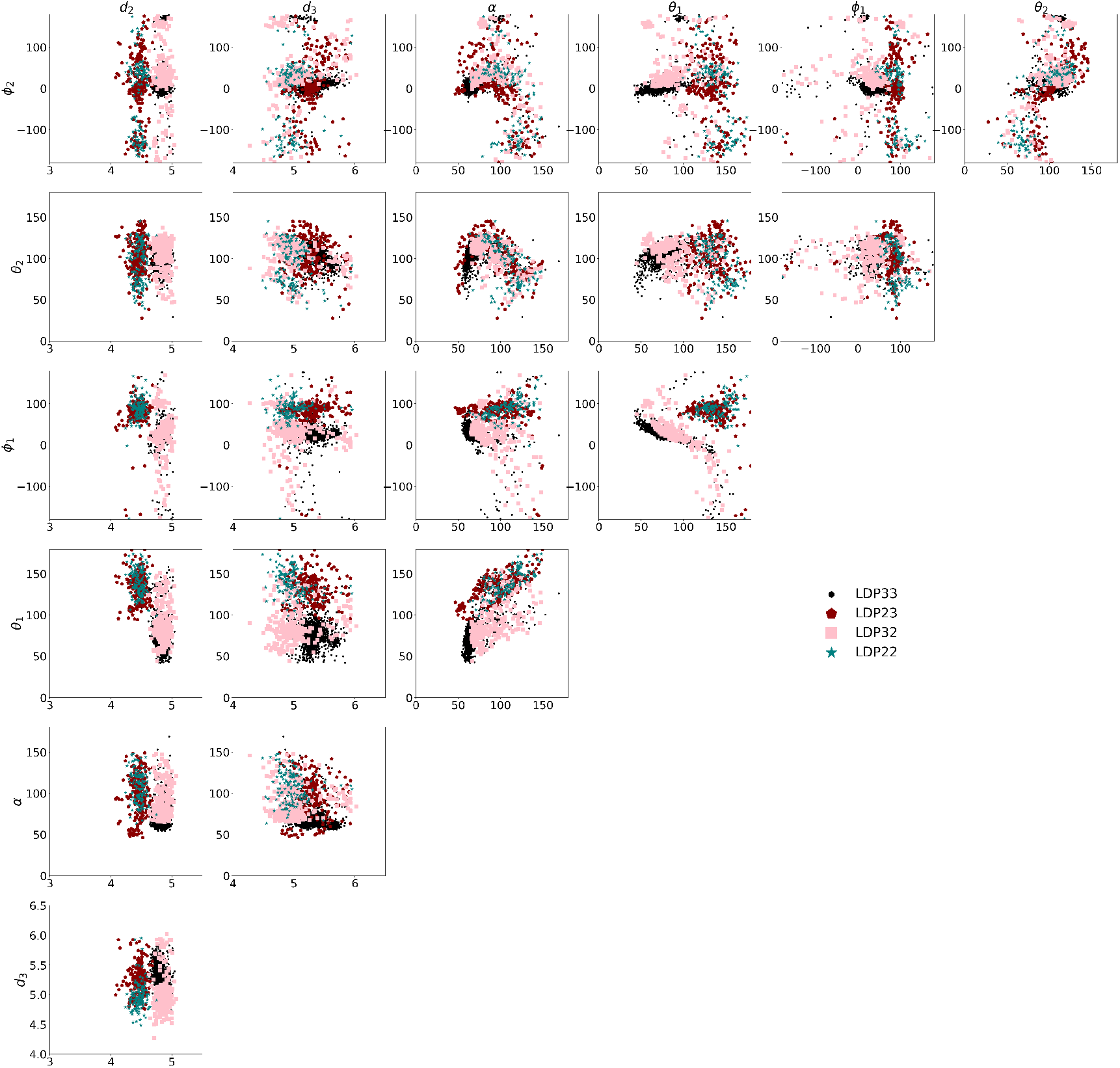
The LD representation of the high resolution training data set, resolved in pucker-pair (introduced in Section 2.1 in the main text), represented by scatter plots of all combinations of parameters (described in Section 2.3.1 in the main text).

### B Low detail test data sets

#### B.1 LDT test data

Table S2 lists the suites contained in the LDT test data set (introduced in Section 2.1). In Figure S3 the four pucker-pair sets of LDT are plotted.

**Table S2:**
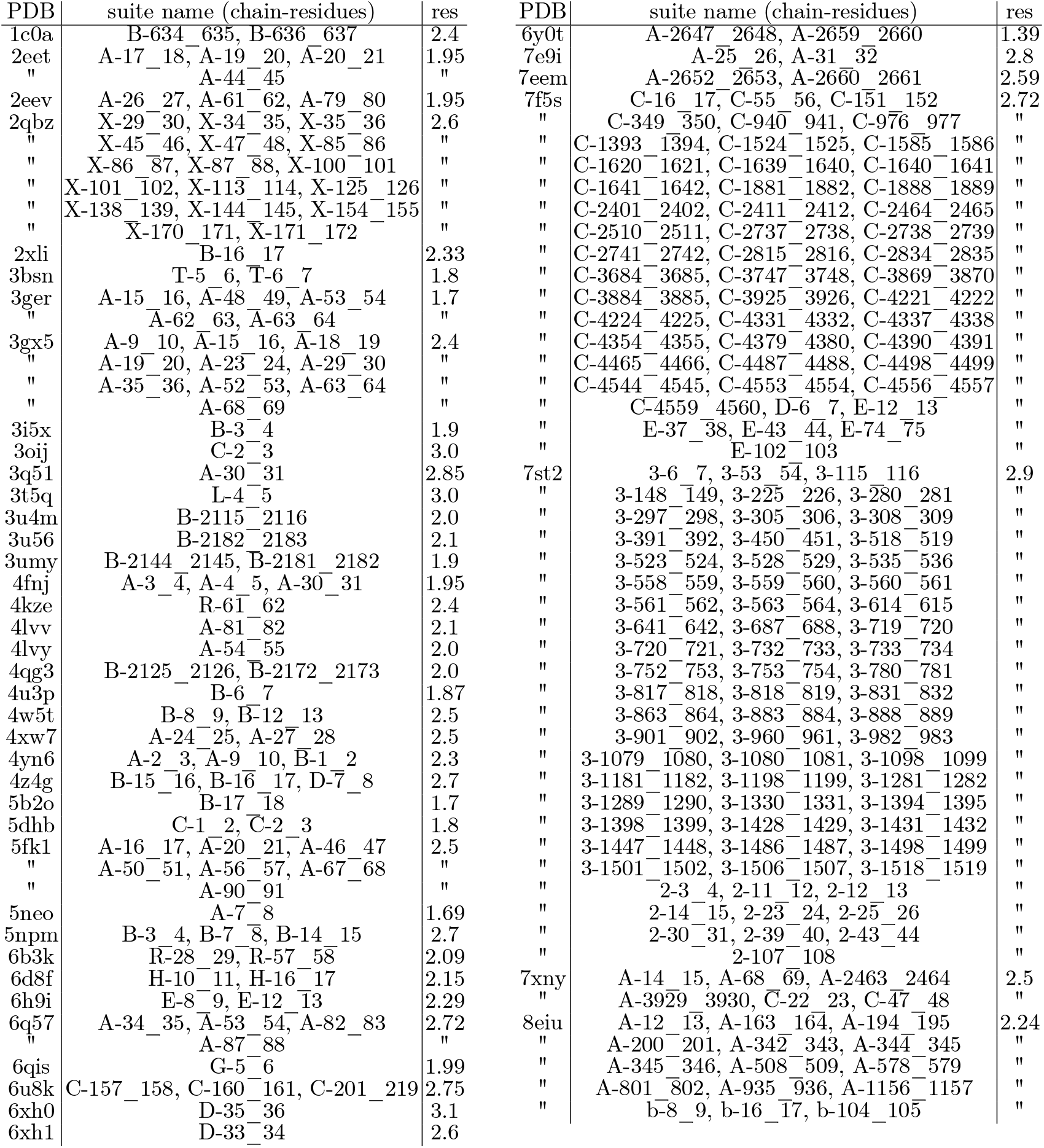
LD data set from Section 2.1 in the main text. The PDB ID, the name of the chain and residue numbers in the suite and the resolution of the respective measurement are listed in alphabetical order.

**Figure S3:**
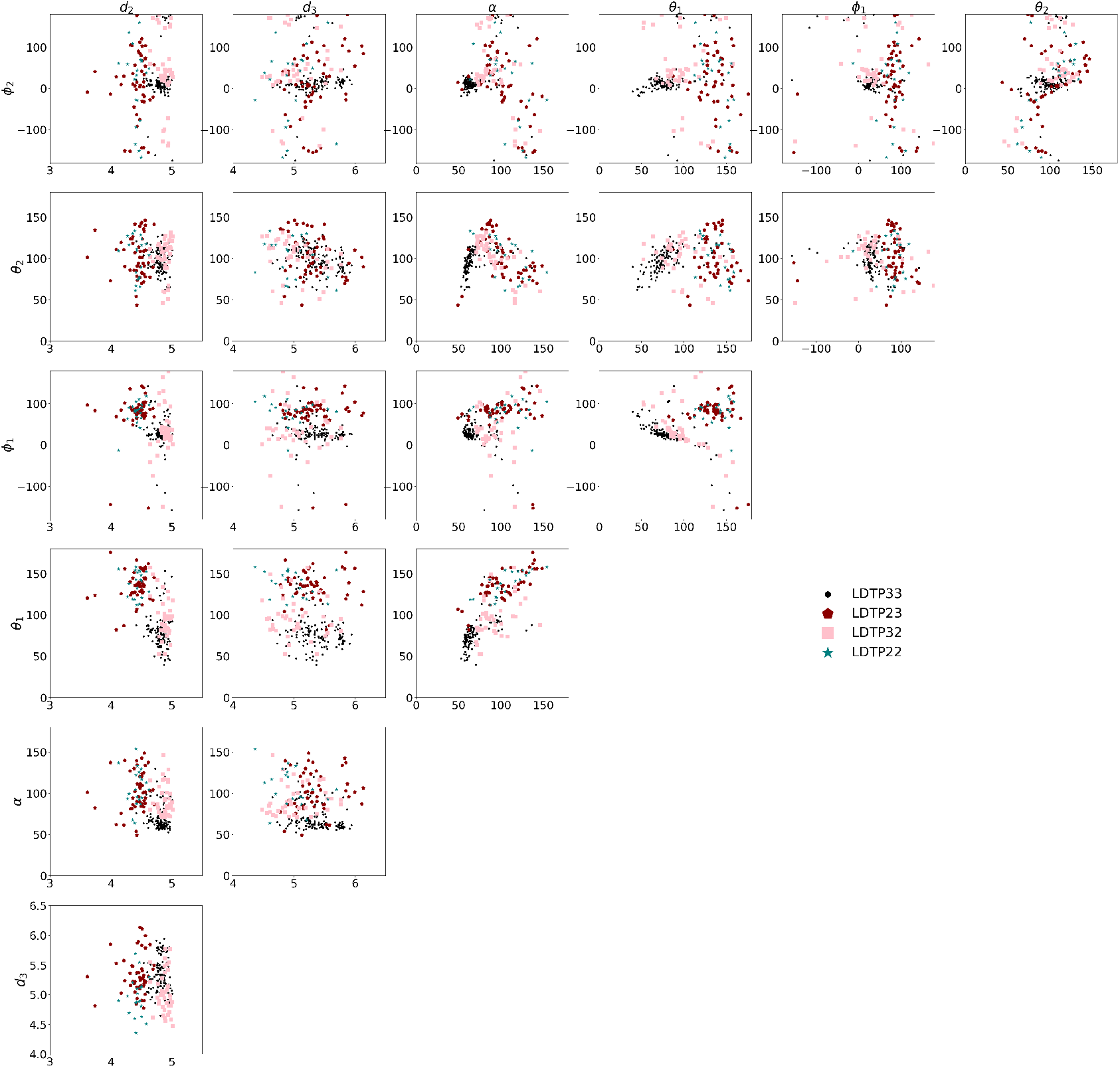
The low detail representation of the LDT data set, resolved in pucker-pair (introduced in Section 2.1 in the main text), represented by scatter plots of all combinations of parameters (described in Section 2.3.1 in the main text).

#### B.2 8b0x test data

Table S3 lists the suites contained in the 8b0x test data set (introduced in Section 2.1). In Figure S4 the four pucker-pair sets of 8b0x are plotted.

**Table S3:** LD data set from Section 2.1 in the main text. The PDB ID, the name of the chain and residue numbers in the suite and the resolution of the respective measurement are listed in alphabetical order.

| PDB | suite name (chain-residues) | res | PDB | suite name (chain-residues) | res |
| --- | --- | --- | --- | --- | --- |
| 7st2 | 1-29_30, 1-45_46, 1-57_58 | 2.9 | 8eiu | a-44_45, a-60_61, a-130_131 | 2.24 |
| " | 1-83_84, 1-95_96, 1-99_100 | " | " | a-537_538, a-864_865, a-1204_1205 | " |
| " | 1-109_110, 1-148_149, 1-198_199 | " | " | a-1240_1241, a-1625_1626, a-1749_1750 | " |
| " | 1-203_204, 1-218_219, 1-219_220 | " | " | a-1790_1791, a-1873_1874, a-1964_1965 | " |
| " | 1-227_228, 1-228_229, 1-229_230 | " | " | a-2079_2080, a-2317_2318, a-2320_2321 | " |
| " | 1-240_241, 1-251_252, 1-254_255 | " | " | a-2343_2344, a-2433_2434, a-2446_2447 | " |
| " | 1-264_265, 1-265_266, 1-293_294 | " | " | a-2474_2475, a-2534_2535 | " |
| " | 1-321_322, 1-322_323, 1-333_334 | " |  |  |  |
| " | 1-370_371, 1-371_372, 1-382_383 | " |  |  |  |
| " | 1-394_395, 1-395_396, 1-402_403 | " |  |  |  |
| " | 1-423_424, 1-442_443, 1-464_465 | " |  |  |  |
| " | 1-466_467, 1-474_475, 1-476_477 | " |  |  |  |
| " | 1-499_500, 1-500_501, 1-516_517 | " |  |  |  |
| " | 1-528_529, 1-561_562, 1-567_568 | " |  |  |  |
| " | 1-621_622, 1-633_634, 1-669_670 | " |  |  |  |
| " | 1-684_685, 1-694_695, 1-788_789 | " |  |  |  |
| " | 1-799_800, 1-818_819, 1-829_830 | " |  |  |  |
| " | 1-857_858, 1-865_866, 1-868_869 | " |  |  |  |
| " | 1-906_907, 1-940_941, 1-972_973 | " |  |  |  |
| " | 1-988_989, 1-998_999, 1-1002_1003 | " |  |  |  |
| " | 1-1004_1005, 1-1020_1021, 1-1024_1025 | " |  |  |  |
| " | 1-1129_1130, 1-1130_1131, 1-1138_1139 | " |  |  |  |
| " | 1-1154_1155, 1-1185_1186, 1-1203_1204 | " |  |  |  |
| " | 1-1205_1206, 1-1210_1211, 1-1211_1212 | " |  |  |  |
| " | 1-1235_1236, 1-1246_1247, 1-1247_1248 | " |  |  |  |
| " | 1-1254_1255, 1-1264_1265, 1-1274_1275 | " |  |  |  |
| " | 1-1344_1345, 1-1358_1359, 1-1367_1368 | " |  |  |  |
| " | 1-1375_1376, 1-1394_1395, 1-1418_1419 | " |  |  |  |
| " | 1-1426_1427, 1-1436_1437, 1-1457_1458 | " |  |  |  |
| " | 1-1477_1478, 1-1503_1504, 1-1521_1522 | " |  |  |  |
| " | 1-1553_1554, 1-1554_1555, 1-1557_1558 | " |  |  |  |
| " | 1-1558_1559, 1-1559_1560, 1-1626_1627 | " |  |  |  |
| " | 1-1633_1634, 1-1638_1639, 1-1653_1654 | " |  |  |  |
| " | 1-1668_1669, 1-1671_1672, 1-1693_1694 | " |  |  |  |
| " | 1-1697_1698, 1-1756_1757, 1-1757_1758 | " |  |  |  |
| " | 1-1775_1776, 1-1779_1780, 1-1785_1786 | " |  |  |  |
| " | 1-1818_1819, 1-1856_1857, 1-1883_1884 | " |  |  |  |
| " | 1-1900_1901, 1-1902_1903, 1-1926_1927 | " |  |  |  |
| " | 1-1935_1936, 1-1965_1966, 1-1979_1980 | " |  |  |  |
| " | 1-1980_1981, 1-1995_1996, 1-1996_1997 | " |  |  |  |
| " | 1-2021_2022, 1-2048_2049, 1-2051_2052 | " |  |  |  |
| " | 1-2092_2093, 1-2202_2203, 1-2242_2243 | " |  |  |  |
| " | 1-2249_2250, 1-2278_2279, 1-2296_2297 | " |  |  |  |
| " | 1-2326_2327, 1-2330_2331, 1-2344_2345 | " |  |  |  |
| " | 1-2349_2350, 1-2360_2361, 1-2444_2445 | " |  |  |  |
| " | 1-2473_2474, 1-2475_2476, 1-2479_2480 | " |  |  |  |
| " | 1-2553_2554, 1-2563_2564, 1-2564_2565 | " |  |  |  |
| " | 1-2571_2572, 1-2614_2615, 1-2645_2646 | " |  |  |  |
| " | 1-2715_2716, 1-2731_2732, 1-2732_2733 | " |  |  |  |
| " | 1-2763_2764, 1-2807_2808, 1-2808_2809 | " |  |  |  |
| " | 1-2817_2818, 1-2822_2823, 1-2858_2859 | " |  |  |  |
| " | 1-2866_2867, 1-2867_2868, 1-2871_2872 | " |  |  |  |
| " | 1-2886_2887, 1-2890_2891 | " |  |  |  |

**Figure S4:**
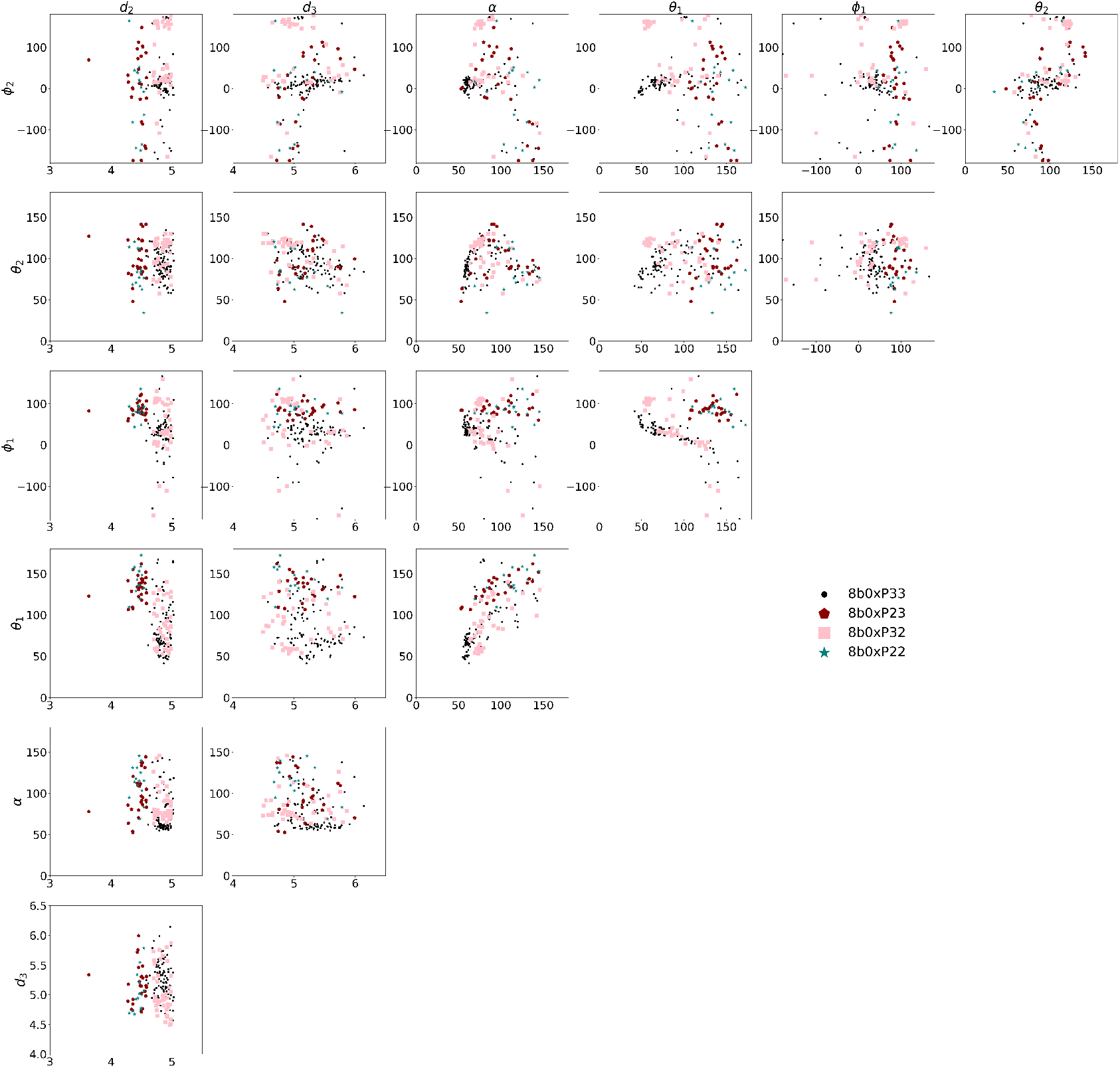
The low detail representation of the 8b0x test data set, resolved in pucker-pair (introduced in Section 2.1 in the main text), represented by scatter plots of all combinations of parameters (described in Section 2.3.1 in the main text).

### C Clustering results of HD and comparison with suitename

This section shows the clustering results of the four datasets HDP33, HDC2C3, HDP32 and HDP22 with the MINT-AGE clustering. The tuning parameters required for MINT-AGE are listed in Table 1 in the main text. The cluster results for the four different data sets are plotted in Figure S5, Figure S6, Figure S7 and Figure S8. In addition, the conformer class was determined for all elements of the four pucker-pair sets with the `phenix.suitename` software. A detailed comparison between the MINT-AGE cluster results and the conformer classification with `phenix.suitename` is provided in Table S4 and in the confusion matrices in Figure S9, Figure S11, Figure S10 and Figure S12.

**Table S4:**
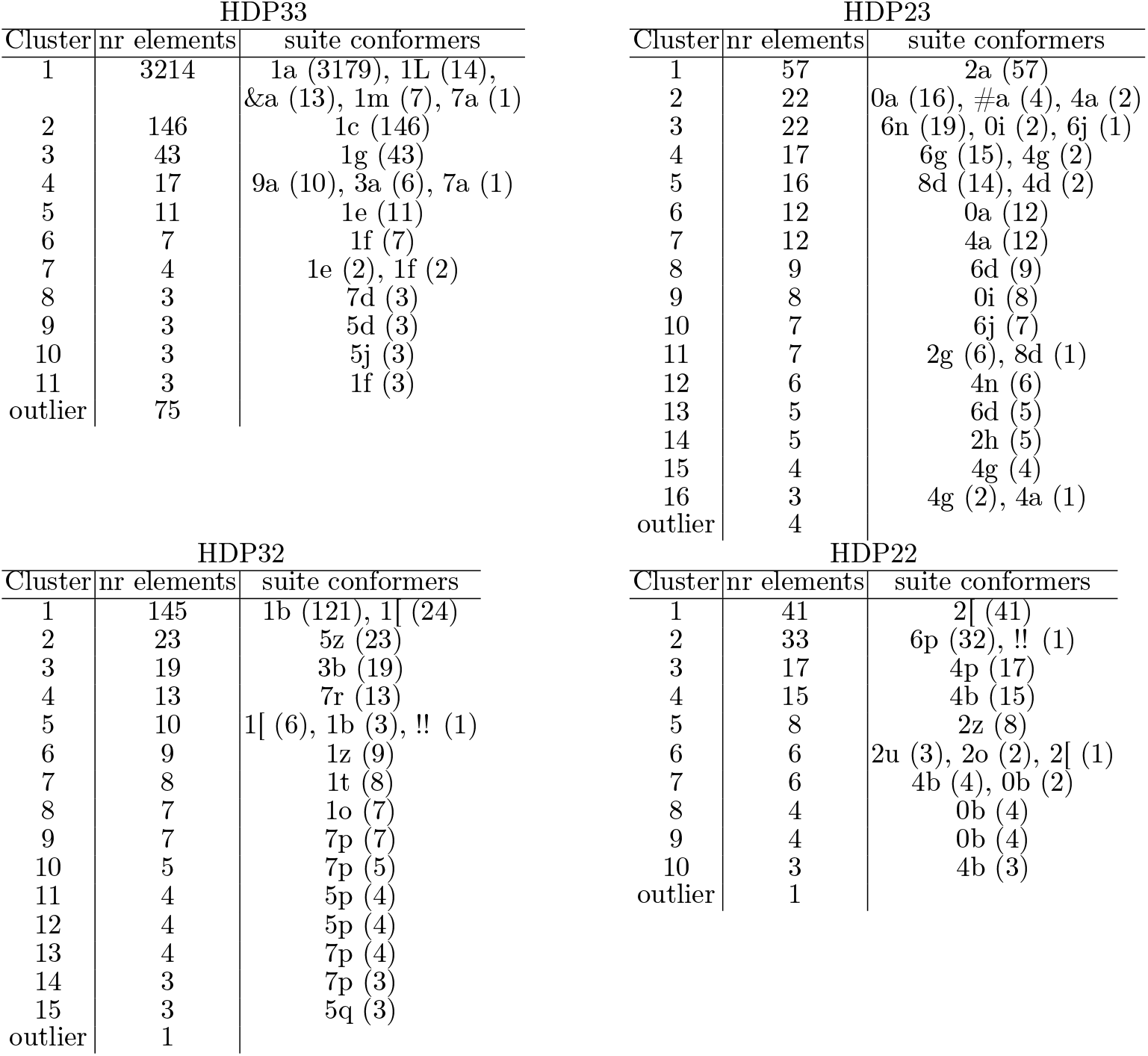
Left and middle column of the tables: MINT-AGE cluster numbers and outliers (left column) with size (middle column) for the different data HDP33, HDP32, HDP23 and HDP22 (introduced in Section 2.1 in the main text). Right column: the corresponding number of elements in the suitename two-digit conformer classes (the name of the class is a number and a letter or a bracket).

**Figure S5:**
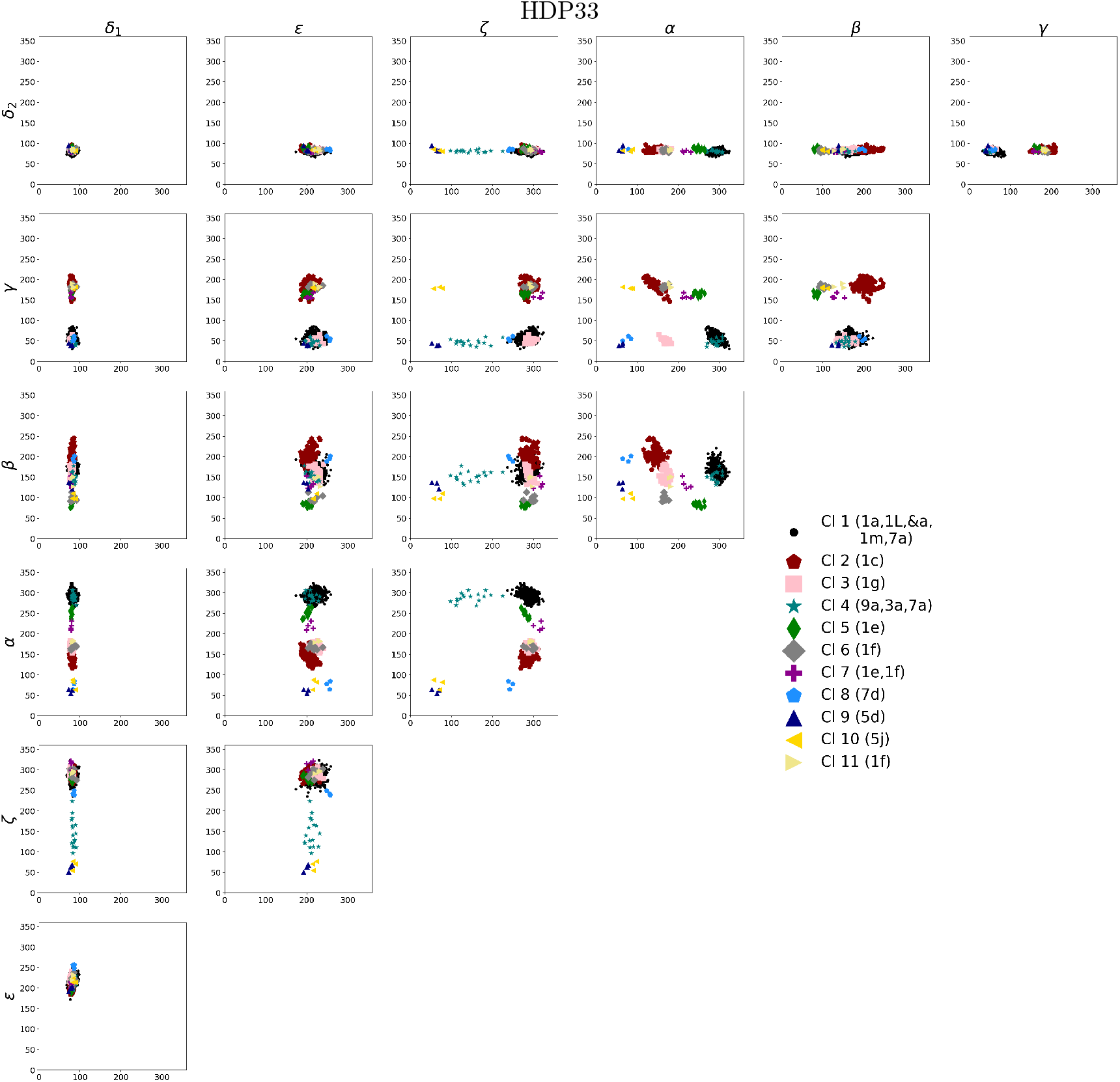
The MINT-AGE clusters (see Section 2.2 in the main text) from the data set HDP33 (introduced in Section 2.1 in the main text) represented by scatter plots of all two-dimensional dihedral angle pairs (given in degrees). See Table S4 for a detailed overview of the clusters and see Figure S9 for a detailed comparison with the conformer classification obtained by the `phenix.suitename` software.

**Figure S6:**
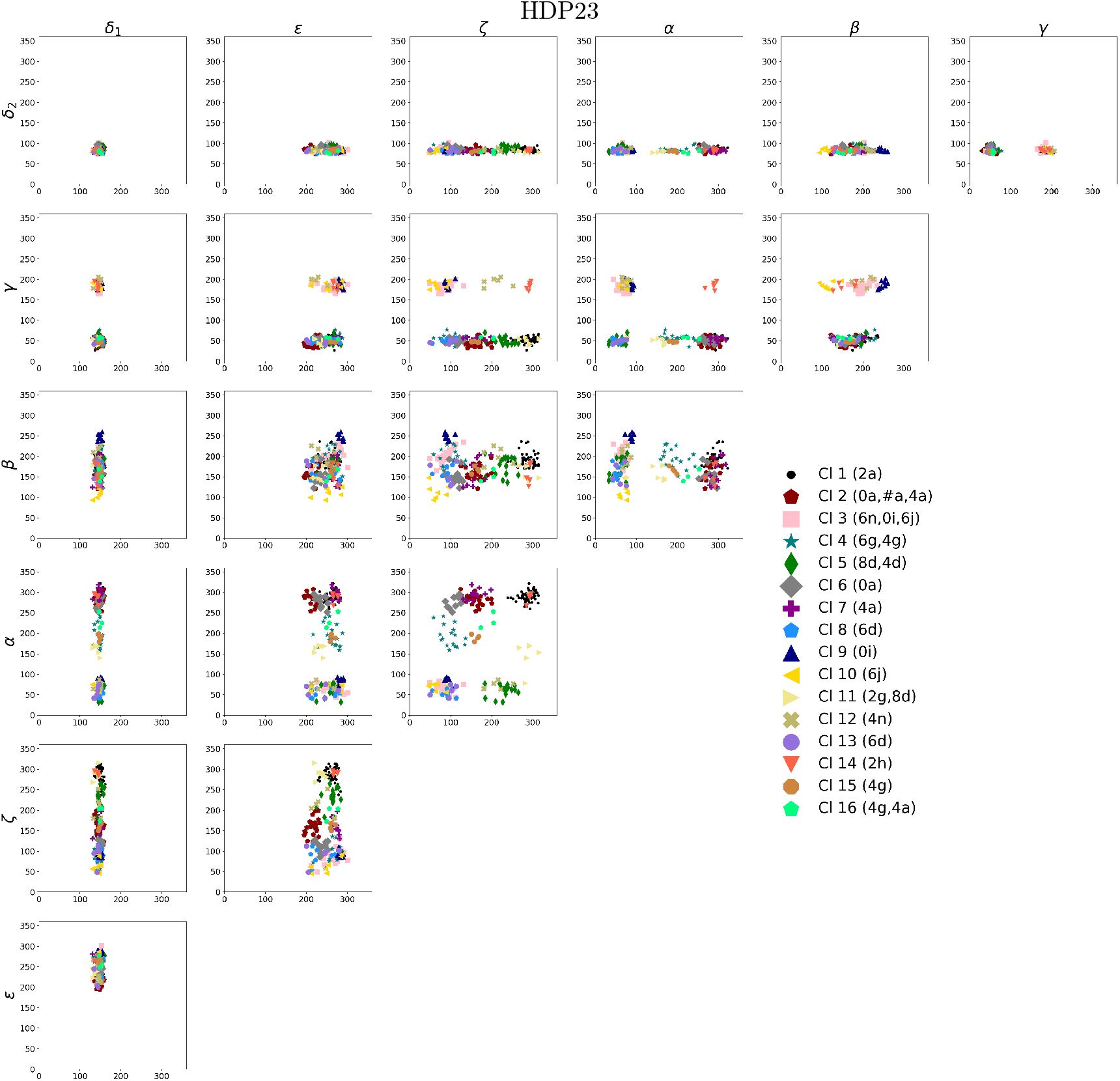
The MINT-AGE clusters (see Section 2.2 in the main text) from the data set HDP23 (introduced in Section 2.1 in the main text) represented by scatter plots of all two-dimensional dihedral angle pairs (given in degrees). See Table S4 for a detailed overview of the clusters and see Figure S11 for a detailed comparison with the conformer classification obtained by the `phenix.suitename` software.

**Figure S7:**
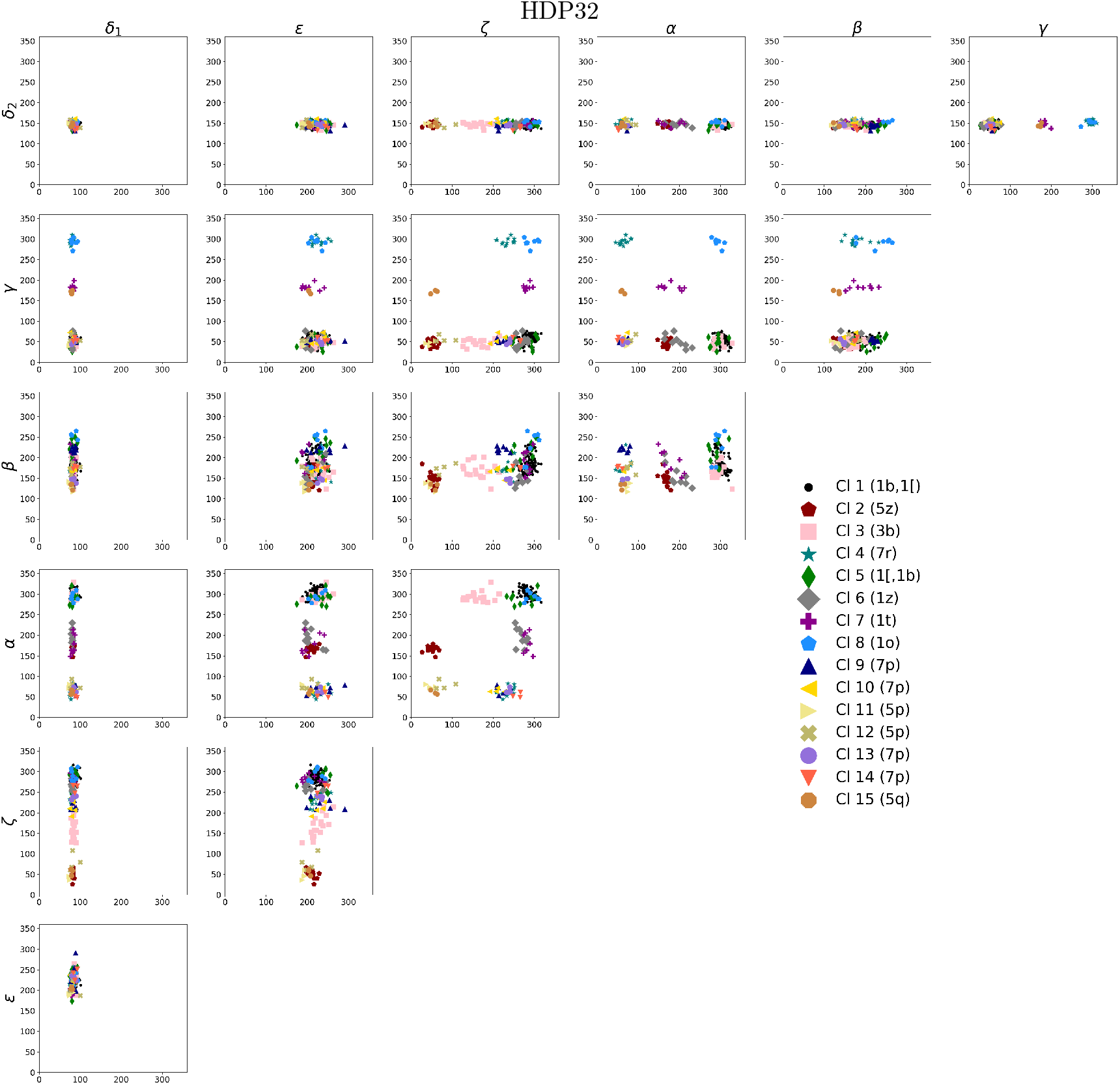
The MINT-AGE clusters (see Section 2.2 in the main text) from the data set HDP32 (introduced in Section 2.1 in the main text) represented by scatter plots of all two-dimensional dihedral angle pairs (given in degrees). See Table S4 for a detailed overview of the clusters and see Figure S10 for a detailed comparison with the conformer classification obtained by the `phenix.suitename` software.

**Figure S8:**
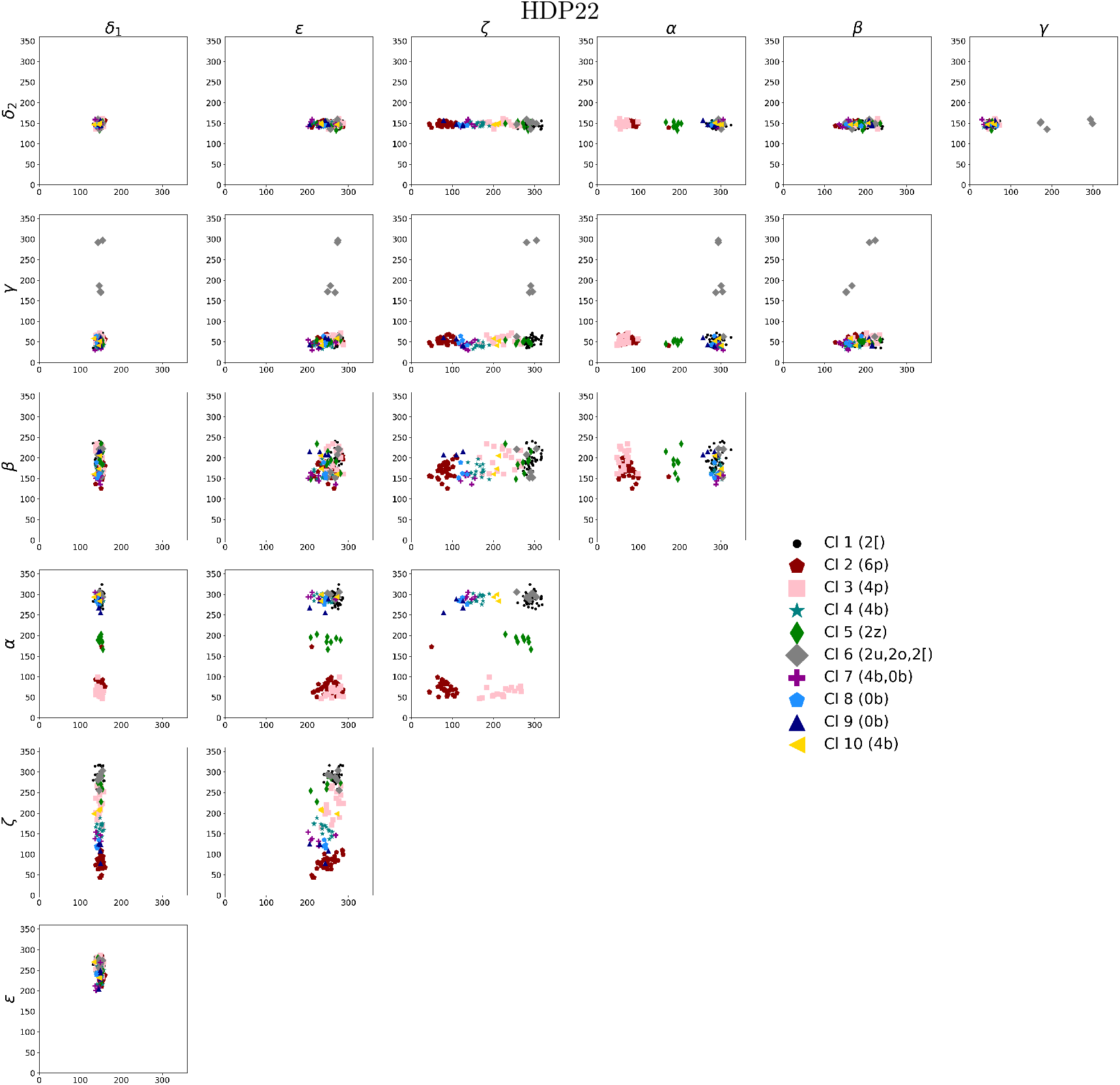
The MINT-AGE clusters (see Section 2.2 in the main text) from the data set HDP22 (introduced in Section 2.1 in the main text) represented by scatter plots of all two-dimensional dihedral angle pairs (given in degrees). See Table S4 for a detailed overview of the clusters and see Figure S11 for a detailed comparison with the conformer classification obtained by the `phenix.suitename` software.

**Figure S9:**
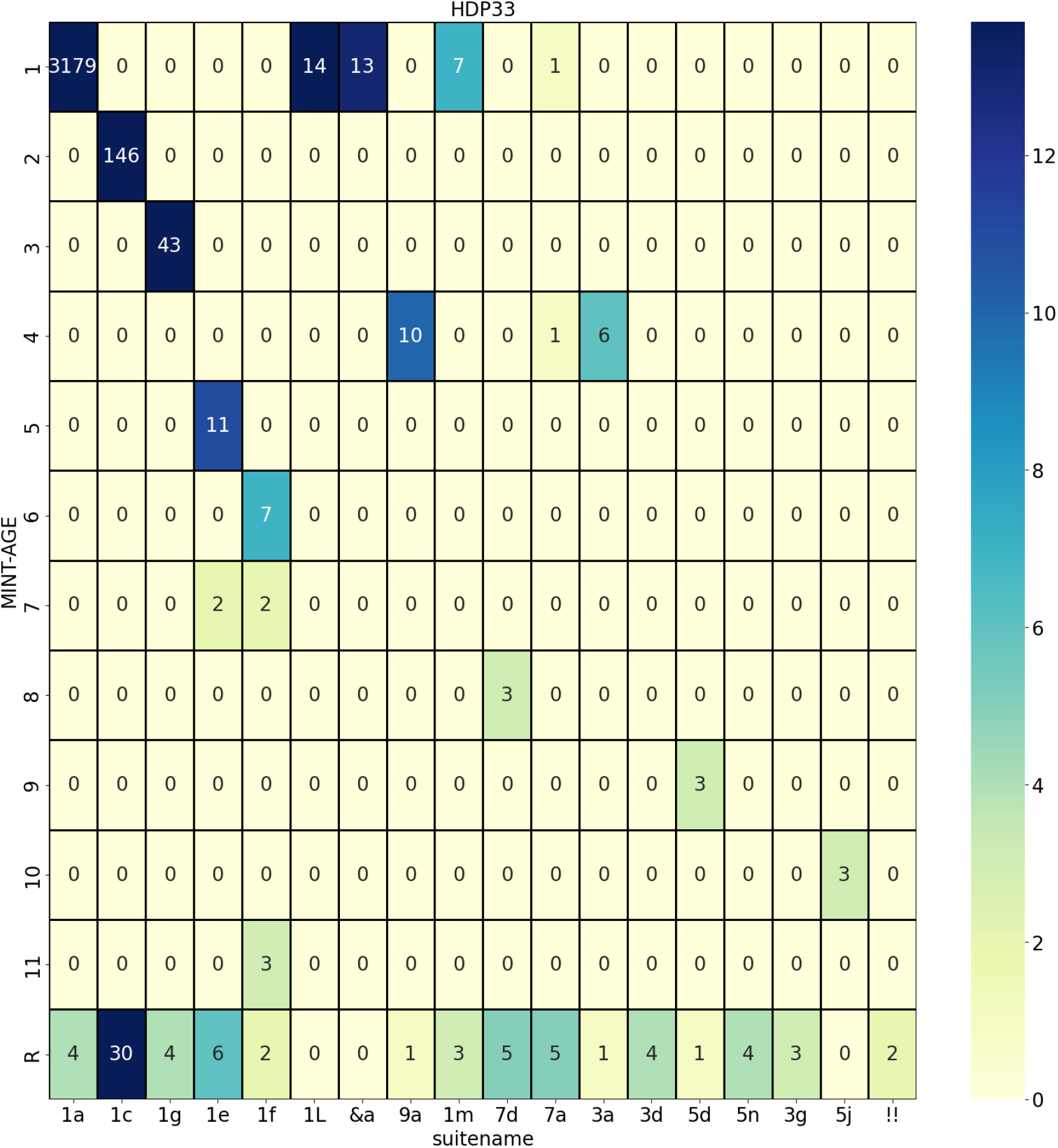
Confusion matrix comparing the MINT-AGE cluster results with suite conformers for HDP33 training data. Every line in the matrix corresponds to a cluster and every column corresponds to a suite conformer. The last line (labeled R for rest) summarizes suites that were not assigned to any cluster by the MINT-AGE algorithm. One can see that most conformers are assigned to a single cluster with overwhelming majority. However, some clusters pool together several conformers.

**Figure S10:**
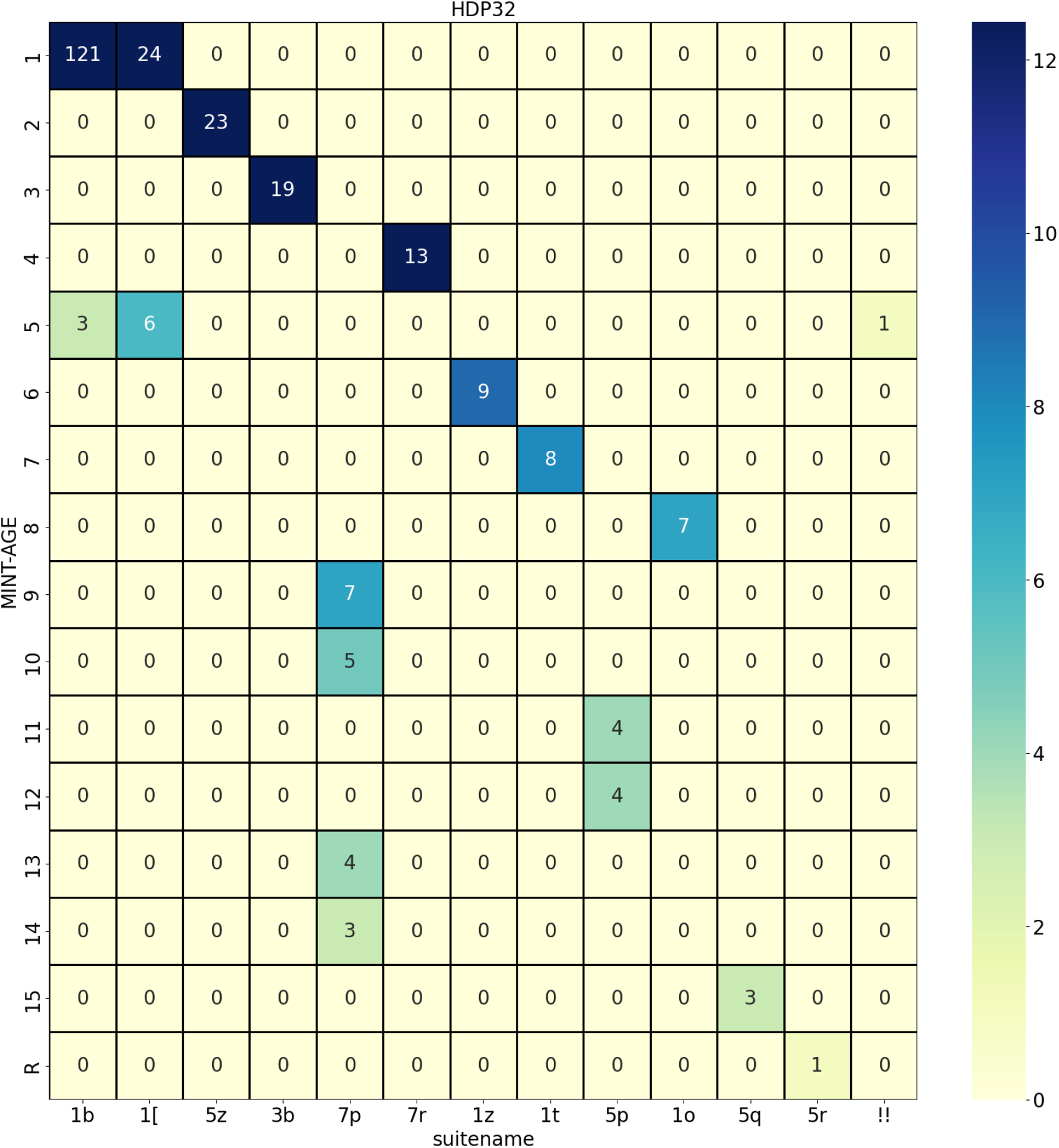
Confusion matrix comparing the MINT-AGE cluster results with suite conformers for HDP32 training data. Every line in the matrix corresponds to a cluster and every column corresponds to a suite conformer. The last line summarizes outliers that were not assigned to any cluster by the MINT-AGE algorithm. Most clusters only comprise a single conformer, except for clusters 1 and 5 which contain both 1b and 1[conformers. Additionally, the 7p and 5p conformers are split up into several cluster.

**Figure S11:**
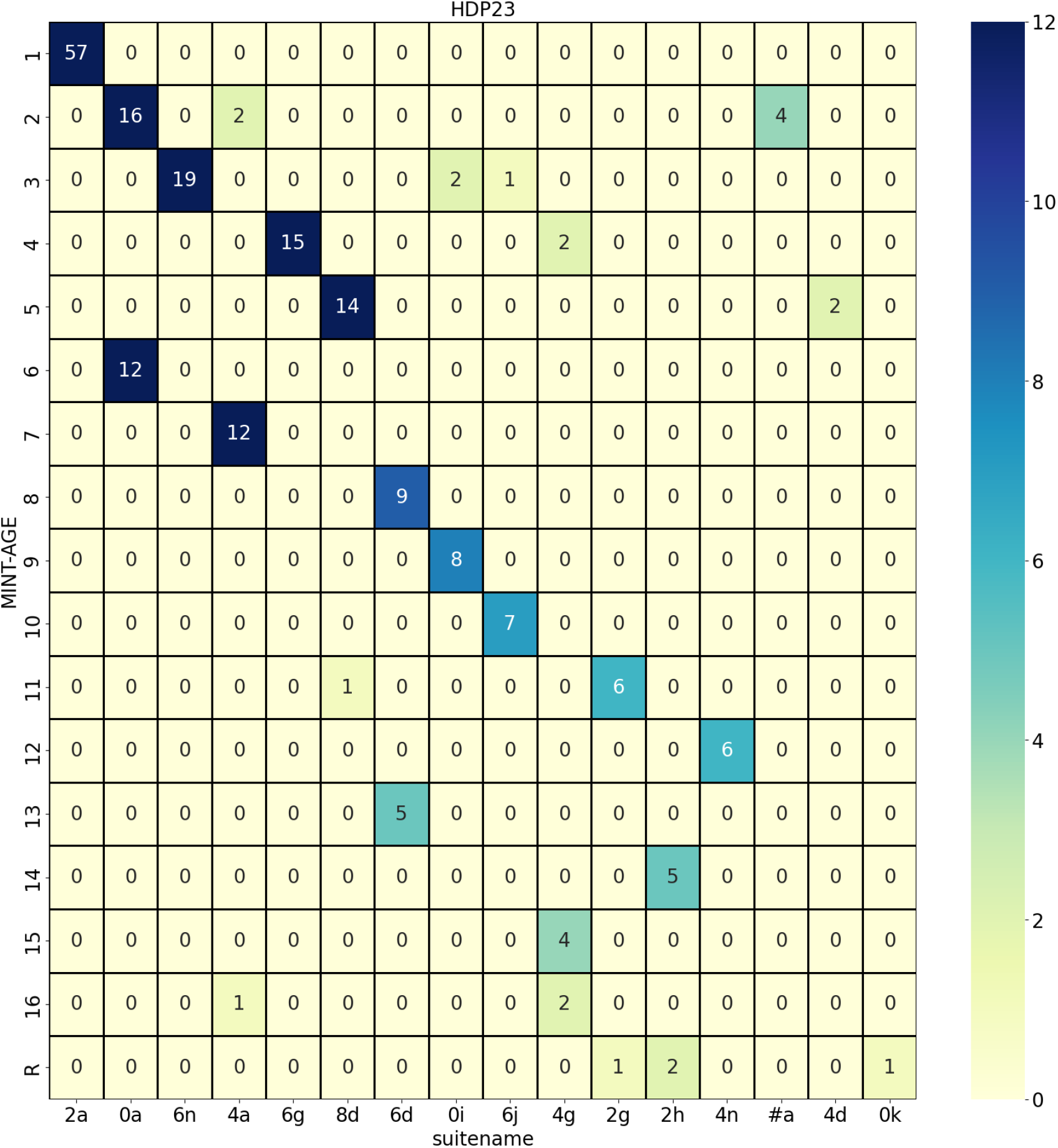
Confusion matrix comparing the MINT-AGE cluster results with suite conformers for HDP23 training data. Every line in the matrix corresponds to a cluster and every column corresponds to a suite conformer. The last line summarizes outliers that were not assigned to any cluster by the MINT-AGE algorithm. Most clusters only comprise a single conformer, except for cluster 2 containing both 0a and #a conformers. Additionally, the 0a and 6d conformers are split up into several cluster.

**Figure S12:**
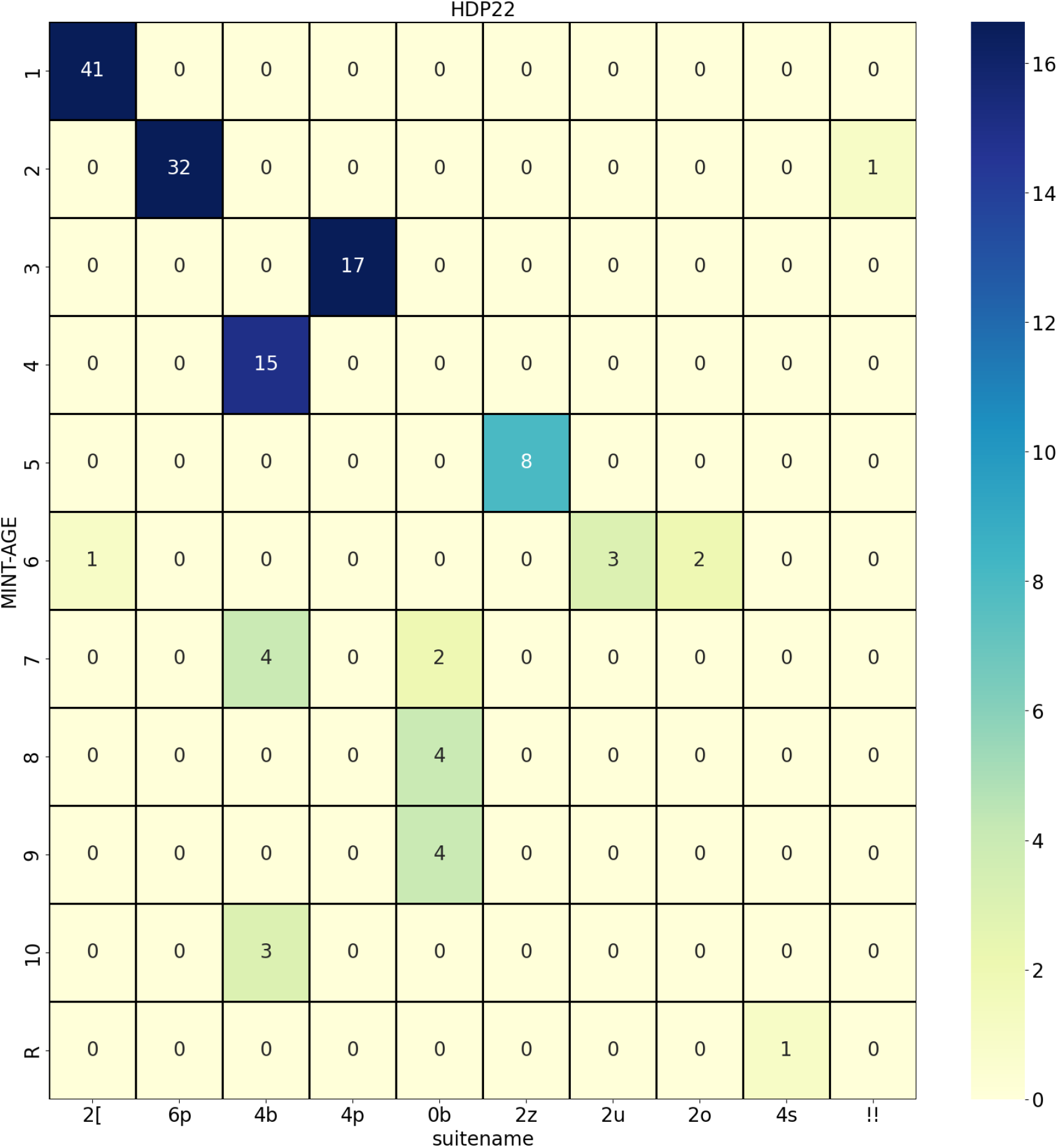
Confusion matrix comparing the MINT-AGE cluster results with suite conformers for HDP22 training data. Every line in the matrix corresponds to a cluster and every column corresponds to a suite conformer. The last line summarizes outliers that were not assigned to any cluster by the MINT-AGE algorithm. While bigger clusters correspond pretty well with abundant conformers, the picture is less clear cut for smaller clusters and rare conformers.

### D Low detail (LD) parameter space

**Figure S13:**
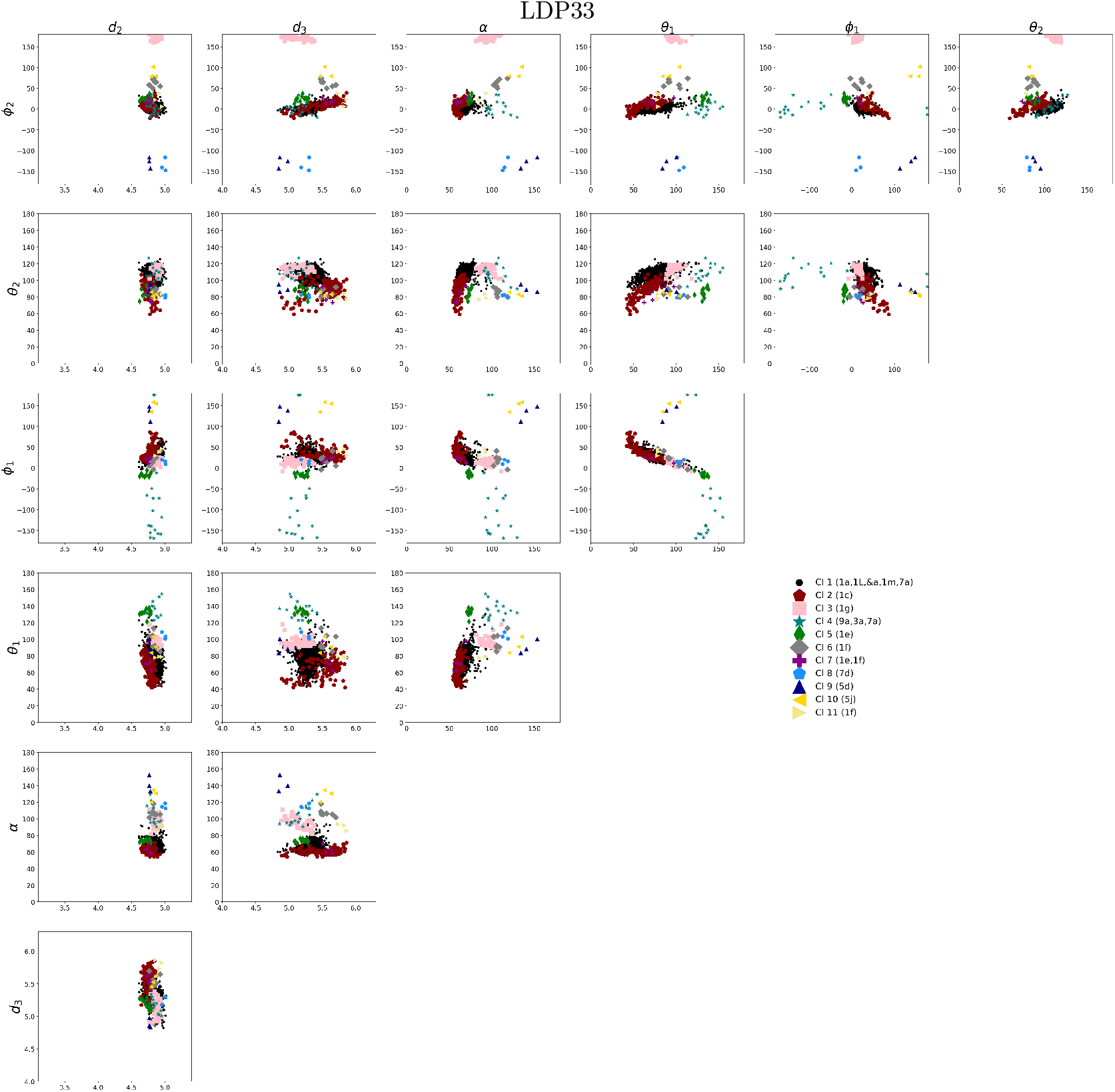
Scatter plot of LDP33 data in low detail parameter space introduced in Section 2.3.1 of the main text.

**Figure S14:**
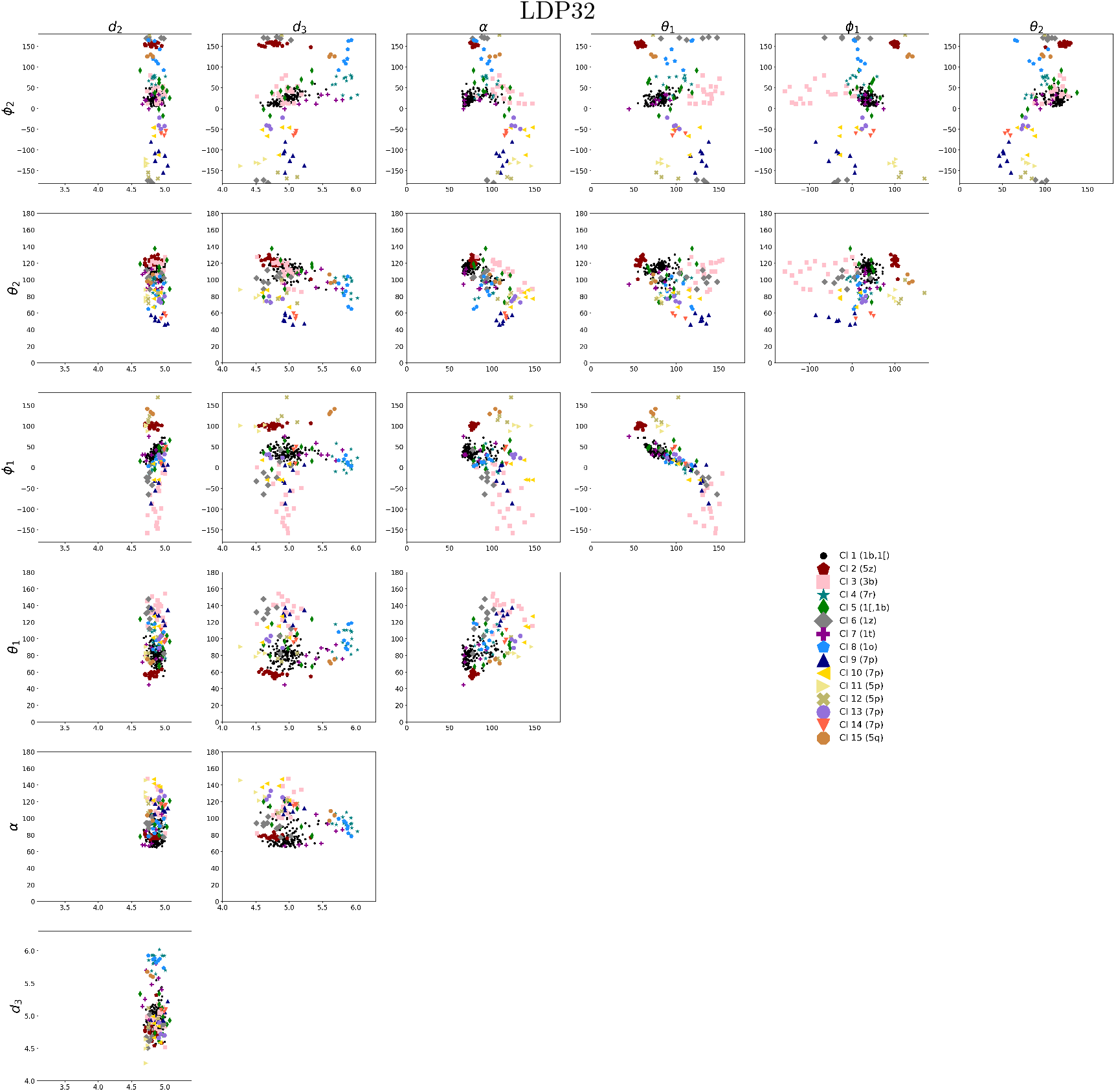
Scatter plot of LDP32 data in low detail parameter space introduced in Section 2.3.1 of the main text.

**Figure S15:**
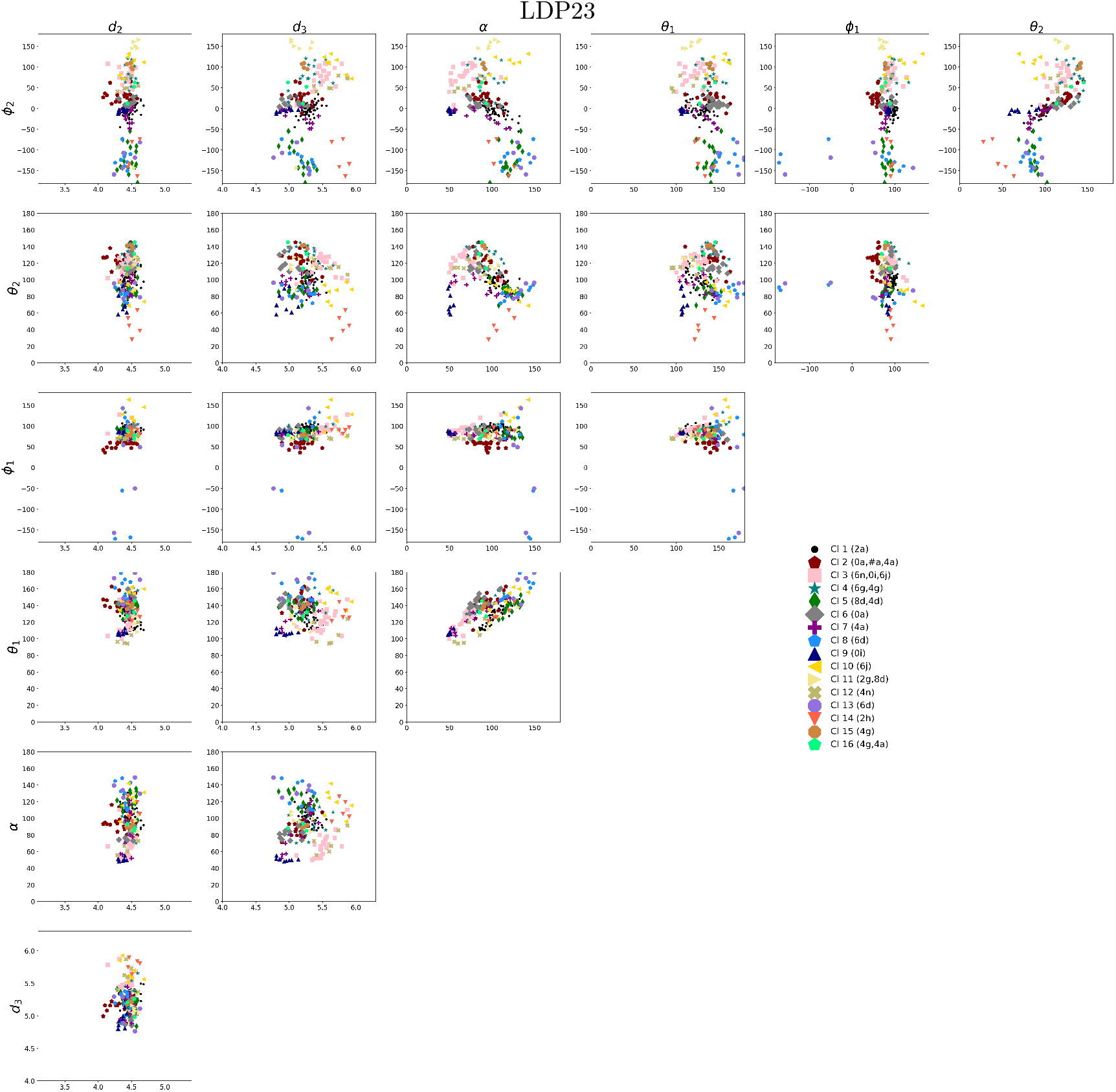
Scatter plot of LDP23 data in low detail parameter space introduced in Section 2.3.1 of the main text.

**Figure S16:**
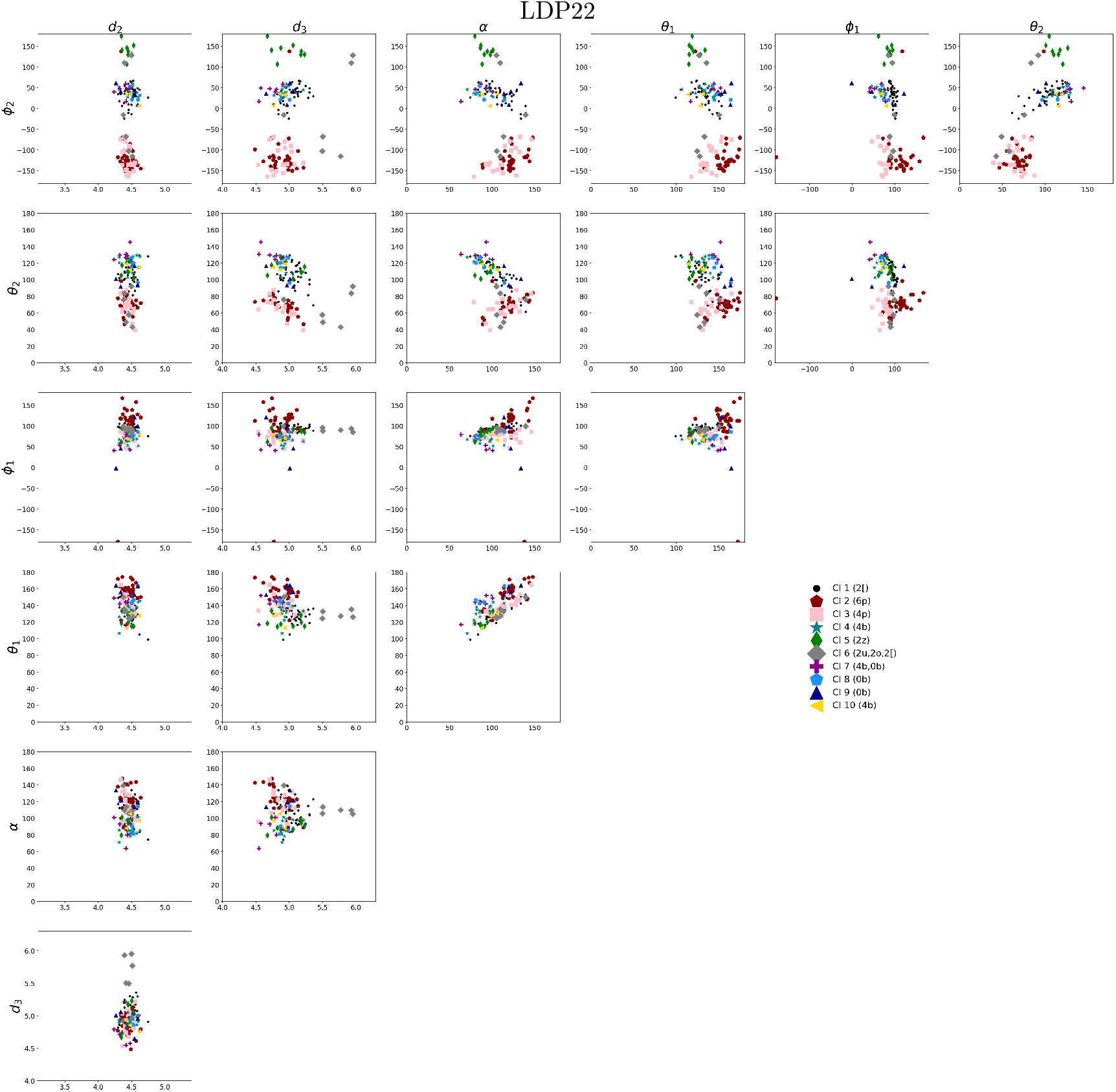
Scatter plot of LDP22 data in low detail parameter space introduced in Section 2.3.1 of the main text.

### E Results RNAprecis

#### E.1 Results for the test data set LDTP33

**Table S5:**
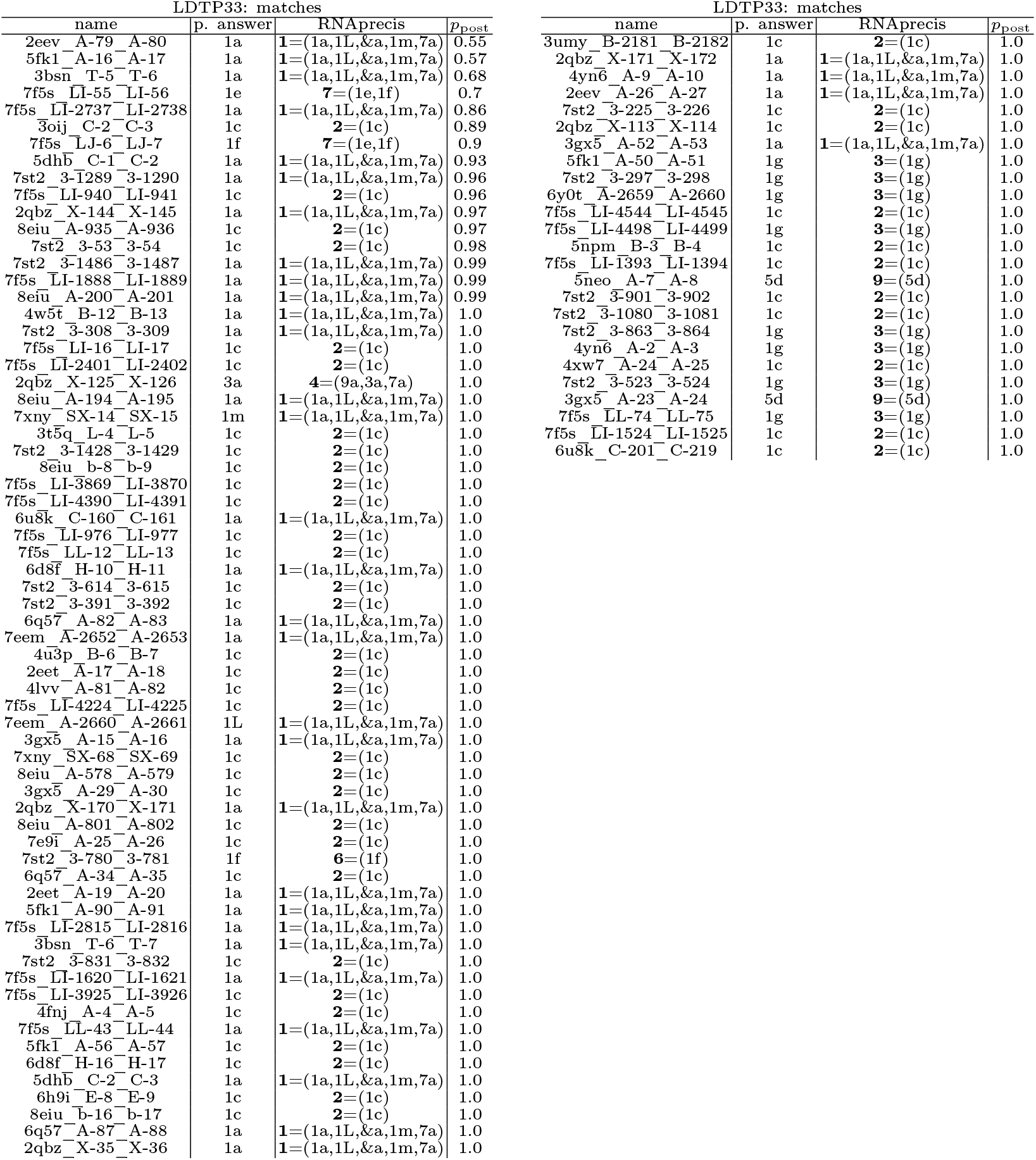
Suites for which the RNAprecis predicted cluster matches the sequence related conformer (blue in the plots) sorted by posterior probability.

**Table S6:**
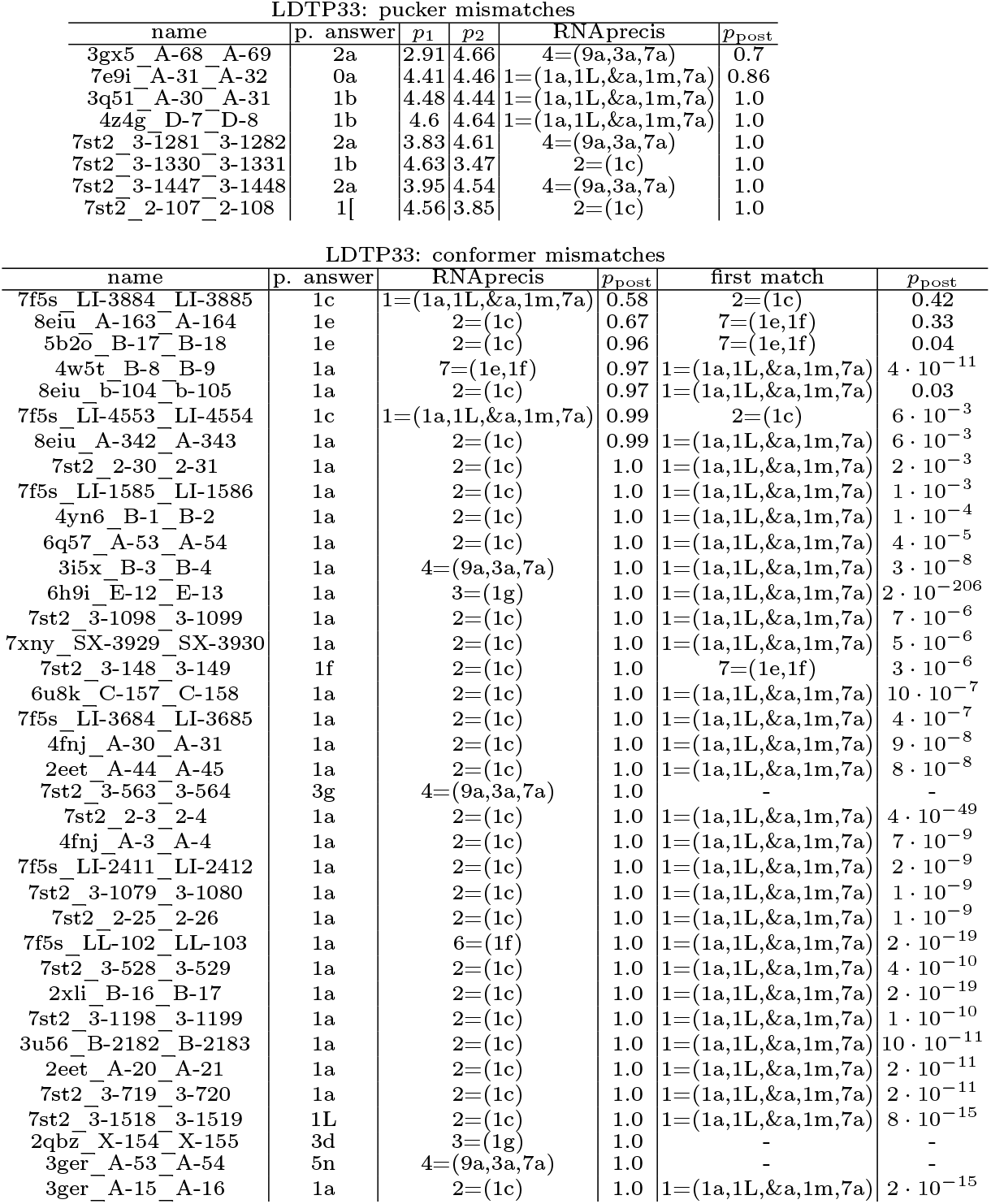
Suites for which the sugar pucker-pair determined by the Pperp criterion do not match the sugar pucker-pair of the sequence related conformers (red in the plots) or the RNAprecis predicted cluster does not match the sequence related conformer (orange in the plots) sorted by posterior probability.

**Figure S17:**
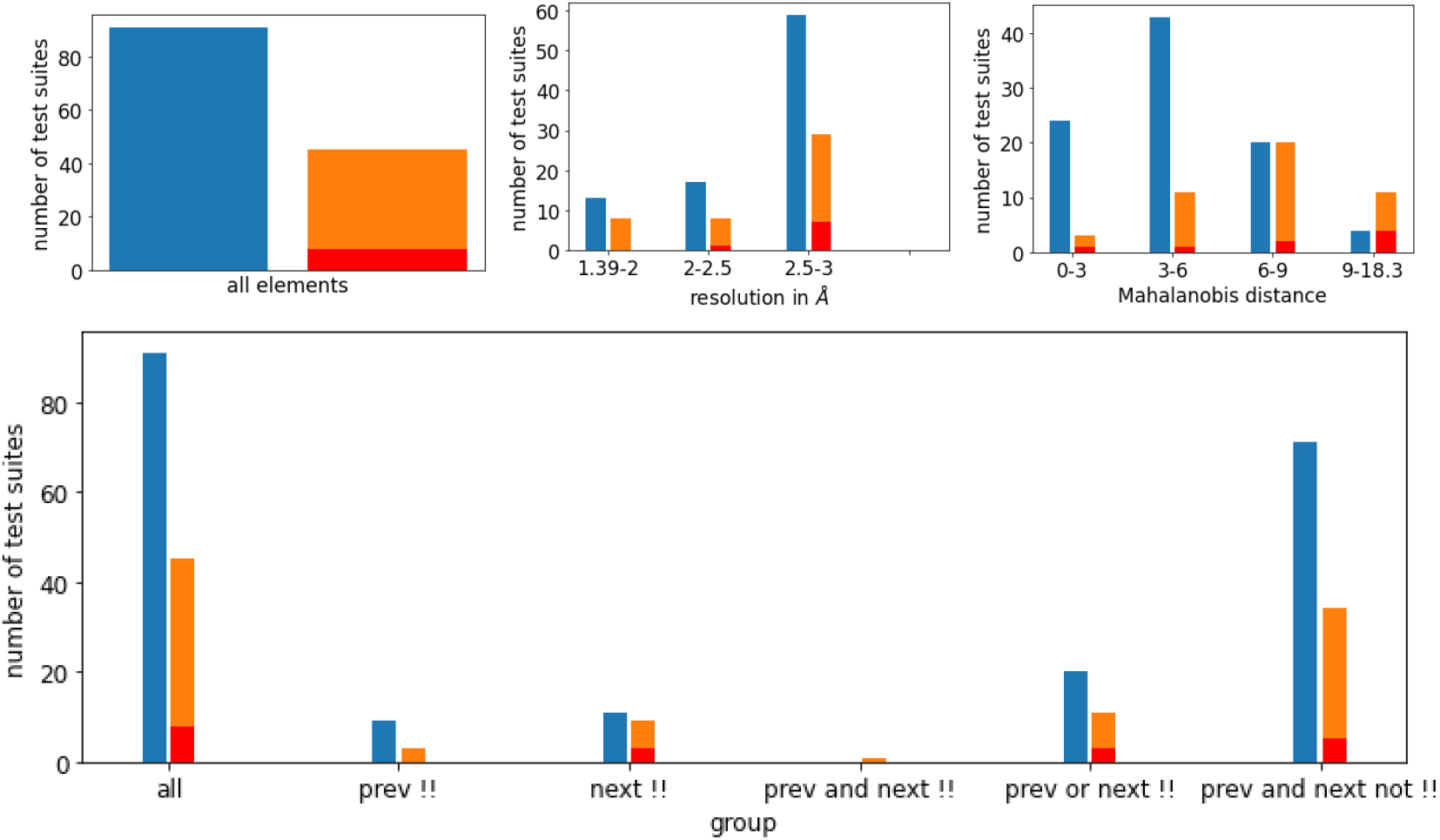
Relative distribution of agreements and disagreements of MINT-AGE cluster prediction with sequence related probable conformer classes for the test set LDTP33 with color scheme (reflecting blue for match, orange for mismatch and red for pucker-pair mismatch) from Section 3.4 in the main text). Top left: overall histogram. Top middle: histogram reflecting varying resolutions of PDB suites. Top right: histogram reflecting varying Mahanalobis distances from the predicted cluster. One can see that with increasing distance the rate of mismatches increases. This indicates that despite its limitations discussed in Remark 2.3 in the main text the distance is useful as an empirical indicator of prediction confidence. Bottom: histogram also reflecting dependence on whether previous, next or both suites have been !! or not. While false predictions are more likely if at least one neighboring suite is a !! suite, roughly half of the total false predictions are from suites without a neighboring !! suite.

#### E.2 Results for the test data set LDTP32

**Table S7:**
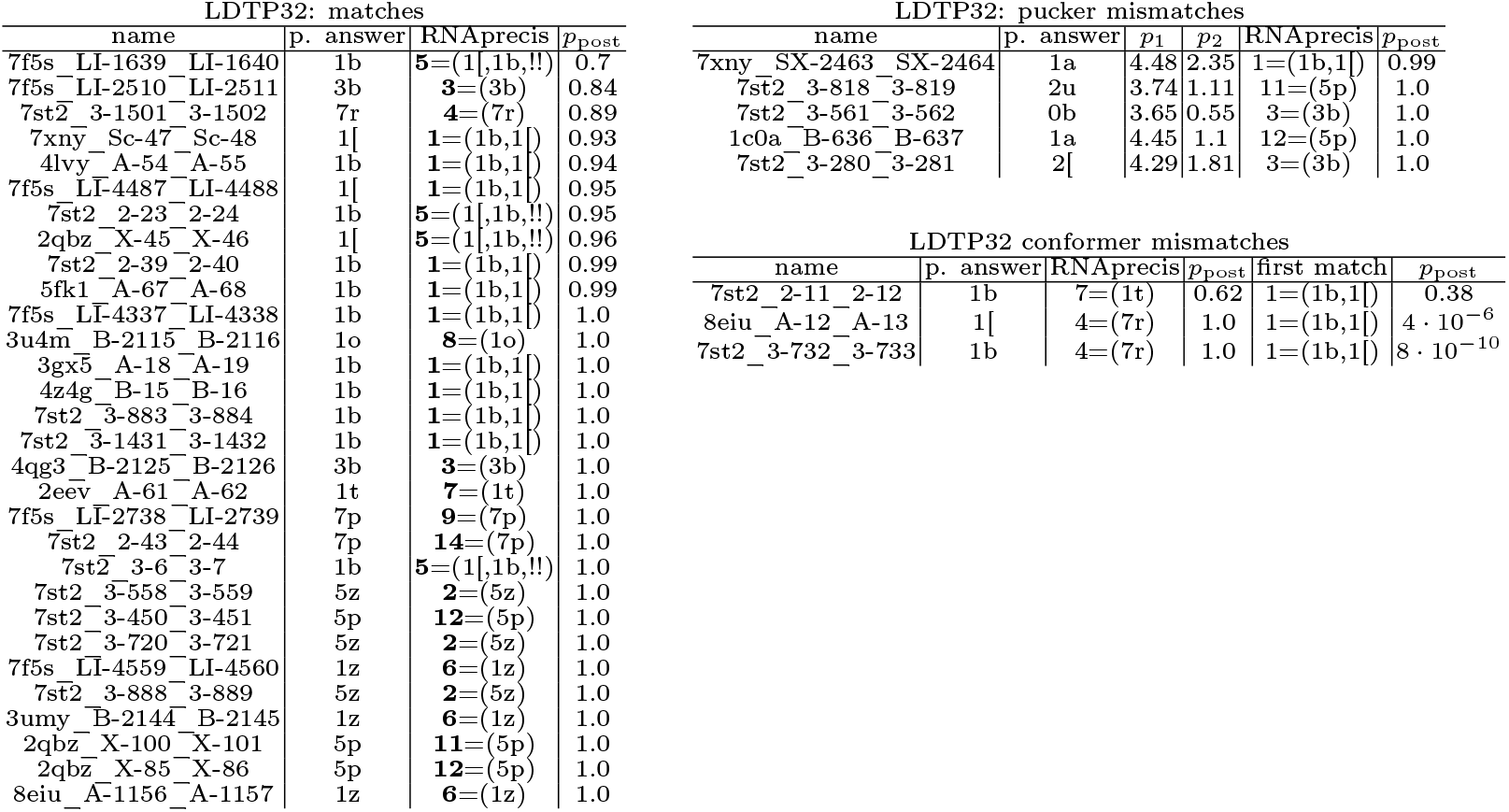
Suites for which the RNAprecis predicted cluster matches the sequence related conformer (blue in the plots), the sugar pucker-pair determined by the Pperp criterion do not match the sugar pucker-pair of the sequence related conformers (red in the plots), or the RNAprecis predicted cluster does not match the sequence related conformer (orange in the plots); each sorted by posterior probability.

**Figure S18:**
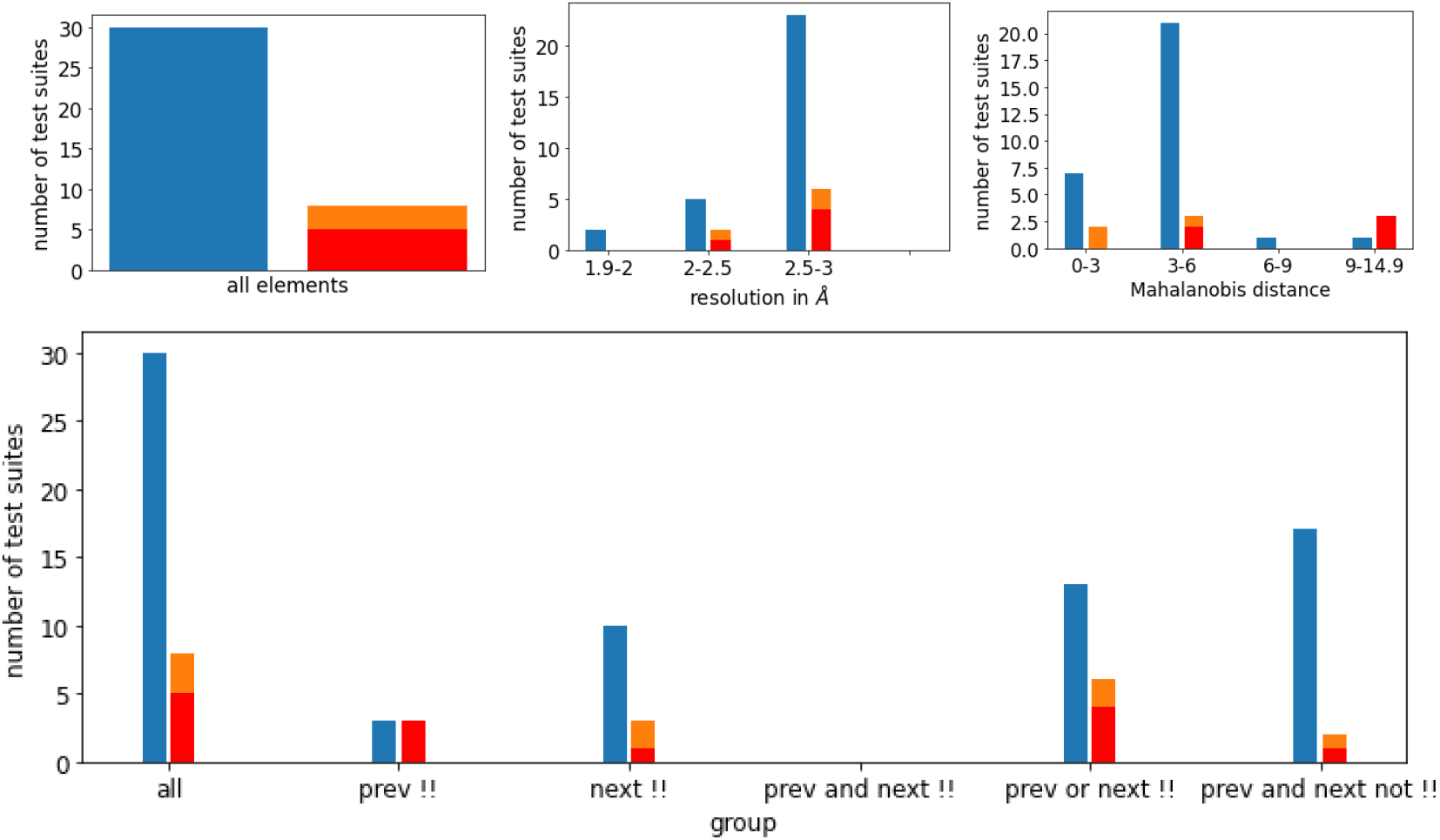
Relative distribution of agreements and disagreements of MINT-AGE cluster prediction with sequence related probable conformer classes for the test set LDT32 with color scheme (reflecting blue for match, orange for mismatch and red for pucker-pair mismatch) from Section 3.4 in the main text). Top left: overall histogram. Top middle: histogram reflecting varying resolutions of PDB suites. Top right: histogram reflecting varying Mahanalobis distances from the predicted cluster. One can see that with increasing distance the rate of mismatches increases. This indicates that despite its limitations discussed in Remark 2.3 in the main text the distance is useful as an empirical indicator of prediction confidence. Bottom: histogram also reflecting dependence on whether previous, next or both suites have been !! or not.

**Figure S19:**
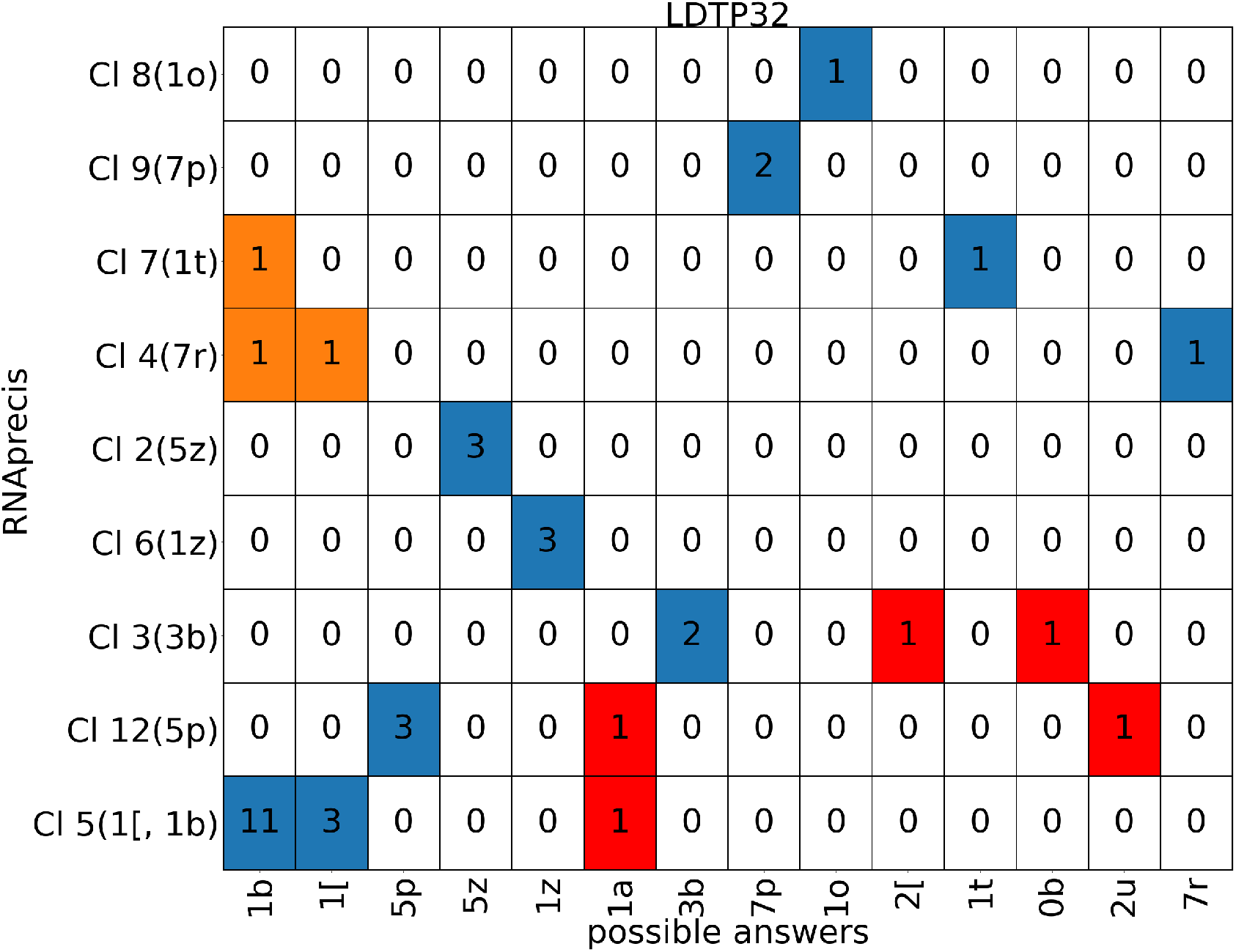
Classification matrix listing counts of probable sequence related conformation classes (horizontal axis) versus MINT-AGE clusters (vertical, with conformer classes in parentheses from MINT-AGE training, see Figure S10) for the low detail training data set of sugar pucker-pair LDP32 with colors (from Section 3.4 in the main text): blue for match, orange for mismatch and red for pucker-pair mismatch. Columns listing multiple suite conformers are from sites solved in different conformations across sequence-related models, and any of the sequence-related conformations were accepted as a matching predictions. As pointed out in Section 3.4, the **1a**,**3g**,**8d** and **3d**,**7p** suites have multiple sequence-related conformations. This means that the counts in these two columns are inflated.

#### E.3 Results for the test data set LDTP23

**Table S8:**
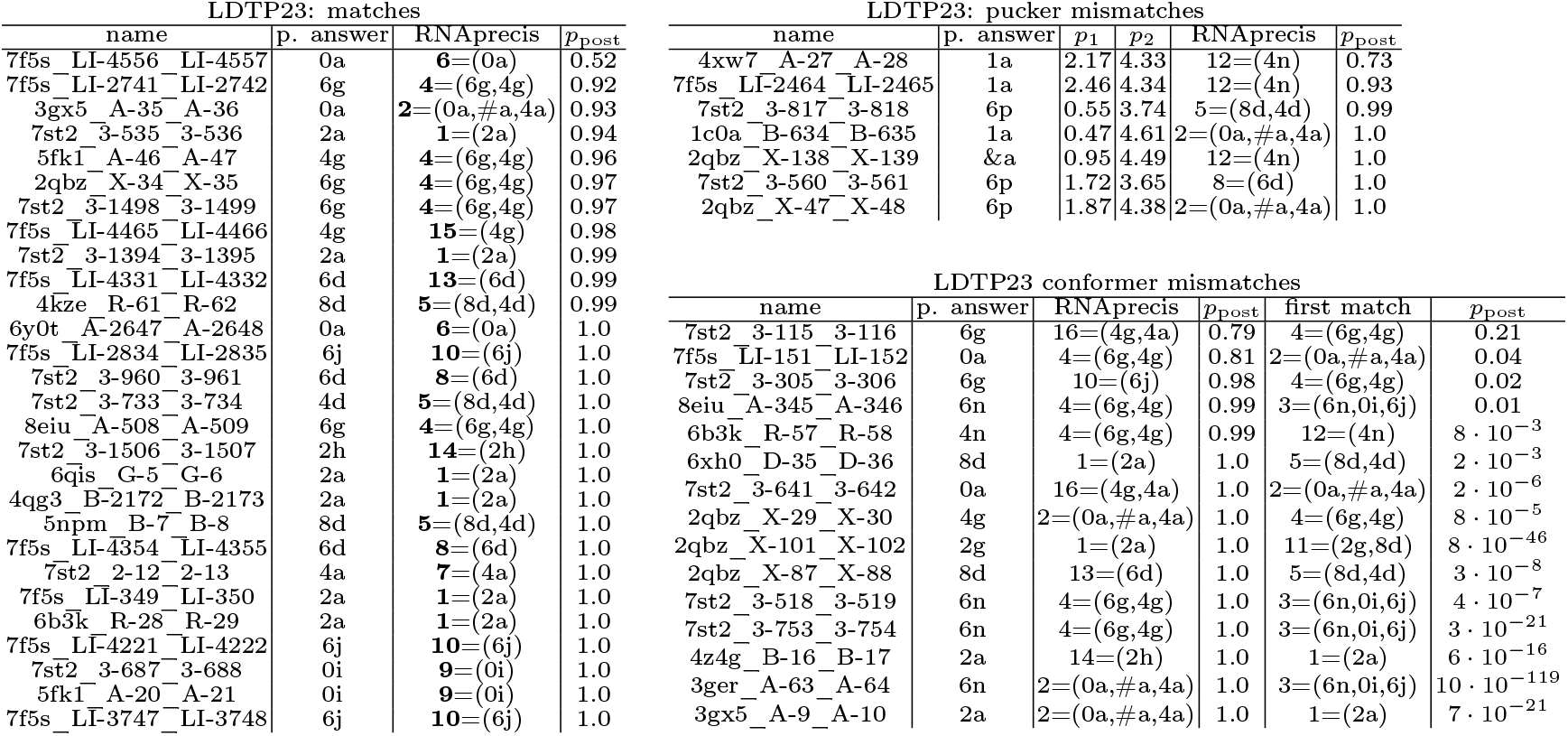
Suites for which the RNAprecis predicted cluster matches the sequence related conformer (blue in the plots), the sugar pucker-pair determined by the Pperp criterion do not match the sugar pucker-pair of the sequence related conformers (red in the plots), or the RNAprecis predicted cluster does not match the sequence related conformer (orange in the plots); each sorted by posterior probability.

| LDTP23: matches |  |  |  | LDTP23: pucker mismatches |  |  |  |  |  |
| --- | --- | --- | --- | --- | --- | --- | --- | --- | --- |
| name | p. answer | RNAprecis | $p_{\text{post}}$ | name | p. answer | $p_1$ | $p_2$ | RNAprecis | $p_{\text{post}}$ |
| 7f5s_LI-4556_LI-4557 | 0a | 6=(0a) | 0.52 | 4xw7_A-27_A-28 | 1a | 2.17 | 4.33 | 12=(4n) | 0.73 |
| 7f5s_LI-2741_LI-2742 | 6g | 4=(6g,4g) | 0.92 | 7f5s_LI-2464_LI-2465 | 1a | 2.46 | 4.34 | 12=(4n) | 0.93 |
| 3gx5_A-35_A-36 | 0a | 2=(0a,#a,4a) | 0.93 | 7st2_3-817_3-818 | 6p | 0.55 | 3.74 | 5=(8d,4d) | 0.99 |
| 7st2_3-535_3-536 | 2a | 1=(2a) | 0.94 | 1c0a_B-634_B-635 | 1a | 0.47 | 4.61 | 2=(0a,#a,4a) | 1.0 |
| 5fk1_A-46_A-47 | 4g | 4=(6g,4g) | 0.96 | 2qbz_X-138_X-139 | &a | 0.95 | 4.49 | 12=(4n) | 1.0 |
| 2qbz_X-34_X-35 | 6g | 4=(6g,4g) | 0.97 | 7st2_3-560_3-561 | 6p | 1.72 | 3.65 | 8=(6d) | 1.0 |
| 7st2_3-1498_3-1499 | 6g | 4=(6g,4g) | 0.97 | 2qbz_X-47_X-48 | 6p | 1.87 | 4.38 | 2=(0a,#a,4a) | 1.0 |
| 7f5s_LI-4465_LI-4466 | 4g | 15=(4g) | 0.98 |  |  |  |  |  |  |
| 7st2_3-1394_3-1395 | 2a | 1=(2a) | 0.99 |  |  |  |  |  |  |
| 7f5s_LI-4331_LI-4332 | 6d | 13=(6d) | 0.99 |  |  |  |  |  |  |
| 4kze_R-61_R-62 | 8d | 5=(8d,4d) | 0.99 |  |  |  |  |  |  |
| 6y0t_A-2647_A-2648 | 0a | 6=(0a) | 1.0 |  |  |  |  |  |  |
| 7f5s_LI-2834_LI-2835 | 6j | 10=(6j) | 1.0 |  |  |  |  |  |  |
| 7st2_3-960_3-961 | 6d | 8=(6d) | 1.0 |  |  |  |  |  |  |
| 7st2_3-733_3-734 | 4d | 5=(8d,4d) | 1.0 |  |  |  |  |  |  |
| 8eiu_A-508_A-509 | 6g | 4=(6g,4g) | 1.0 |  |  |  |  |  |  |
| 7st2_3-1506_3-1507 | 2h | 14=(2h) | 1.0 |  |  |  |  |  |  |
| 6qis_G-5_G-6 | 2a | 1=(2a) | 1.0 |  |  |  |  |  |  |
| 4qg3_B-2172_B-2173 | 2a | 1=(2a) | 1.0 |  |  |  |  |  |  |
| 5npm_B-7_B-8 | 8d | 5=(8d,4d) | 1.0 |  |  |  |  |  |  |
| 7f5s_LI-4354_LI-4355 | 6d | 8=(6d) | 1.0 |  |  |  |  |  |  |
| 7st2_2-12_2-13 | 4a | 7=(4a) | 1.0 |  |  |  |  |  |  |
| 7f5s_LI-349_LI-350 | 2a | 1=(2a) | 1.0 |  |  |  |  |  |  |
| 6b3k_R-28_R-29 | 2a | 1=(2a) | 1.0 |  |  |  |  |  |  |
| 7f5s_LI-4221_LI-4222 | 6j | 10=(6j) | 1.0 |  |  |  |  |  |  |
| 7st2_3-687_3-688 | 0i | 9=(0i) | 1.0 |  |  |  |  |  |  |
| 5fk1_A-20_A-21 | 0i | 9=(0i) | 1.0 |  |  |  |  |  |  |
| 7f5s_LI-3747_LI-3748 | 6j | 10=(6j) | 1.0 |  |  |  |  |  |  |

Table S8: Suites for which the RNAprecis predicted cluster matches the sequence related conformer (blue in the plots), the sugar pucker-pair determined by the Pperp criterion do not match the sugar pucker-pair of the sequence related conformers (red in the plots), or the RNAprecis predicted cluster ...
| LDTP23 conformer mismatches |  |  |  |  |  |
| --- | --- | --- | --- | --- | --- |
| name | p. answer | RNAprecis | $p_{\text{post}}$ | first match | $p_{\text{post}}$ |
| 7st2_3-115_3-116 | 6g | 16=(4g,4a) | 0.79 | 4=(6g,4g) | 0.21 |
| 7f5s_LI-151_LI-152 | 0a | 4=(6g,4g) | 0.81 | 2=(0a,#a,4a) | 0.04 |
| 7st2_3-305_3-306 | 6g | 10=(6j) | 0.98 | 4=(6g,4g) | 0.02 |
| 8eiu_A-345_A-346 | 6n | 4=(6g,4g) | 0.99 | 3=(6n,0i,6j) | 0.01 |
| 6b3k_R-57_R-58 | 4n | 4=(6g,4g) | 0.99 | 12=(4n) | $8 \cdot 10^{-3}$ |
| 6xh0_D-35_D-36 | 8d | 1=(2a) | 1.0 | 5=(8d,4d) | $2 \cdot 10^{-3}$ |
| 7st2_3-641_3-642 | 0a | 16=(4g,4a) | 1.0 | 2=(0a,#a,4a) | $2 \cdot 10^{-6}$ |
| 2qbz_X-29_X-30 | 4g | 2=(0a,#a,4a) | 1.0 | 4=(6g,4g) | $8 \cdot 10^{-5}$ |
| 2qbz_X-101_X-102 | 2g | 1=(2a) | 1.0 | 11=(2g,8d) | $8 \cdot 10^{-46}$ |
| 2qbz_X-87_X-88 | 8d | 13=(6d) | 1.0 | 5=(8d,4d) | $3 \cdot 10^{-8}$ |
| 7st2_3-518_3-519 | 6n | 4=(6g,4g) | 1.0 | 3=(6n,0i,6j) | $4 \cdot 10^{-7}$ |
| 7st2_3-753_3-754 | 6n | 4=(6g,4g) | 1.0 | 3=(6n,0i,6j) | $3 \cdot 10^{-21}$ |
| 4z4g_B-16_B-17 | 2a | 14=(2h) | 1.0 | 1=(2a) | $6 \cdot 10^{-16}$ |
| 3ger_A-63_A-64 | 6n | 2=(0a,#a,4a) | 1.0 | 3=(6n,0i,6j) | $10 \cdot 10^{-119}$ |
| 3gx5_A-9_A-10 | 2a | 2=(0a,#a,4a) | 1.0 | 1=(2a) | $7 \cdot 10^{-21}$ |

**Figure S20:**
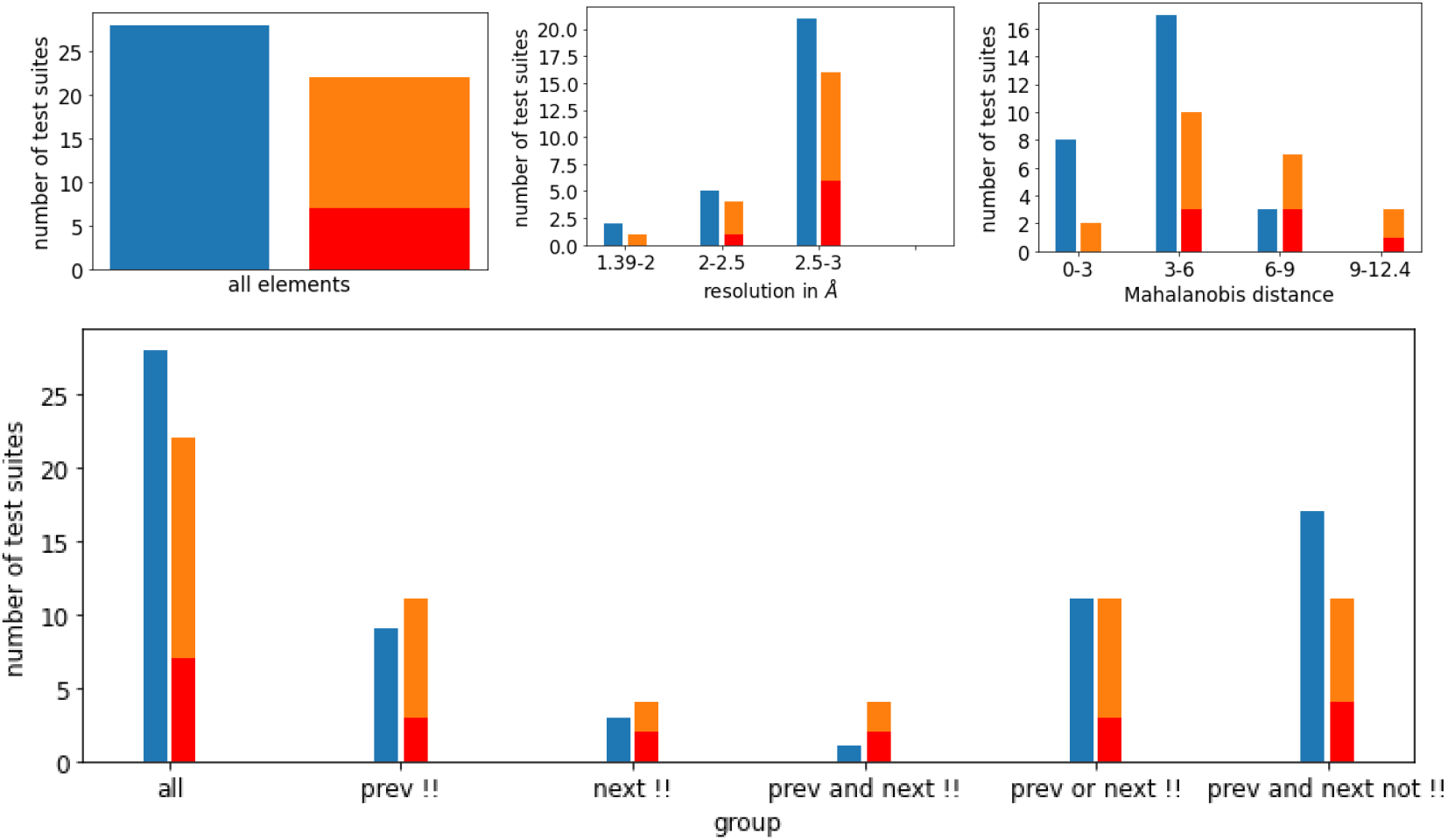
Relative distribution of agreements and disagreements of MINT-AGE cluster prediction with sequence related probable conformer classes for the test set LDT23 with color scheme (reflecting blue for match, orange for mismatch and red for pucker-pair mismatch) from Section 3.4 in the main text). Top left: overall histogram. Top middle: histogram reflecting varying resolutions of PDB suites. Top right: histogram reflecting varying Mahanalobis distances from the predicted cluster. One can see that with increasing distance the rate of mismatches increases. This indicates that despite its limitations discussed in Remark 2.3 in the main text the distance is useful as an empirical indicator of prediction confidence. Bottom: histogram also reflecting dependence on whether previous, next or both suites have been !! or not.

**Figure S21:**
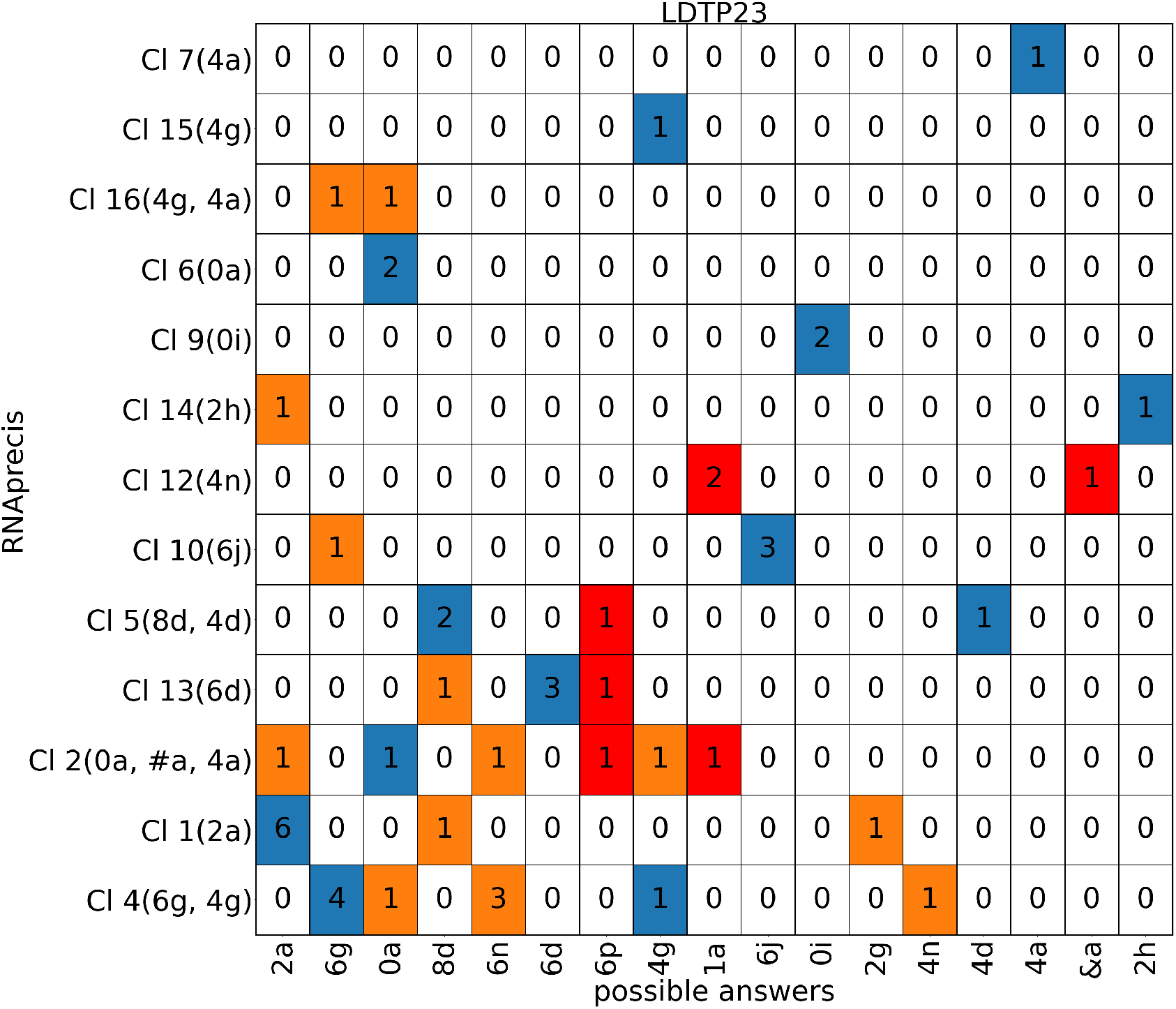
Classification matrix listing counts of probable sequence related conformation classes (horizontal axis) versus MINT-AGE clusters (vertical, with conformer classes in parentheses from MINT-AGE training, see Figure S11) for the low detail training data set of sugar pucker-pair LDP23 with colors (from Section 3.4 in the main text): blue for match, orange for mismatch and red for pucker-pair mismatch. Columns listing multiple suite conformers are from sites solved in different conformations across sequence-related models, and any of the sequence-related conformations were accepted as a matching predictions. Columns listing multiple suite conformers are from sites solved in different conformations across sequence-related models, and any of the sequence-related conformations were accepted as a matching predictions.

#### E.4 Results for the test data set LDTP22

**Table S9:**
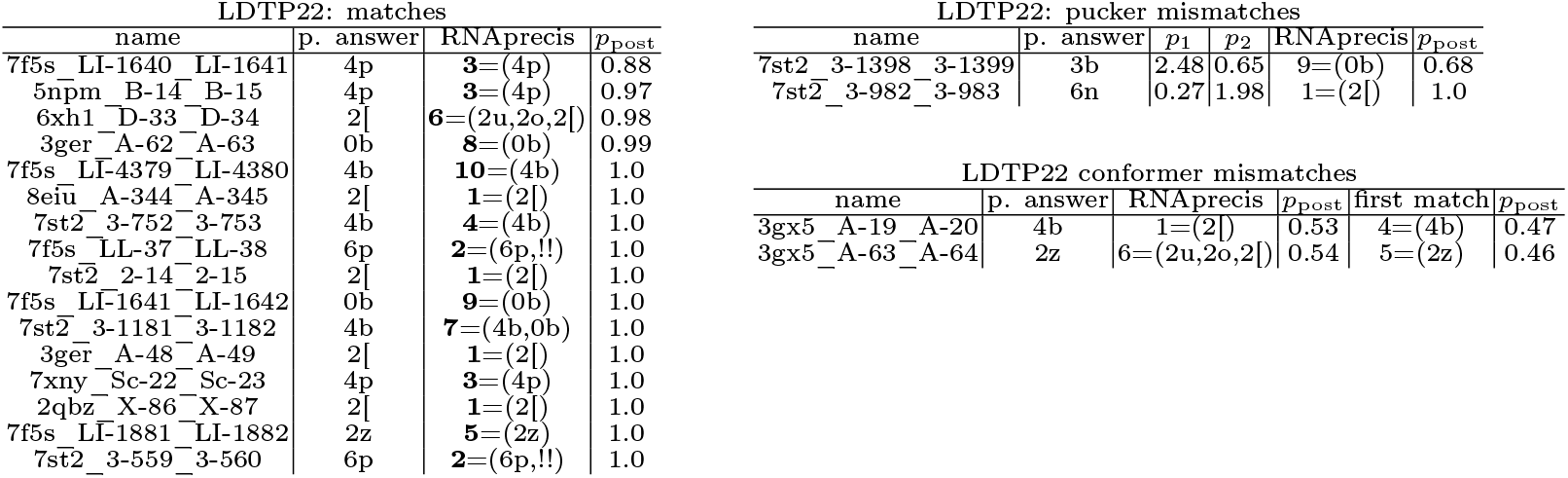
Suites for which the RNAprecis predicted cluster matches the sequence related conformer (blue in the plots), the sugar pucker-pair determined by the Pperp criterion do not match the sugar pucker-pair of the sequence related conformers (red in the plots), or the RNAprecis predicted cluster does not match the sequence related conformer (orange in the plots); each sorted by posterior probability.

**Figure S22:**
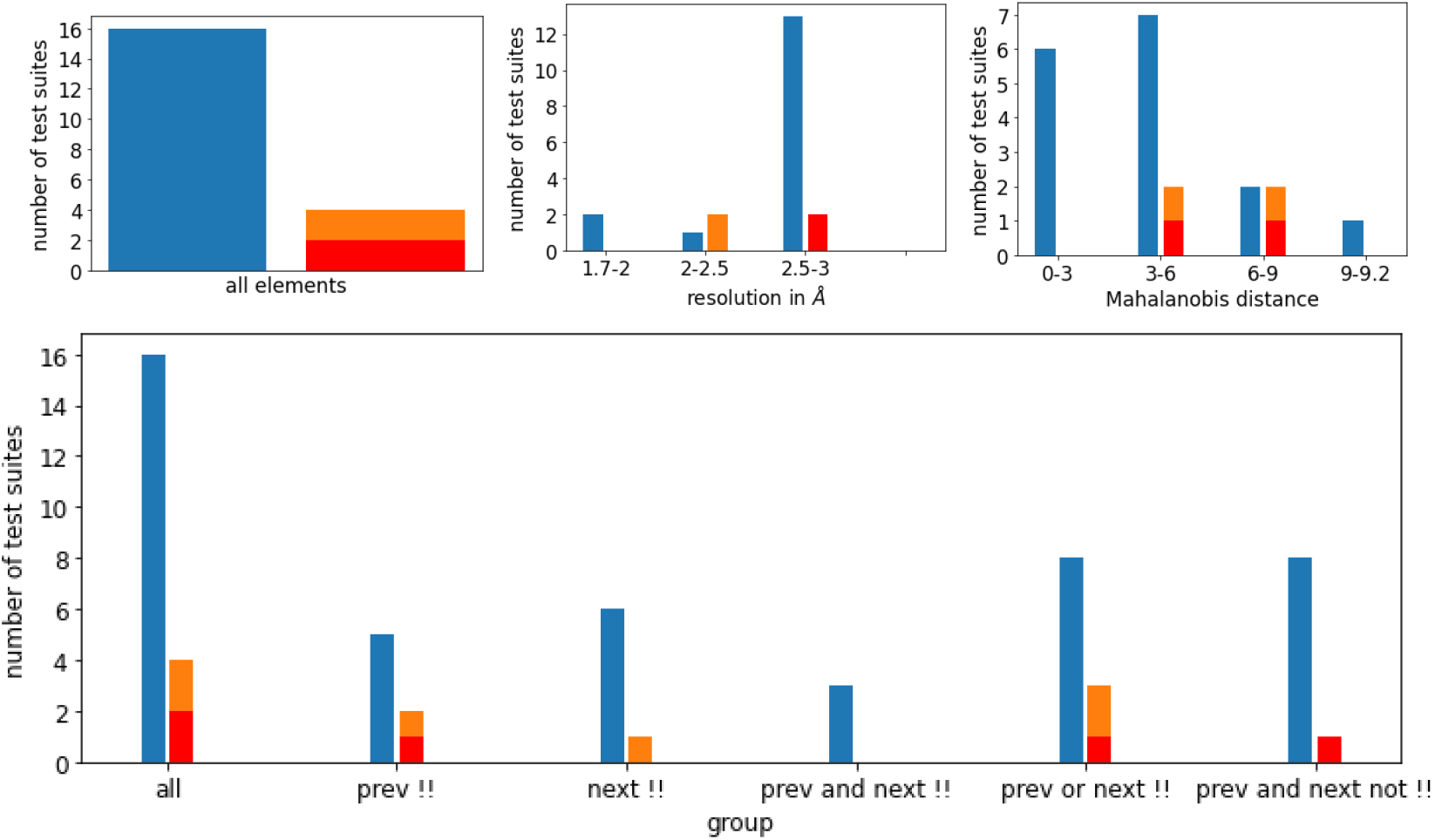
Relative distribution of agreements and disagreements of MINT-AGE cluster prediction with sequence related probable conformer classes for the test set LDT22 with color scheme (reflecting blue for match, orange for mismatch and red for pucker-pair mismatch) from Section 3.4 in the main text). Top left: overall histogram. Top middle: histogram reflecting varying resolutions of PDB suites. Top right: histogram reflecting varying Mahanalobis distances from the predicted cluster. One can see that with increasing distance the rate of mismatches increases. This indicates that despite its limitations discussed in Remark 2.3 in the main text the distance is useful as an empirical indicator of prediction confidence. Bottom: histogram also reflecting dependence on whether previous, next or both suites have been !! or not.

**Figure S23:**
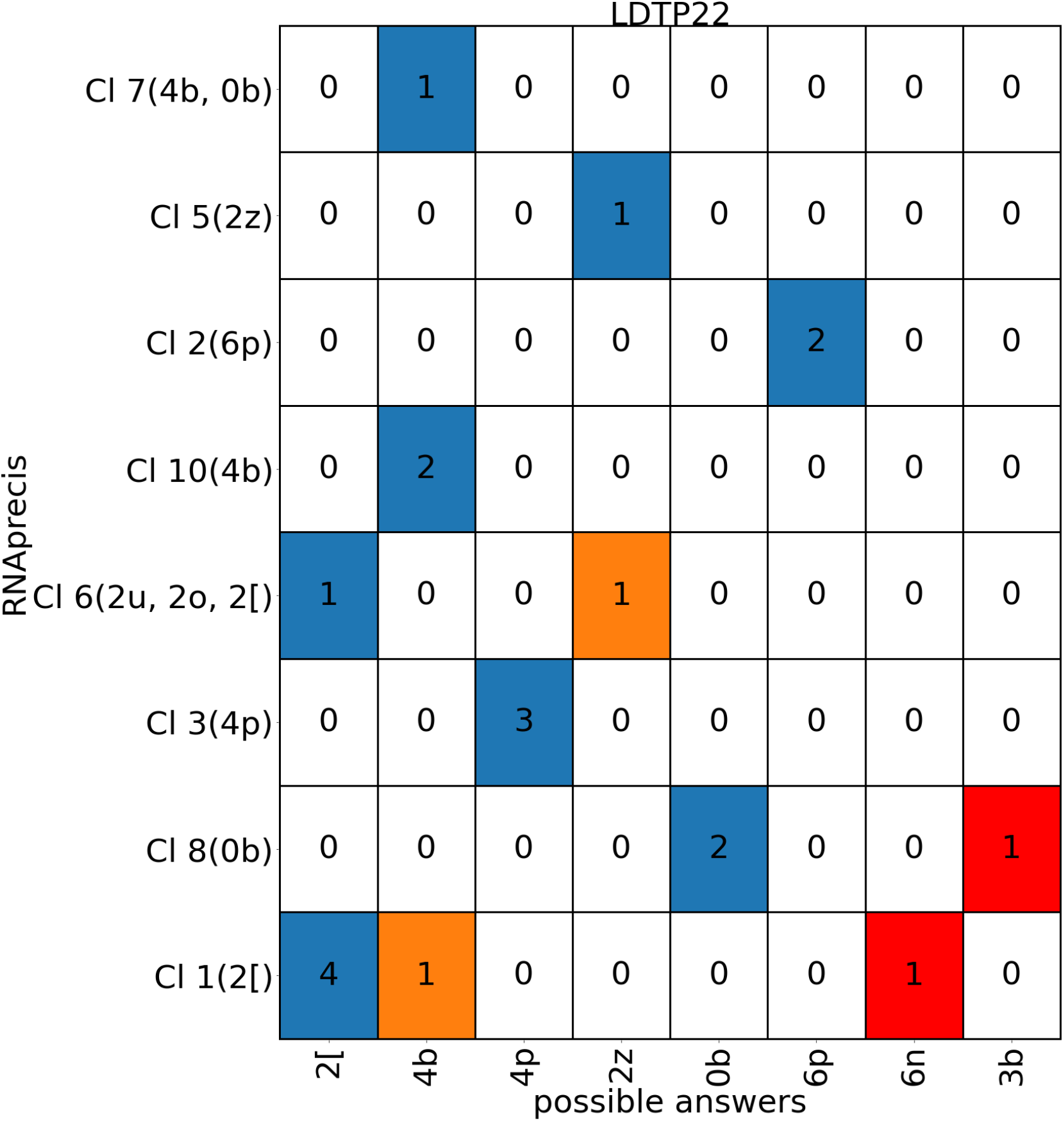
Classification matrix listing counts of probable sequence related conformation classes (horizontal axis) versus MINT-AGE clusters (vertical, with conformer classes in parentheses from MINT-AGE training, see Figure S12) for the low detail training data set of sugar pucker-pair LDP22 with colors (from Section 3.4 in the main text): blue for match, orange for mismatch and red for pucker-pair mismatch. Columns listing multiple suite conformers are from sites solved in different conformations across sequence-related models, and any of the sequence-related conformations were accepted as a matching predictions. Columns listing multiple suite conformers are from sites solved in different conformations across sequence-related models, and any of the sequence-related conformations were accepted as a matching predictions.

#### E.5 Results for the test data set 8b0xP33

**Table S10:**
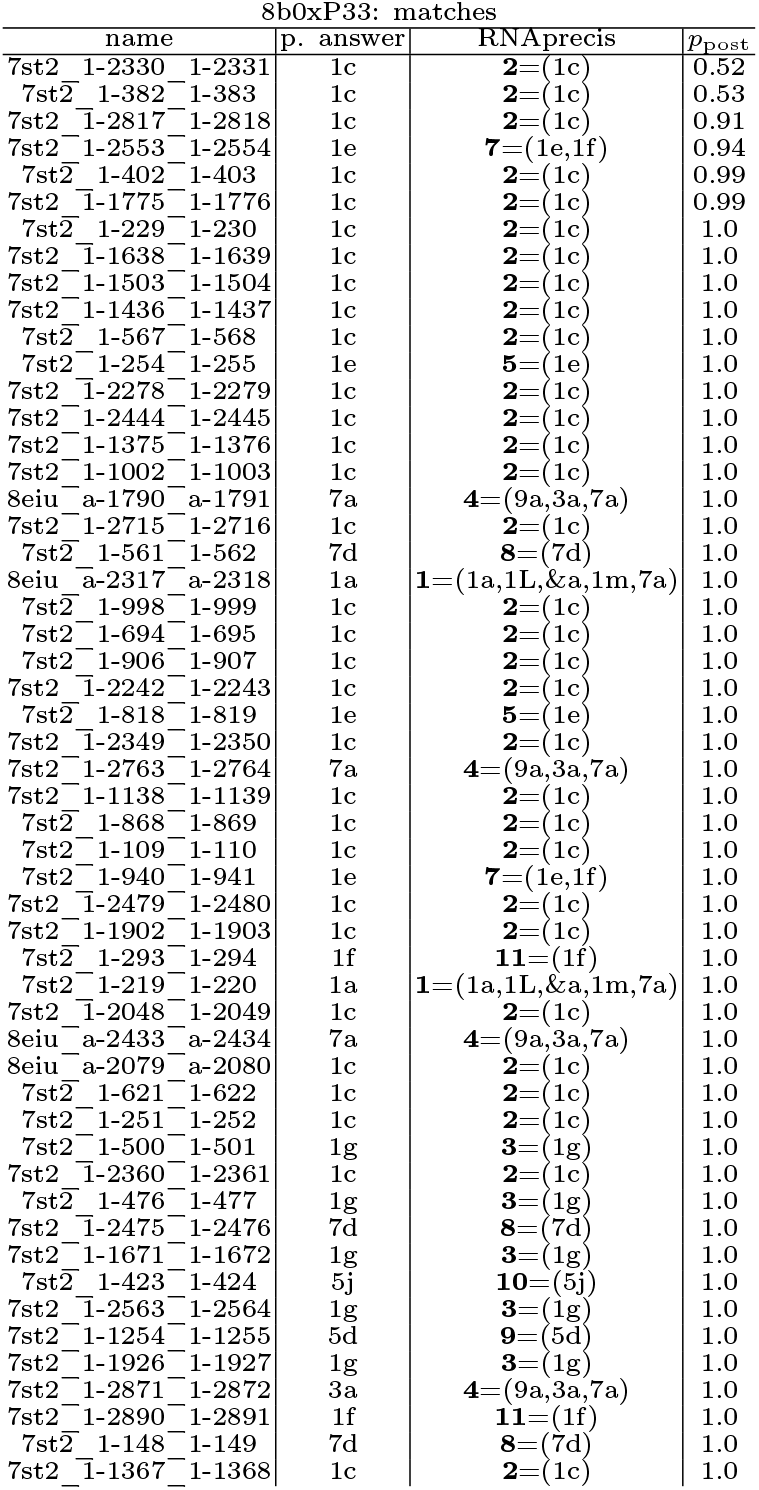
Suites for which the RNAprecis predicted cluster matches the sequence related conformer (blue in the plots) sorted by posterior probability.

**Table S11:** Suites for which the sugar pucker-pair determined by the Pperp criterion do not match the sugar pucker-pair of the sequence related conformers (red in the plots) or the RNAprecis predicted cluster does not match the sequence related conformer (orange in the plots) sorted by posterior probability.

| 8b0xP33: pucker mismatches |  |  |  |  |  |  |  |
| --- | --- | --- | --- | --- | --- | --- | --- |
| name | | | p. answer | $p_1$ | $p_2$ | RNAprecis | $p_{\text{post}}$ |
| 7st2 | 1-1418 | 1-1419 | 1b | 4.78 | 3.78 | 5=(1e) | 1.0 |
| 7st2 | 1-2807 | 1-2808 | 1b | 4.57 | 3.36 | 2=(1c) | 1.0 |
| 7st2 | 1-395 | 1-396 | 4g | 3.89 | 4.7 | 1=(1a,1L,&a,1m,7a) | 1.0 |
| 7st2 | 1-394 | 1-395 | 1b | 4.74 | 3.89 | 2=(1c) | 1.0 |
| 7st2 | 1-228 | 1-229 | 6n | 4.05 | 4.57 | 4=(9a,3a,7a) | 1.0 |
| 7st2 | 1-857 | 1-858 | 1b | 4.63 | 2.96 | 2=(1c) | 1.0 |
| 7st2 | 1-1457 | 1-1458 | 3b | 4.64 | 3.17 | 4=(9a,3a,7a) | 1.0 |
| 7st2 | 1-1559 | 1-1560 | 2a | 3.93 | 4.61 | 4=(9a,3a,7a) | 1.0 |
| 7st2 | 1-1626 | 1-1627 | 0a | 3.86 | 4.67 | 4=(9a,3a,7a) | 1.0 |
| 7st2 | 1-1856 | 1-1857 | 1b | 4.64 | 4.12 | 2=(1c) | 1.0 |
| 7st2 | 1-1883 | 1-1884 | 1b | 4.44 | 3.99 | 2=(1c) | 1.0 |
| 7st2 | 1-1900 | 1-1901 | 6j | 3.63 | 4.5 | 10=(5j) | 1.0 |
| 7st2 | 1-2808 | 1-2809 | 0a | 3.36 | 4.61 | 4=(9a,3a,7a) | 1.0 |

| 8b0xP33: conformer mismatches |  |  |  |  |  |  |
| --- | --- | --- | --- | --- | --- | --- |
| name | p. answer | RNAprecis | $p_{\text{post}}$ | first match | $p_{\text{post}}$ | |
| 7st2_1-95_1-96 | 1c | 7=(1e,1f) | 0.64 | 2=(1c) | 0.36 |  |
| 8eiu_a-1749_a-1750 | 1a | 2=(1c) | 0.68 | 1=(1a,1L,&a,1m,7a) | 0.32 |  |
| 7st2_1-333_1-334 | 1a | 2=(1c) | 0.77 | 1=(1a,1L,&a,1m,7a) | 0.23 |  |
| 7st2_1-57_1-58 | 1c | 1=(1a,1L,&a,1m,7a) | 0.79 | 2=(1c) | 0.21 |  |
| 8eiu_a-1240_a-1241 | &a | 5=(1e) | 0.92 | 1=(1a,1L,&a,1m,7a) | $2 \cdot 10^{-28}$ | |
| 8eiu_a-2534_a-2535 | 1f | 2=(1c) | 0.96 | 7=(1e,1f) | 0.04 |  |
| 7st2_1-1477_1-1478 | 1a | 2=(1c) | 0.98 | 1=(1a,1L,&a,1m,7a) | 0.02 |  |
| 8eiu_a-44_a-45 | 1f | 2=(1c) | 0.98 | 7=(1e,1f) | 0.02 |  |
| 7st2_1-29_1-30 | 1a | 2=(1c) | 0.98 | 1=(1a,1L,&a,1m,7a) | 0.02 |  |
| 7st2_1-2326_1-2327 | 1a | 6=(1f) | 0.99 | 1=(1a,1L,&a,1m,7a) | $4 \cdot 10^{-9}$ | |
| 7st2_1-2473_1-2474 | 1a | 2=(1c) | 1.0 | 1=(1a,1L,&a,1m,7a) | $2 \cdot 10^{-3}$ | |
| 7st2_1-516_1-517 | 1a | 2=(1c) | 1.0 | 1=(1a,1L,&a,1m,7a) | $1 \cdot 10^{-3}$ | |
| 7st2_1-2886_1-2887 | 1a | 2=(1c) | 1.0 | 1=(1a,1L,&a,1m,7a) | $3 \cdot 10^{-4}$ | |
| 8eiu_a-1873_a-1874 | 1a | 2=(1c) | 1.0 | 1=(1a,1L,&a,1m,7a) | $1 \cdot 10^{-4}$ | |
| 8eiu_a-537_a-538 | 7a | 5=(1e) | 1.0 | 4=(9a,3a,7a) | $10 \cdot 10^{-6}$ | |
| 7st2_1-1154_1-1155 | 1L | 4=(9a,3a,7a) | 1.0 | 1=(1a,1L,&a,1m,7a) | $2 \cdot 10^{-8}$ | |
| 8eiu_a-130_a-131 | 1e | 2=(1c) | 1.0 | 7=(1e,1f) | $2 \cdot 10^{-8}$ | |
| 7st2_1-2564_1-2565 | 1a | 2=(1c) | 1.0 | 1=(1a,1L,&a,1m,7a) | $4 \cdot 10^{-9}$ | |
| 7st2_1-499_1-500 | 1a | 2=(1c) | 1.0 | 1=(1a,1L,&a,1m,7a) | $2 \cdot 10^{-9}$ | |
| 8eiu_a-2474_a-2475 | 1a | 2=(1c) | 1.0 | 1=(1a,1L,&a,1m,7a) | $1 \cdot 10^{-9}$ | |
| 7st2_1-2822_1-2823 | 1g | 8=(7d) | 1.0 | 3=(1g) | $6 \cdot 10^{-10}$ | |
| 7st2_1-633_1-634 | 1a | 2=(1c) | 1.0 | 1=(1a,1L,&a,1m,7a) | $7 \cdot 10^{-12}$ | |
| 7st2_1-2858_1-2859 | 1a | 2=(1c) | 1.0 | 1=(1a,1L,&a,1m,7a) | $10 \cdot 10^{-13}$ | |
| 7st2_1-1185_1-1186 | 3d | 3=(1g) | 1.0 | - | - |  |
| 7st2_1-2202_1-2203 | 3d | 3=(1g) | 1.0 | - | - |  |
| 7st2_1-464_1-465 | 1a | 2=(1c) | 1.0 | 1=(1a,1L,&a,1m,7a) | $8 \cdot 10^{-14}$ | |
| 7st2_1-45_1-46 | 5n | 4=(9a,3a,7a) | 1.0 | - | - |  |
| 7st2_1-1004_1-1005 | 3g | 3=(1g) | 1.0 | - | - |  |
| 7st2_1-1756_1-1757 | 3d | 8=(7d) | 1.0 | - | - |  |
| 7st2_1-1358_1-1359 | &a | 4=(9a,3a,7a) | 1.0 | 1=(1a,1L,&a,1m,7a) | $4 \cdot 10^{-21}$ | |
| 7st2_1-218_1-219 | 1a | 2=(1c) | 1.0 | 1=(1a,1L,&a,1m,7a) | $4 \cdot 10^{-18}$ | |
| 7st2_1-466_1-467 | 1a | 2=(1c) | 1.0 | 1=(1a,1L,&a,1m,7a) | $1 \cdot 10^{-19}$ | |

**Figure S24:**
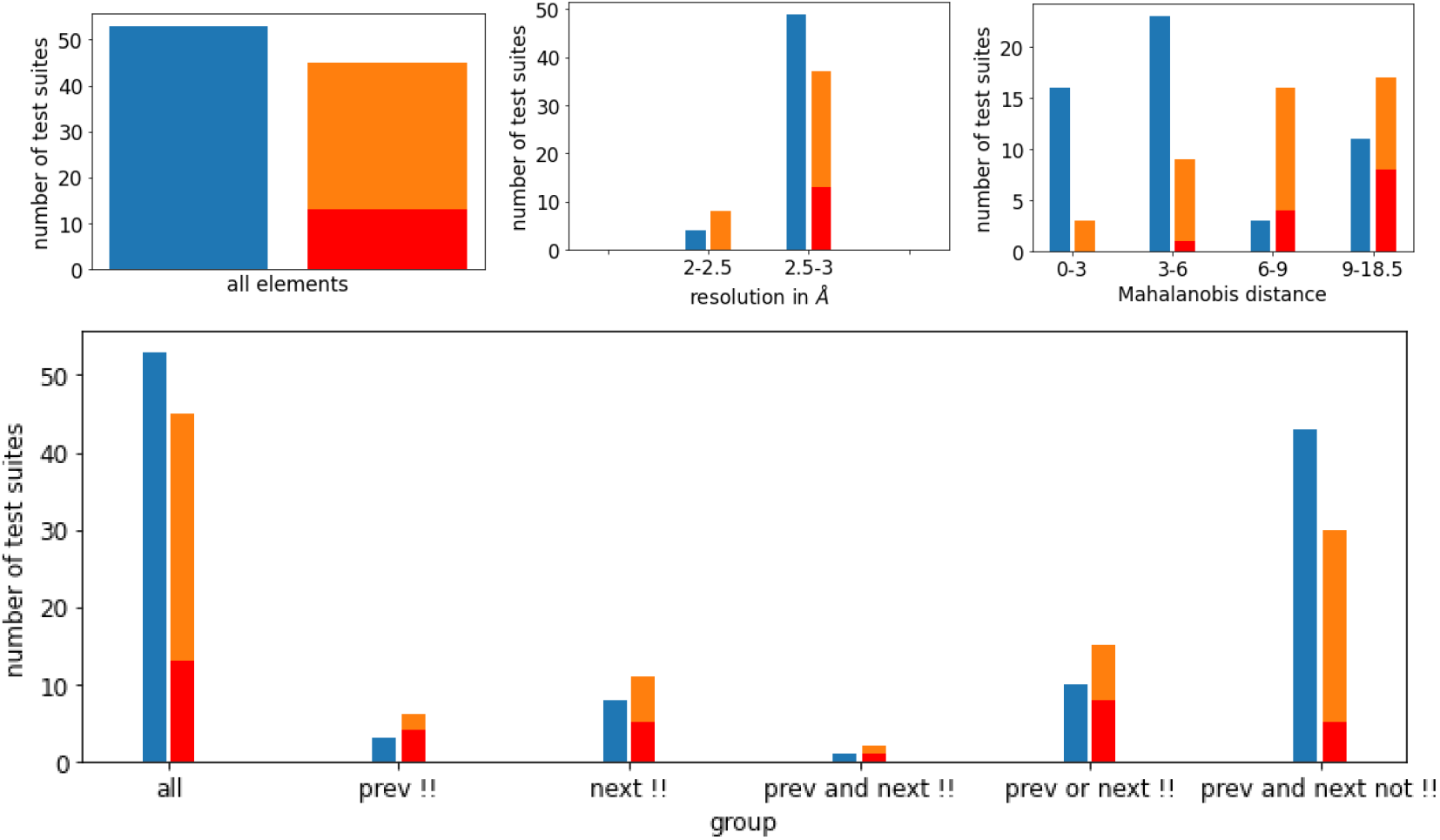
Relative distribution of agreements and disagreements of MINT-AGE cluster prediction with sequence related probable conformer classes for the test set LDTP33 with color scheme (reflecting blue for match, orange for mismatch and red for pucker-pair mismatch) from Section 3.4 in the main text). Top left: overall histogram. Top middle: histogram reflecting varying resolutions of PDB suites. Top right: histogram reflecting varying Mahanalobis distances from the predicted cluster. One can see that with increasing distance the rate of mismatches increases. This indicates that despite its limitations discussed in Remark 2.3 in the main text the distance is useful as an empirical indicator of prediction confidence. Bottom: histogram also reflecting dependence on whether previous, next or both suites have been !! or not. While false predictions are more likely if at least one neighboring suite is a !! suite, roughly half of the total false predictions are from suites without a neighboring !! suite.

**Figure S25:**
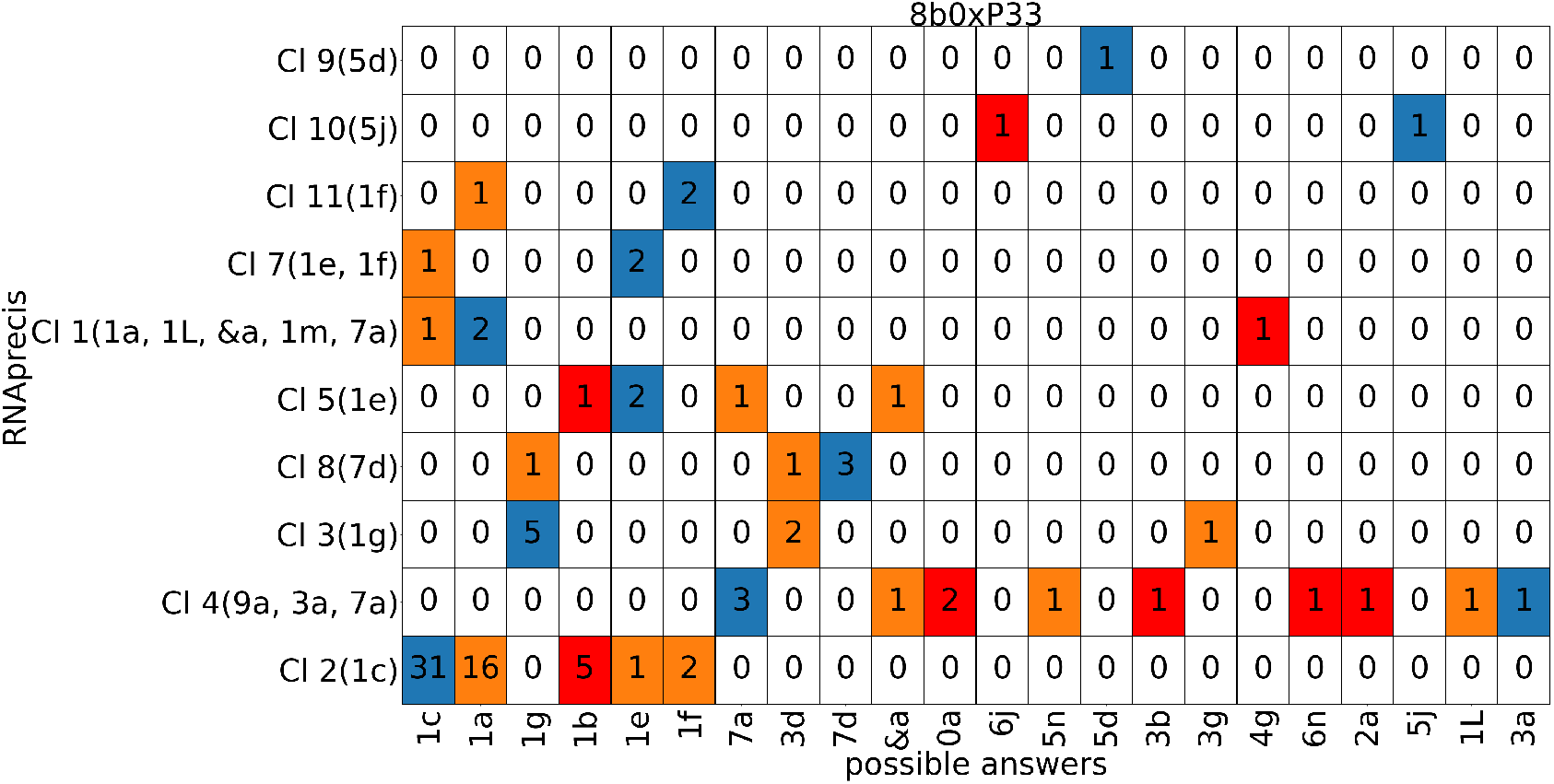
Classification matrix listing counts of probable sequence related conformation classes (horizontal axis) versus MINT-AGE clusters (vertical, with conformer classes in parentheses from MINT-AGE training, see Figure S9) for the low detail training data set of sugar pucker-pair LDP33 with colors (from Section 3.4 in the main text): blue for match, orange for mismatch and red for pucker-pair mismatch. Columns listing multiple suite conformers are from sites solved in different conformations across sequence-related models, and any of the sequence-related conformations were accepted as a matching predictions. Columns listing multiple suite conformers are from sites solved in different conformations across sequence-related models, and any of the sequence-related conformations were accepted as a matching predictions.

#### E.6 Results for the test data set 8b0xP32

**Table S12:**
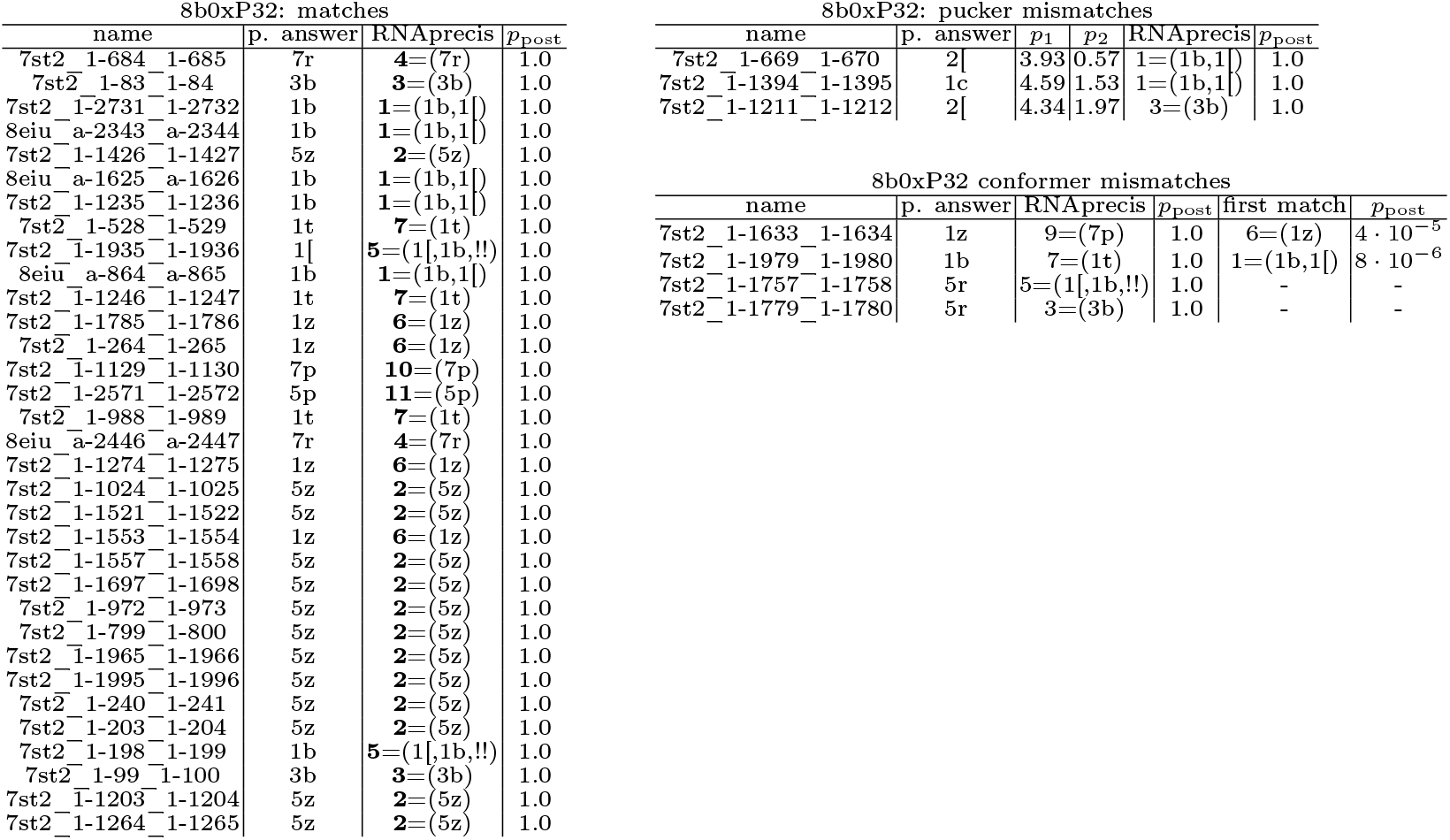
Suites for which the RNAprecis predicted cluster matches the sequence related conformer (blue in the plots), the sugar pucker-pair determined by the Pperp criterion do not match the sugar pucker-pair of the sequence related conformers (red in the plots), or the RNAprecis predicted cluster does not match the sequence related conformer (orange in the plots); each sorted by posterior probability.

**Figure S26:**
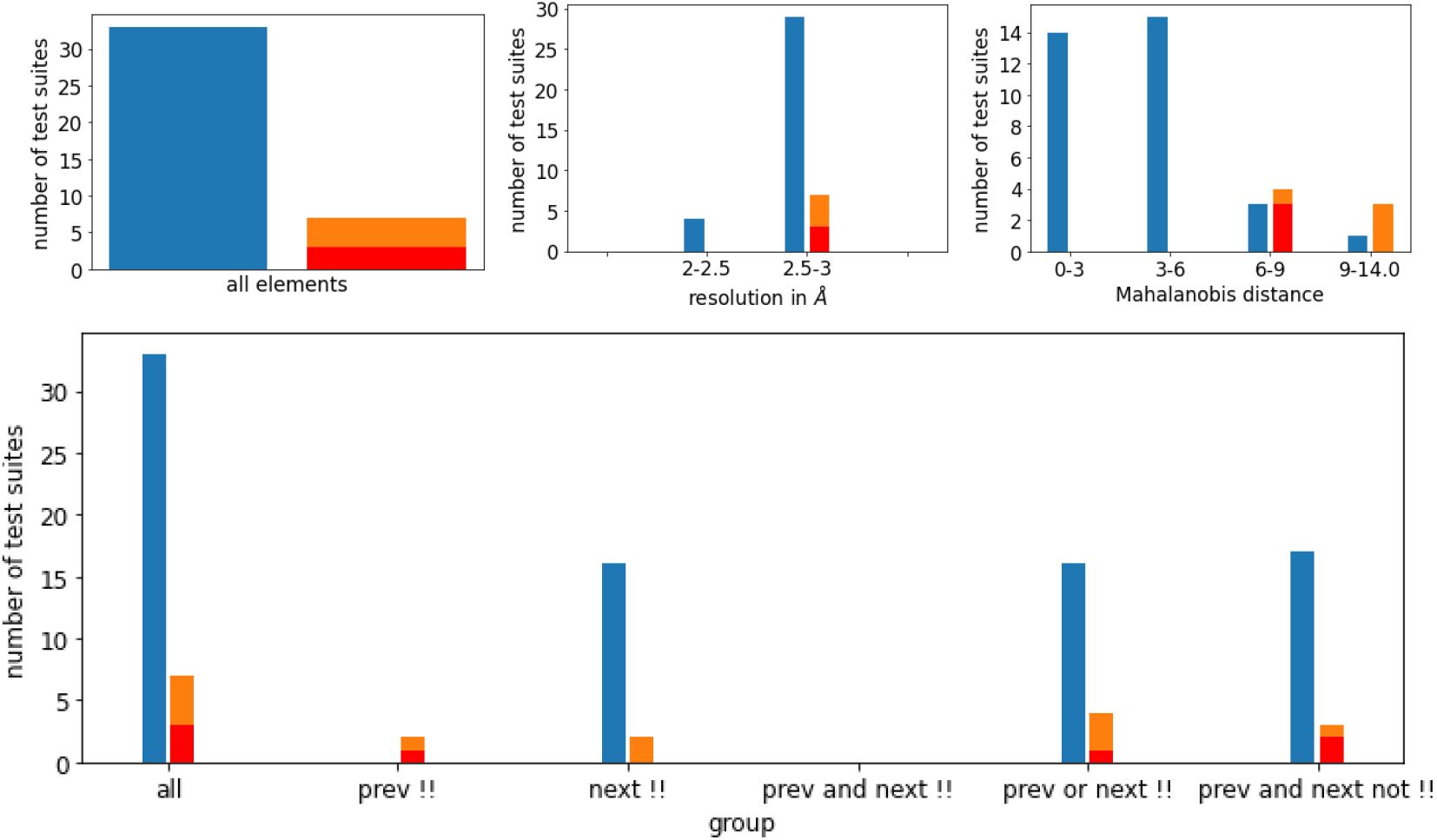
Relative distribution of agreements and disagreements of MINT-AGE cluster prediction with sequence related probable conformer classes for the test set LDT32 with color scheme (reflecting blue for match, orange for mismatch and red for pucker-pair mismatch) from Section 3.4 in the main text). Top left: overall histogram. Top middle: histogram reflecting varying resolutions of PDB suites. Top right: histogram reflecting varying Mahanalobis distances from the predicted cluster. One can see that with increasing distance the rate of mismatches increases. This indicates that despite its limitations discussed in Remark 2.3 in the main text the distance is useful as an empirical indicator of prediction confidence. Bottom: histogram also reflecting dependence on whether previous, next or both suites have been !! or not.

**Figure S27:**
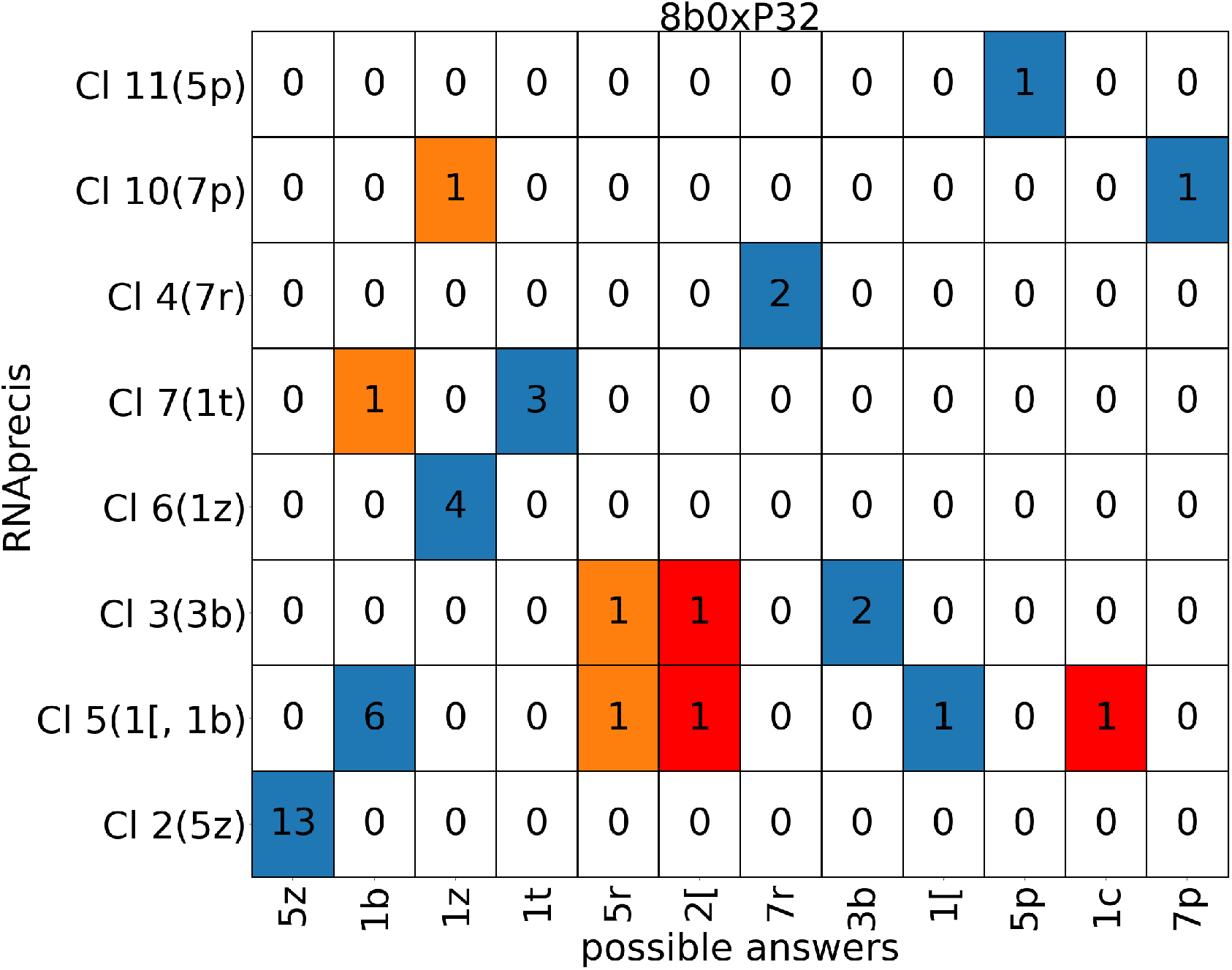
Classification matrix listing counts of probable sequence related conformation classes (horizontal axis) versus MINT-AGE clusters (vertical, with conformer classes in parentheses from MINT-AGE training, see Figure S10) for the low detail training data set of sugar pucker-pair LDP32 with colors (from Section 3.4 in the main text): blue for match, orange for mismatch and red for pucker-pair mismatch. Columns listing multiple suite conformers are from sites solved in different conformations across sequence-related models, and any of the sequence-related conformations were accepted as a matching predictions. As pointed out in Section 3.4, the **1a**,**3g**,**8d** and **3d**,**7p** suites have multiple sequence-related conformations. This means that the counts in these two columns are inflated.

#### E.7 Results for the test data set 8b0xP23

**Table S13:**
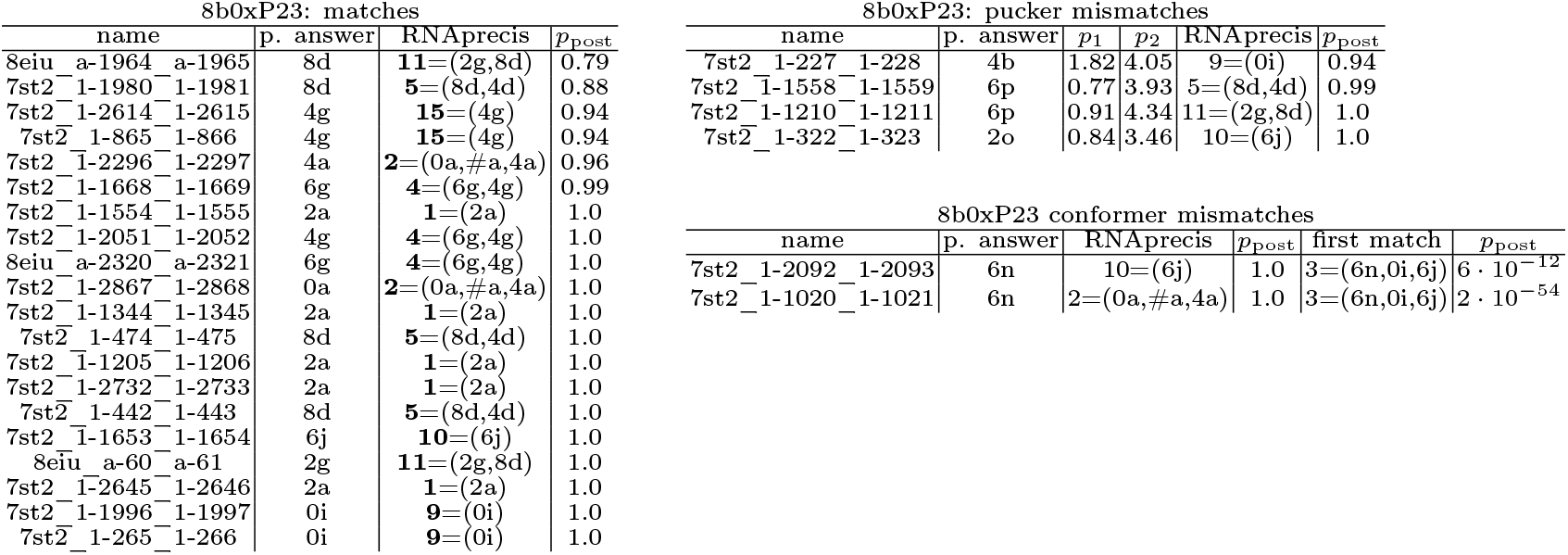
Suites for which the RNAprecis predicted cluster matches the sequence related conformer (blue in the plots), the sugar pucker-pair determined by the Pperp criterion do not match the sugar pucker-pair of the sequence related conformers (red in the plots), or the RNAprecis predicted cluster does not match the sequence related conformer (orange in the plots); each sorted by posterior probability.

| 8b0xP23: matches |  |  |  | 8b0xP23: pucker mismatches |  |  |  |  |  |
| --- | --- | --- | --- | --- | --- | --- | --- | --- | --- |
| name | p. answer | RNAprecis | $p_{\text{post}}$ | name | p. answer | $p_1$ | $p_2$ | RNAprecis | $p_{\text{post}}$ |
| 8eiu_a-1964_a-1965 | 8d | **11**=(2g,8d) | 0.79 | 7st2_1-227_1-228 | 4b | 1.82 | 4.05 | 9=(0i) | 0.94 |
| 7st2_1-1980_1-1981 | 8d | **5**=(8d,4d) | 0.88 | 7st2_1-1558_1-1559 | 6p | 0.77 | 3.93 | 5=(8d,4d) | 0.99 |
| 7st2_1-2614_1-2615 | 4g | **15**=(4g) | 0.94 | 7st2_1-1210_1-1211 | 6p | 0.91 | 4.34 | 11=(2g,8d) | 1.0 |
| 7st2_1-865_1-866 | 4g | **15**=(4g) | 0.94 | 7st2_1-322_1-323 | 2o | 0.84 | 3.46 | 10=(6j) | 1.0 |
| 7st2_1-2296_1-2297 | 4a | **2**=(0a,#a,4a) | 0.96 |  |  |  |  |  |  |
| 7st2_1-1668_1-1669 | 6g | **4**=(6g,4g) | 0.99 |  |  |  |  |  |  |
| 7st2_1-1554_1-1555 | 2a | **1**=(2a) | 1.0 |  |  |  |  |  |  |
| 7st2_1-2051_1-2052 | 4g | **4**=(6g,4g) | 1.0 |  |  |  |  |  |  |
| 8eiu_a-2320_a-2321 | 6g | **4**=(6g,4g) | 1.0 |  |  |  |  |  |  |
| 7st2_1-2867_1-2868 | 0a | **2**=(0a,#a,4a) | 1.0 |  |  |  |  |  |  |
| 7st2_1-1344_1-1345 | 2a | **1**=(2a) | 1.0 |  |  |  |  |  |  |
| 7st2_1-474_1-475 | 8d | **5**=(8d,4d) | 1.0 |  |  |  |  |  |  |
| 7st2_1-1205_1-1206 | 2a | **1**=(2a) | 1.0 |  |  |  |  |  |  |
| 7st2_1-2732_1-2733 | 2a | **1**=(2a) | 1.0 |  |  |  |  |  |  |
| 7st2_1-442_1-443 | 8d | **5**=(8d,4d) | 1.0 |  |  |  |  |  |  |
| 7st2_1-1653_1-1654 | 6j | **10**=(6j) | 1.0 |  |  |  |  |  |  |
| 8eiu_a-60_a-61 | 2g | **11**=(2g,8d) | 1.0 |  |  |  |  |  |  |
| 7st2_1-2645_1-2646 | 2a | **1**=(2a) | 1.0 |  |  |  |  |  |  |
| 7st2_1-1996_1-1997 | 0i | **9**=(0i) | 1.0 |  |  |  |  |  |  |
| 7st2_1-265_1-266 | 0i | **9**=(0i) | 1.0 |  |  |  |  |  |  |

Table S13: Suites for which the RNAprecis predicted cluster matches the sequence related conformer (blue in the plots), the sugar pucker-pair determined by the Pperp criterion do not match the sugar pucker-pair of the sequence related conformers (red in the plots), or the RNAprecis predicted ...
| 8b0xP23 conformer mismatches |  |  |  |  |  |
| --- | --- | --- | --- | --- | --- |
| name | p. answer | RNAprecis | $p_{\text{post}}$ | first match | $p_{\text{post}}$ |
| 7st2_1-2092_1-2093 | 6n | 10=(6j) | 1.0 | 3=(6n,0i,6j) | $6 \cdot 10^{-12}$ |
| 7st2_1-1020_1-1021 | 6n | 2=(0a,#a,4a) | 1.0 | 3=(6n,0i,6j) | $2 \cdot 10^{-54}$ |

**Figure S28:**
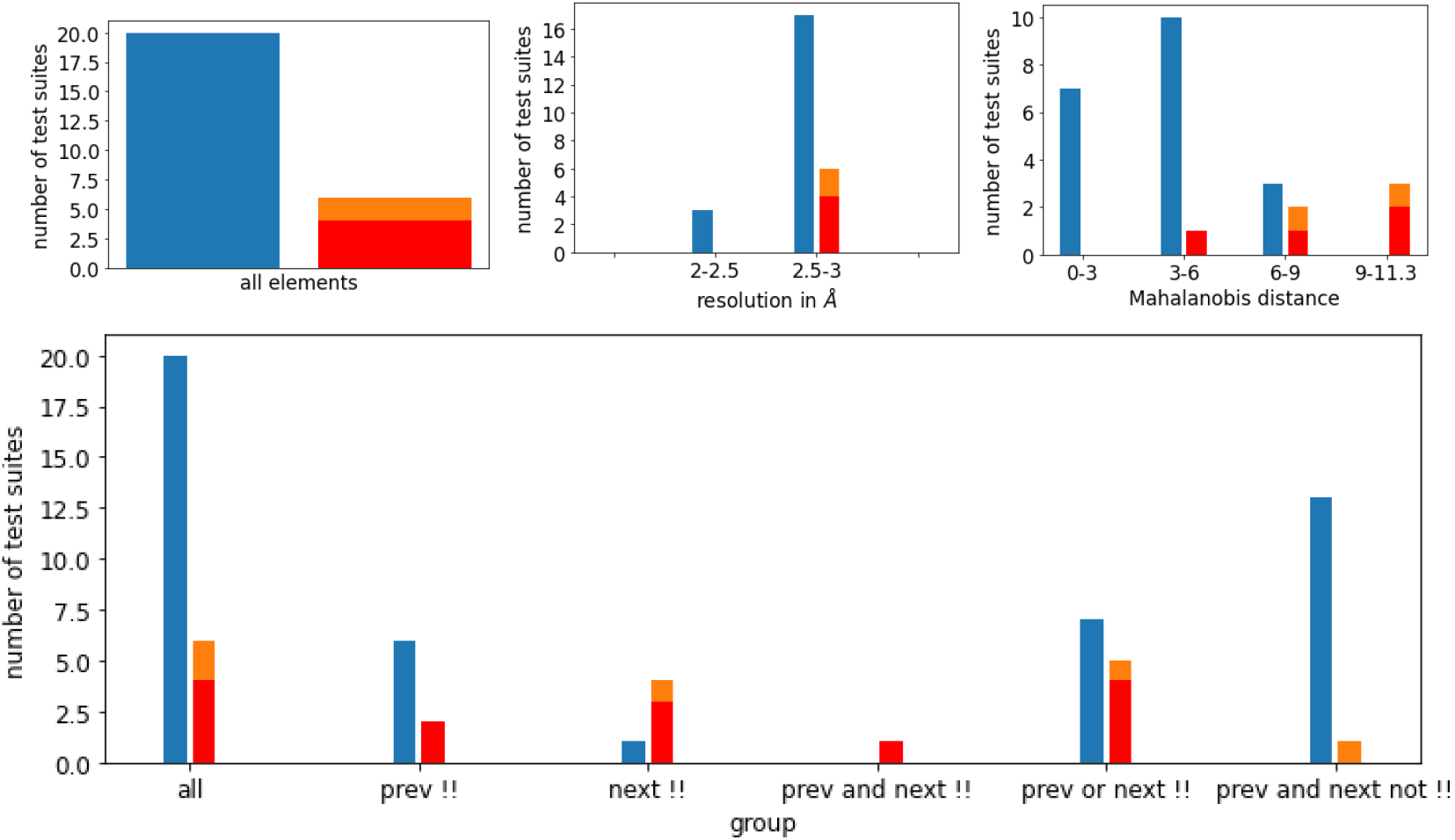
Relative distribution of agreements and disagreements of MINT-AGE cluster prediction with sequence related probable conformer classes for the test set LDT23 with color scheme (reflecting blue for match, orange for mismatch and red for pucker-pair mismatch) from Section 3.4 in the main text). Top left: overall histogram. Top middle: histogram reflecting varying resolutions of PDB suites. Top right: histogram reflecting varying Mahanalobis distances from the predicted cluster. One can see that with increasing distance the rate of mismatches increases. This indicates that despite its limitations discussed in Remark 2.3 in the main text the distance is useful as an empirical indicator of prediction confidence. Bottom: histogram also reflecting dependence on whether previous, next or both suites have been !! or not.

**Figure S29:**
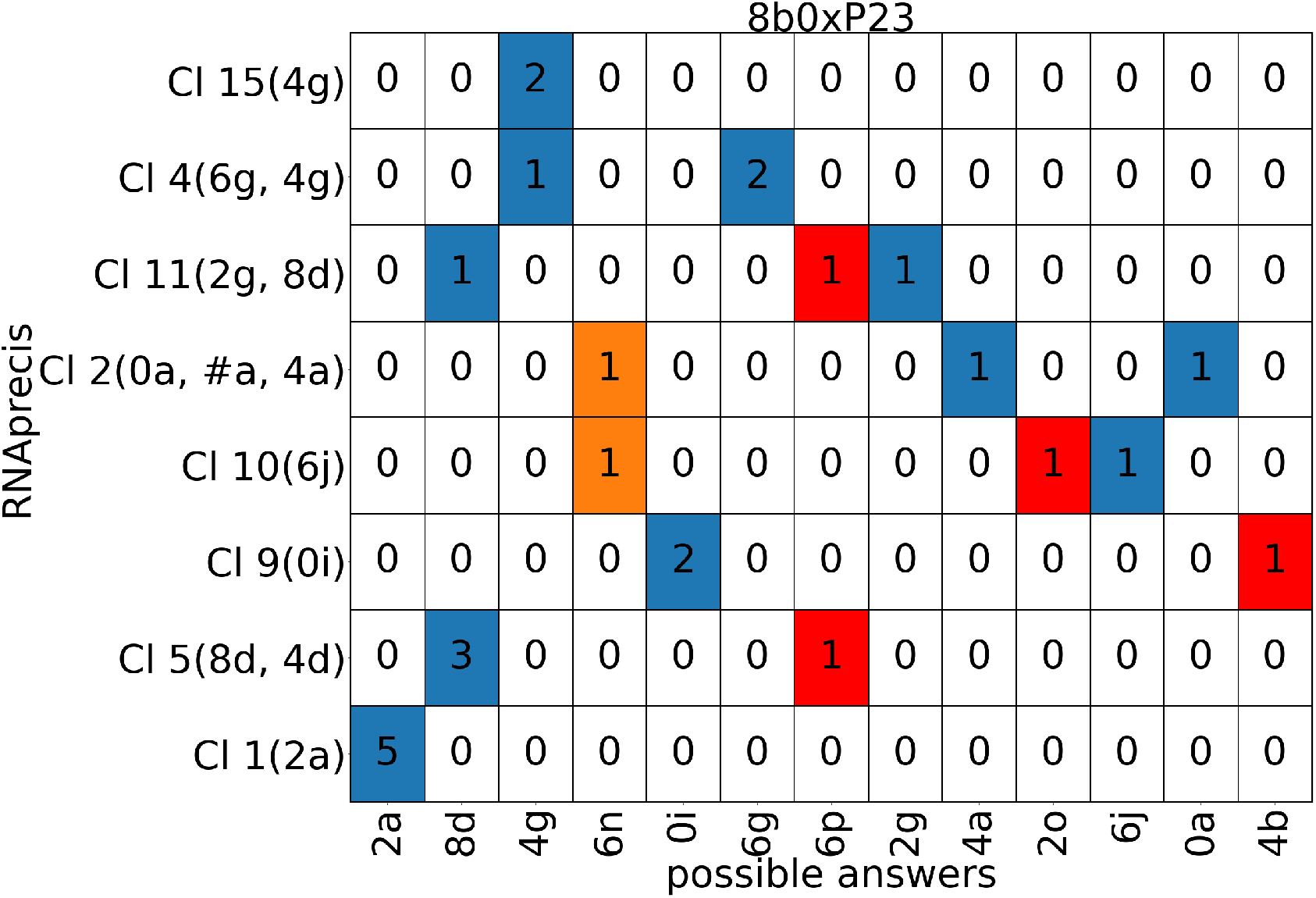
Classification matrix listing counts of probable sequence related conformation classes (horizontal axis) versus MINT-AGE clusters (vertical, with conformer classes in parentheses from MINT-AGE training, see Figure S11) for the low detail training data set of sugar pucker-pair LDP23 with colors (from Section 3.4 in the main text): blue for match, orange for mismatch and red for pucker-pair mismatch. Columns listing multiple suite conformers are from sites solved in different conformations across sequence-related models, and any of the sequence-related conformations were accepted as a matching predictions. Columns listing multiple suite conformers are from sites solved in different conformations across sequence-related models, and any of the sequence-related conformations were accepted as a matching predictions.

#### E.8 Results for the test data set 8b0xP22

**Table S14:**
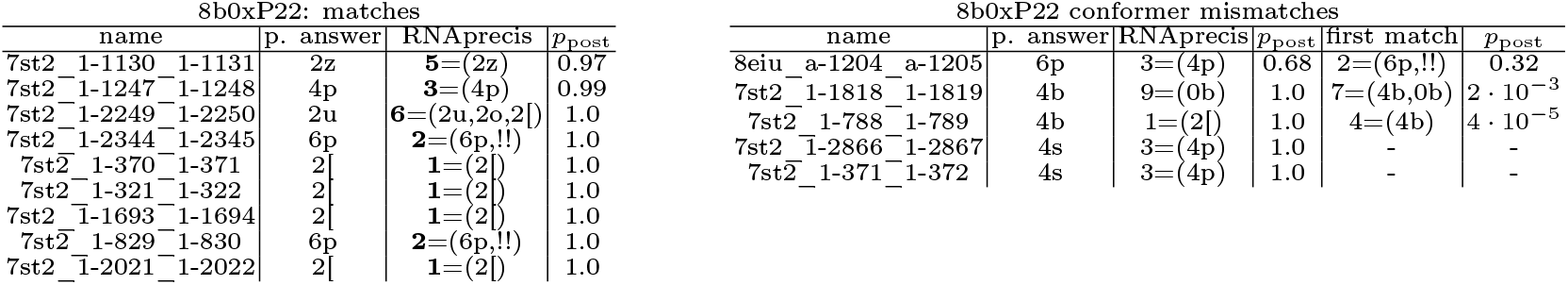
Suites for which the RNAprecis predicted cluster matches the sequence related conformer (blue in the plots), or the RNAprecis predicted cluster does not match the sequence related conformer (orange in the plots); each sorted by posterior probability. Since there are no cases where the sugar pucker-pair determined by the Pperp criterion do not match the sugar pucker-pair of the sequence related conformers (red in the plots) in this data set, this table is omitted.

**Figure S30:**
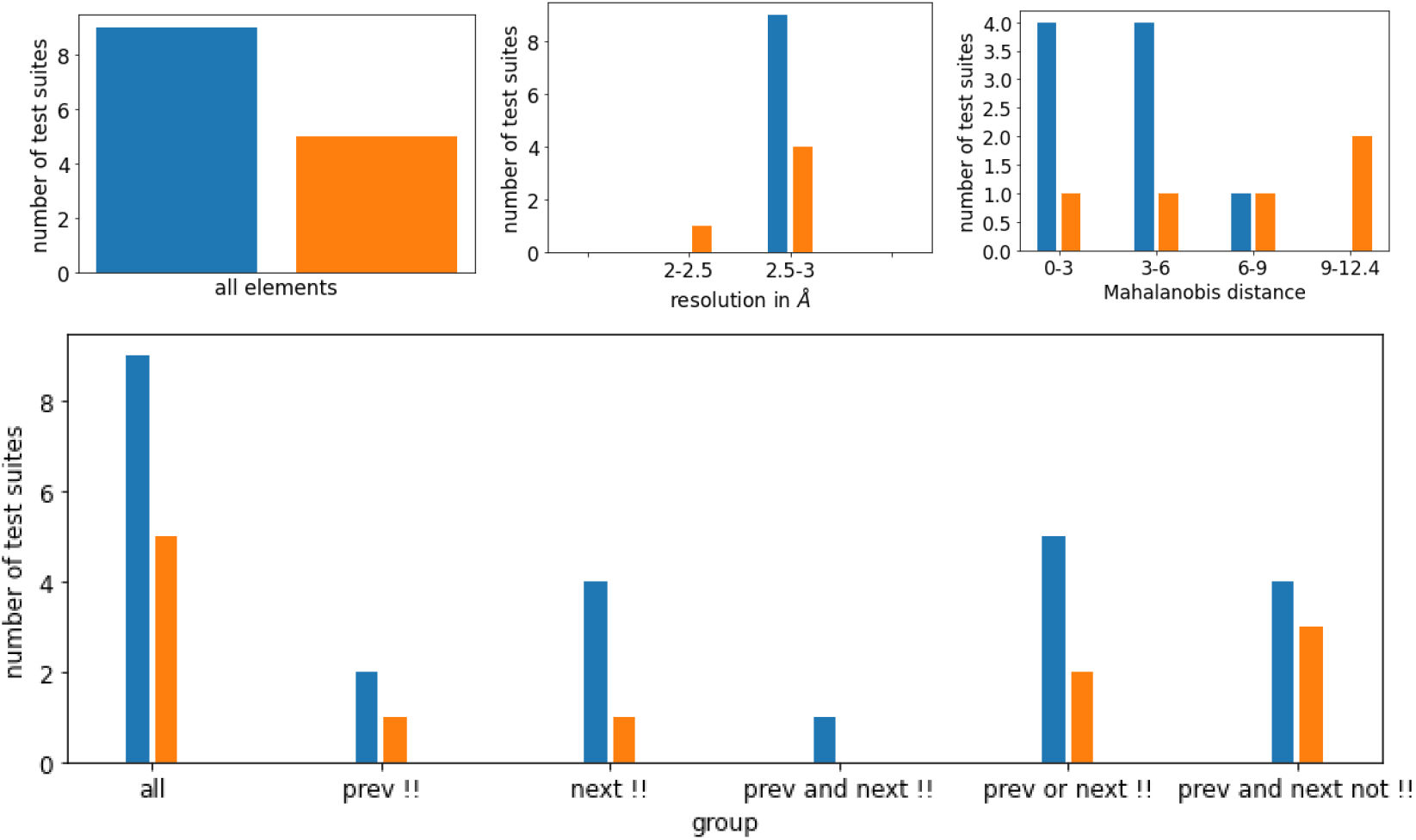
Relative distribution of agreements and disagreements of MINT-AGE cluster prediction with sequence related probable conformer classes for the test set LDT22 with color scheme (reflecting blue for match, orange for mismatch and red for pucker-pair mismatch) from Section 3.4 in the main text). Top left: overall histogram. Top middle: histogram reflecting varying resolutions of PDB suites. Top right: histogram reflecting varying Mahanalobis distances from the predicted cluster. One can see that with increasing distance the rate of mismatches increases. This indicates that despite its limitations discussed in Remark 2.3 in the main text the distance is useful as an empirical indicator of prediction confidence. Bottom: histogram also reflecting dependence on whether previous, next or both suites have been !! or not.

**Figure S31:**
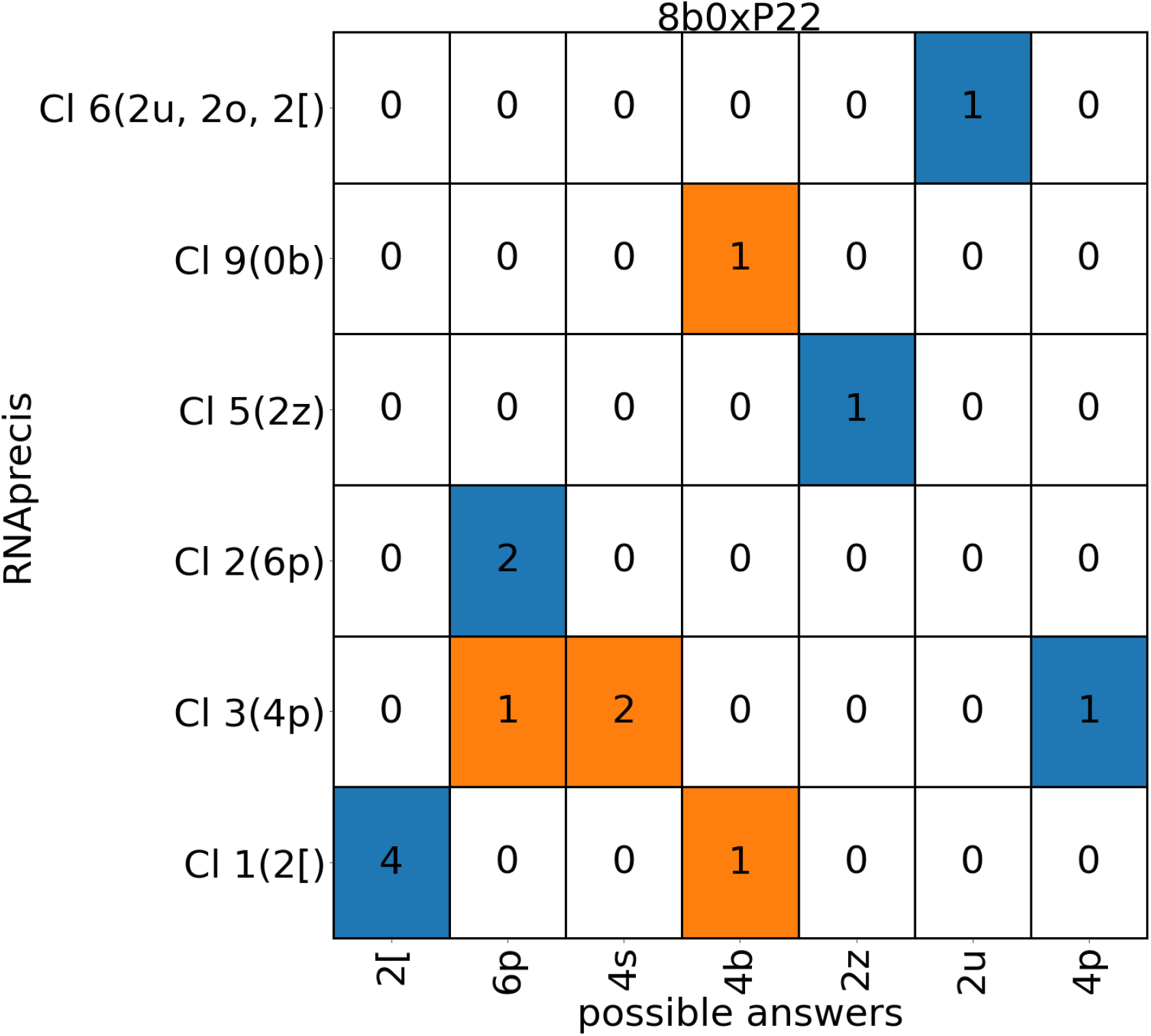
Classification matrix listing counts of probable sequence related conformation classes (horizontal axis) versus MINT-AGE clusters (vertical, with conformer classes in parentheses from MINT-AGE training, see Figure S12) for the low detail training data set of sugar pucker-pair LDP22 with colors (from Section 3.4 in the main text): blue for match, orange for mismatch and red for pucker-pair mismatch. Columns listing multiple suite conformers are from sites solved in different conformations across sequence-related models, and any of the sequence-related conformations were accepted as a matching predictions. Columns listing multiple suite conformers are from sites solved in different conformations across sequence-related models, and any of the sequence-related conformations were accepted as a matching predictions.

## Footnotes

1 http://rna.bgsu.edu/rna3dhub/nrlist

